# Illuminating the Ligandable Human Proteome with AI Protein Profiling

**DOI:** 10.1101/2025.09.07.670677

**Authors:** Guy W. Dayhoff, Daniel Kortzak, Mingzhe Shen, Ruibin Liu, Jianping Lin, Zhong-Yin Zhang, Jana Shen

## Abstract

Most human proteins lack chemical probes or pharmaceutical modulators, leaving much of the proteome unexplored. ^1,2^ Activity-based protein profiling has enabled proteome-scale discovery of protein-ligand interactions and covalent inhibitors, ^3–5^ but remains limited by probe chemistry, protein abundance, and discordant ligandability assignments across studies. ^6–9^ Machine-learning (ML) models can in principle generate proteomewide ligandability maps, but the predictive utility of current models is limited by the requirement for structures and the use of incomplete and weak training labels. ^9–12^ Here we developed an artificial intelligence protein profiling (AiPP), a sequence-based multitask platform built on the ESMC protein language model, ^13,14^ to accelerate proteomewide therapeutic discovery and target identification. While its primary task is identification of covalently ligandable cysteines, seven additional task heads and two external modules provide broader context by annotating reversible ligand-binding residues, disordered molecular recognition features, cysteine functional context, and reactivities. Central to the development is the LatentLift clustering approach, which leverages latent space similarities to reconcile conflicting experimental labels and facilitate model training. Applied to the human proteome, AiPP generated a cysteine-directed ligandability atlas that overcomes the limitations of current chemo-proteomic maps. As a proof of concept, we demonstrate that AiPP can guide the discovery of covalent allosteric inhibitors targeting the previously undruggable protein tyrosine phosphatase PTPN6. By linking sequence, ligandability and biological context, AiPP provides an open resource for therapeutic discovery, while LatentLift offers a general strategy for harmonizing proteomics data for ML applications.

---

The human proteome comprises ∼20,300 distinct proteins and modified variants, collectively amounting to at least 200,000 proteoforms. ^1^ Yet fewer than 900 proteins have been targeted by FDA-approved drugs. ^2^ This gap underscores both the need and the opportunity to expand the druggable proteome. Activity-based protein profiling (ABPP), which makes use of activity- or reactivity-based probes, fragment screening, and mass spectrometry (MS), profiles protein activity, reactivity, and ligand interactions in complex biological systems. ABPP has enabled proteome-scale discovery of reactive cysteines, ^3^ protein-ligand interactions, ^4^ and covalent inhibitors. ^5,15^ In particular, isoTOP-ABPP, ^3,16^ which quantifies competition between covalent probes and fragments, has enabled cysteine ligandability profiling across more than 10,000 proteins in diverse human cell lines. ^4,17–29^ However, a complete and accurate proteome-wide map of covalent ligandable sites remains out of reach. ABPP coverage is constrained by probe chemistry, protein abundance, and cell type ^5–8,30,31^ Even among quantified sites, ligandability assignments can vary across studies. ^7–9,11^ This variability arises largely from differences in probe and fragment identity, concentration, treatment time, sample context (for example, lysates versus live cells), and quantification protocol, ^7^ and to a lesser extent from differences in cellular state. ^8,11,32^ In addition, the shotgun nature of proteomics can preclude site-specific assignment for a subset of liganded cysteines. ^8,23,33^

Structural methods such as X-ray crystallography and cryo-EM can localize ligandable cysteines unambiguously, but only for a limited set of proteins. Databases derived from structures in the Protein Data Bank (PDB) report nearly 800 proteins with chemically modified cysteines, ^10,34,35^ and these data have enabled structure-based machine-learning (ML) models for predicting cysteine ligandability. ^10,12,36,37^ Most recently, ABPP data have been used to train structure-based models of cysteine reactivity and ligandability, including the tree-based CIAA, ^38^ and TopCySPAL, ^9^ and a convolutional neural network. ^11^ Of these, TopCySPAL is especially relevant as its training made use of both ABPP and PDB-derived data. ^9^ Yet three problems remain. First, no rigorous framework exists for learning ligandability from discordant experimental evidence, in which the same cysteine may be liganded, unliganded, or undetected depending on the study. Second, current models have limited external validation. Models trained on structure-derived labels have limited external testing, whereas models trained on ABPP data are rarely validated against orthogonal structural evidence. Third, existing models require protein structures, making them impractical or unreliable for structurally unresolved proteins and leaving a major coverage gap across the proteome.

Here we developed AiPP, a multitask AI protein profiling platform for proteome-wide ligandability prediction directly from protein sequence. Built on the protein language model ESMC, ^13,14^ AiPP uses LatentLift, a latent-space clustering approach, to reconcile heterogeneous chemoproteomic evidence. AiPP predicts covalently ligandable cysteines, reversible binding residues, and cysteine functional context, including disulfide formation and coordination with heme or metals. We used AiPP to extrapolate from limited, discordant experimental data and create an atlas of ligandable cysteine sites across the human proteome (AiPP Atlas 1.0). We present extensive benchmarking, case studies, and a proof-of-concept application to the discovery of covalent allosteric inhibitors targeting the undruggable phosphatase PTPN6.

## Limited proteome coverage and heterogeneous ligandability assignments

Based on 15 independent ABPP datasets published 2016–2025, ^4,8,17–29^ we constructed the LigCysABPP database, comprising 703,135 site-level records for 140,459 cysteines across 10,649 proteins (Supp. Fig. S1). These ABPP-quantified proteins cover 54% of the human cysteine-containing proteome, leaving the rest undetected (Fig. 1a and Supp. Fig. S1). Among the quantified proteins, 6,502 proteins (33% of the proteome) are liganded while 4,147 (21% of the proteome) are unliganded (Fig. 1a). Here, liganded refers to having at least one positive ABPP record, while unliganded refers to having only negative records. ABPP liganded 31% of known drug targets. The coverage is low for ion channels (3%) and GPCRs (7%, Fig. 1b), likely because most cysteines are not positioned near binding pockets. It is also noteworthy that among ABPP-quantified proteins, only 33% of cysteines are quantified (liganded or unliganded) whereas 67% are undetected (Fig. 1d). The limited proteome and drug target coverage by ABPP underscores the need for ML approaches.

**Figure 1.**
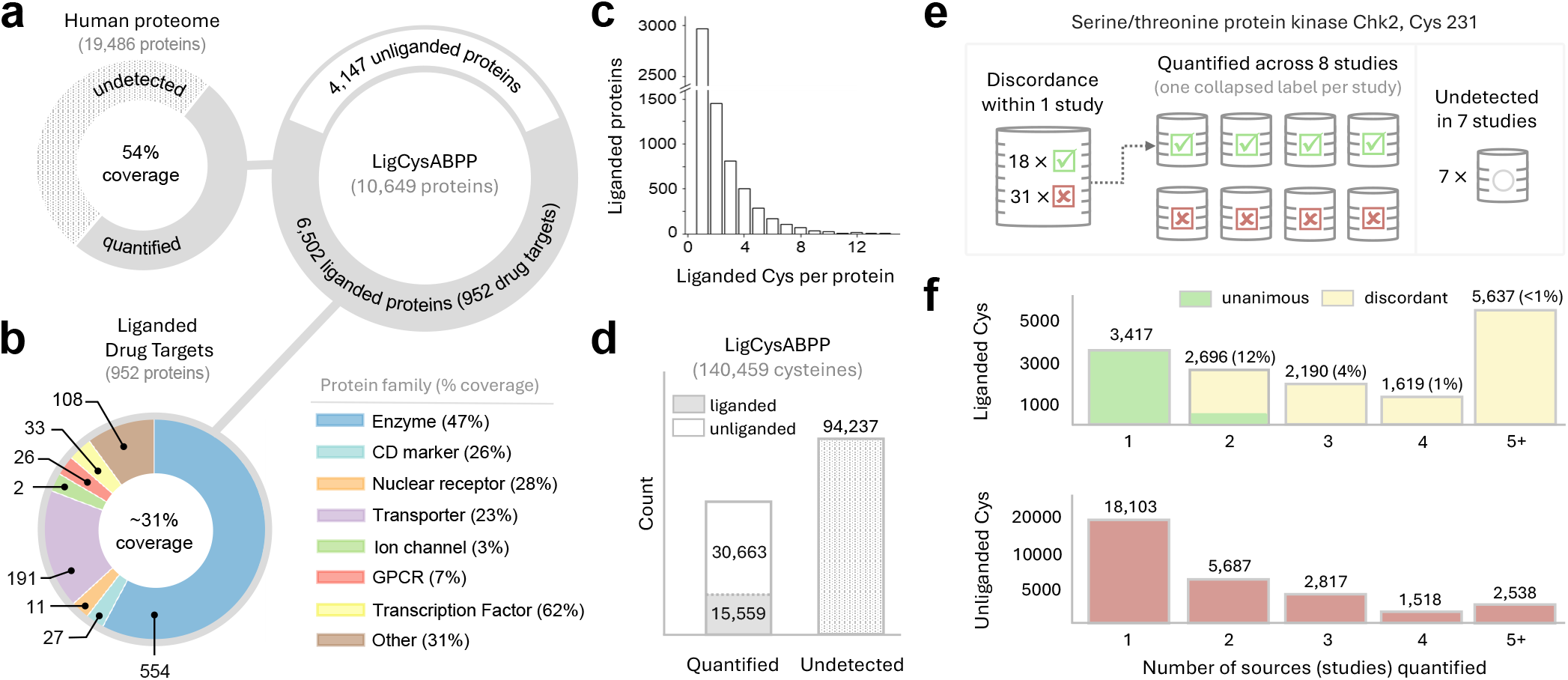
Limited coverage and label heterogeneity in ABPP motivate ML approaches. **a.** Coverage of the human proteome by the ABPP data in the LigCysABPP database. ABPP quantifies 54% of human cysteine-containing proteins, including 6,502 liganded and 4,147 unliganded proteins. Here, liganded refers to having at least 1 positive ABPP record; unliganded refers to having only negative records; and undetected refers to lacking any record. **b**. Functional classes of the 952 liganded drug targets (from Human Protein Atlas ^39^). Percentages indicate coverage of known human dr liganded protein. **d**. Distribution of label heterogeneity (CHEK2 liganded cysteines per CysABPP. **e**. An example e) and discordant study-level labels across eight studies. A source-level label is positive given at least one positive record and negative given all negative records. **f**. Source-level support for liganded and unliganded cysteines. Top, each bin shows the overall count and percentage of cysteines with unanimous label support at the specified number of sources. Bottom, unliganded cysteines by number of sources.

While most ABPP-liganded proteins contain a single liganded cysteine, a sizable fraction harbor 2–4 (Fig. 1c), indicating that ABPP fragments engage secondary pockets, which is extremely valuable for therapeutic discovery. The overlap between proteomic and structural evidence is limited: only 70 proteins are shared with the LC3D database (Supp. Fig. S1), a refined subset of the LigCys3D database ^10^ comprising cysteineliganded co-crystal structures. The minimal over-lap is consistent with the historical focus of structural studies on bacterial and other non-human proteins.

Analysis of LigCysABPP records demonstrated that ligandability labels are variable among different studies. For example, CHEK2 C231 was quantified in 8 and undetected in 7 ABPP studies. Within a single study, both positive and negative records were observed, while across studies, source-level labels vary (Fig. 1e). Here, a source-level label is positive given at least one positive record and negative given all negative records. We analyzed the consensus among source-level labels among liganded cysteines (Fig. 1f top). Agreement on positive labels is rare and drops sharply with more sources: only 12% of cysteines quantified by 2 sources and 4% of those quantified by 3 sources receive unanimous positive labels (Fig. 1f top and Supp. Fig. S2a), highlighting a major challenge in assembling positive-labeled data for ML. By contrast, cysteines with exclusively negative records are abundant, even among those quantified by multiple sources (Fig. 1f bottom). To quantify consensus, we considered cysteines supported by *n* sources and *m* positive records (the *n*S–*m*R criterion). The number of liganded cysteines declines steeply as support requirements increase; at 5 sources, fewer than 30 cysteines have at least 5 positive records (Supp. Fig. S2b). This analysis suggests that requiring agreement across more than 4 sources would be too stringent for ML because it would sharply reduce training-set size.

The variability in ligandability labels likely reflects experimental conditions and, to a lesser extent, cellular state. Supporting this view, a large-scale ABPP study across 400 cancer cell lines (DrugMap ^11^) found that most ligandable cysteines are engaged consistently across cell lines, while a subset shows heterogeneous engagement by a single fragment. We therefore hypothesized that cysteines with consistent labels provide a reliable foundation for training initial ML models for predicting cysteine intrinsic ligandabilities.

## LatentLift and consensus for harmonizing labels

To address label variability, we developed LatentLift, a clustering approach that leverages latent space similarity to reconcile heterogeneous ABPP-derived labels. Here, cysteines are clustered according to pairwise similarity scores computed from residue-specific ESMC embedding vectors (Fig. 2a, Supp. Methods, and Supp. Fig. S4). A ligandability label is assigned to a cluster according to a specified consensus criterion, *n*S– *m*R, where *n*S and *m*R denote the minimum number of supporting sources and records, respectively. To avoid biasing overrepresented cysteines and proteins, only cluster representatives (with the cluster label) were used for baseline model training. Analysis of the Shannon entropy of binary distributions suggested that across consensus levels, the 4S–4R criterion provided the best balance between label confidence and data coverage (Supp. Fig. S3), and was therefore used for creating an initial dataset for training the baseline LigCys model. The LatentLift protocol also enforces leakage control by keeping embedding-similar clusters and sites from the same protein within the same data partition (Fig. 2a and Supp. Methods).

**Figure 2.**
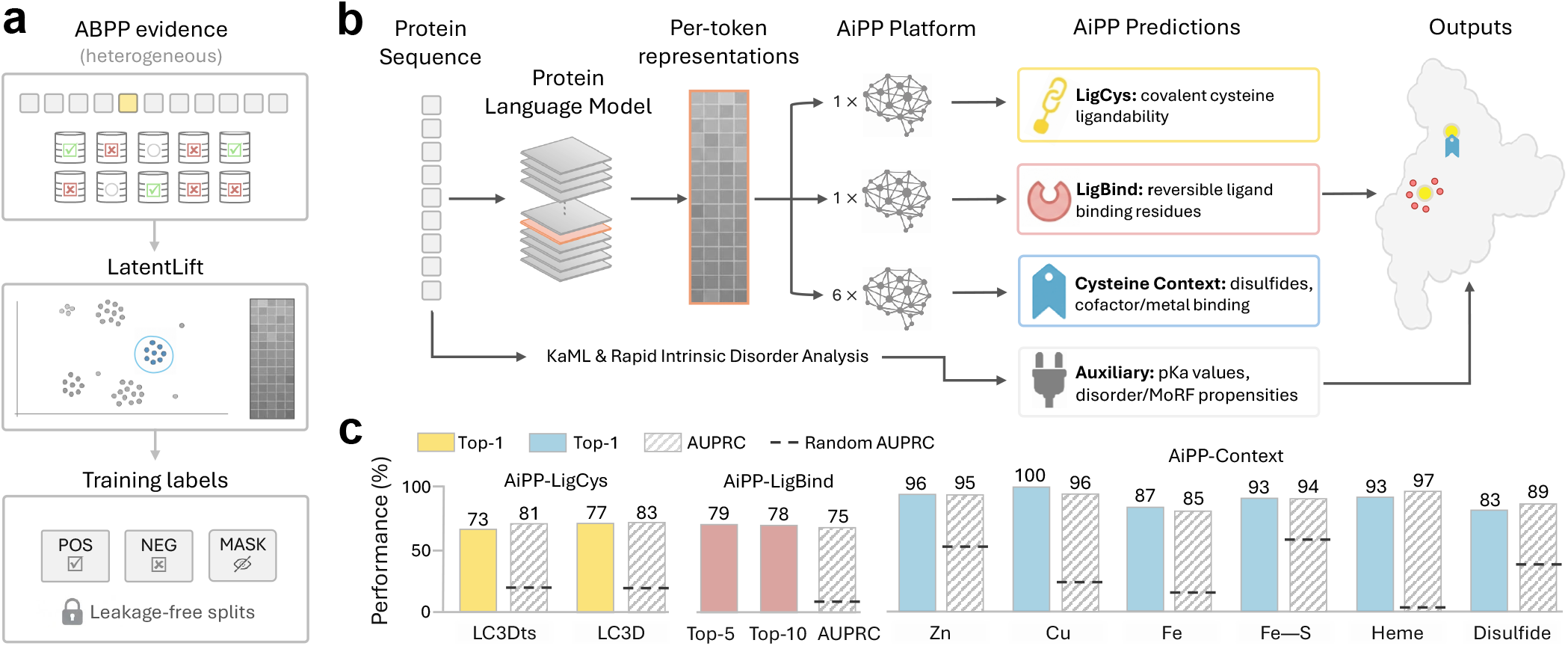
AiPP is a pLLM-derived multi-task platform for identification and contextualization of ligand binding sites directly from protein sequence. **a.** LatentLift reconciles heterogeneous ABPP datasets to produce cysteine ligandability labels and principled data partitions for ML. **b**. Given a protein sequence, a pLLM (ESMC) generates per-residue representations, which are passed to 8 trained AiPP task heads to predict a comprehensive ligandability map, encompassing covalent ligandable cysteines (LigCys), reversible binding residues (LigBind), and cysteine functional context (disulfides and cofactor/metal coordination). Two integrated external sequence-based modules provide disorder/MoRF and cysteine p*K* _a_ annotations. **c**. Performance of the 8 AiPP task heads. Dashed lines indicate random-classifier AUPRCs. LigCys (trained exclusively on ABPP) is evaluated on the structure-based benchmark LC3D and time-stamped external test set LC3Dts. Top-5 and Top-10 recoveries of binding pocket residues are displayed for LigBind.

## A sequence-only baseline LigCys model

We recently demonstrated that a sequence-only multilayer perceptron (MLP) task head on ESMC ^14^ (a state-of-the-art pLLM for protein representation learning ^13,14^) and trained on a small experimental dataset accurately predicts cysteine p*K* _a_ values. ^40^ As cysteine ligandability is similarly governed by local structural and biochemical environment, ^10,41,42^ we hypothesized that it could also be predicted directly from sequence using an ESMC task head. Therefore, we set out to build LigCys, an MLP task head that leverages frozen per-residue embeddings of ESMC and training on label-harmonized ABPP data (Fig. 2b and Supp. Methods). Due to label variability among ABPP data (see later discussion of the ABPP hold-out test), two structure-derived datasets from the PDB were curated for external evaluation: LC3D for benchmarking during model development and the time-stamped LC3Dts for final evaluation and model comparison (see Methods and Supp. Figure S5 for leakage prevention in latent and sequence space). While structural evidence of cysteine ligation represents ground-truth positives, negative labels are weak as they do not represent definitive evidence. We therefore emphasized ranked recovery metrics such as Top-1, defined as the probability that the cysteine with the highest predicted classification score in a protein is a true positive given that the protein has at least one positive cysteine. Top-1 is most relevant in testing against structural evidence, as most proteins in LC3D and LC3Dts contain only one positive cysteine.

Since our analysis (Supp. Fig. S2b and S3) suggests that 4S–4R balances label confidence with data coverage, we trained a baseline Lig-Cys model on the corresponding dataset. On the LC3D benchmark, the model achieved AUPRC and Top-1 of 81% and 75%, respectively (Fig. 3 and Supp. Table S2). Given that ligandability evidenced from crystallography is “orthogonal” to proteomic evidence, this performance suggests that the model generalizes well. This result is particularly notable considering the small training set (389 proteins) and the comparatively large benchmark set (223 proteins). To test how training label consensus affects model generalization, models trained with datasets across consensus levels were compared (Fig. 3 and Supp. Table S2). The 1S–1R model shows the lowest Top-1 recovery (44.5%) despite being trained on the largest dataset. At 1-to-3 source levels, performance improves dramatically as the number of required positive records increases, with the 3S–5R model achieving Top-1 recovery of 73.5%. Interestingly, performance saturates at the 4S–4R level, with the Top-1 recoveries of the 4S–5R and 5S–5R models being 0.9% and 2.6% lower, respectively. This experiment indicates that model performance degrades with increased label variability and reduced dataset size, supporting 4S–4R as the operating consensus criterion for training the baseline model.

**Figure 3.**
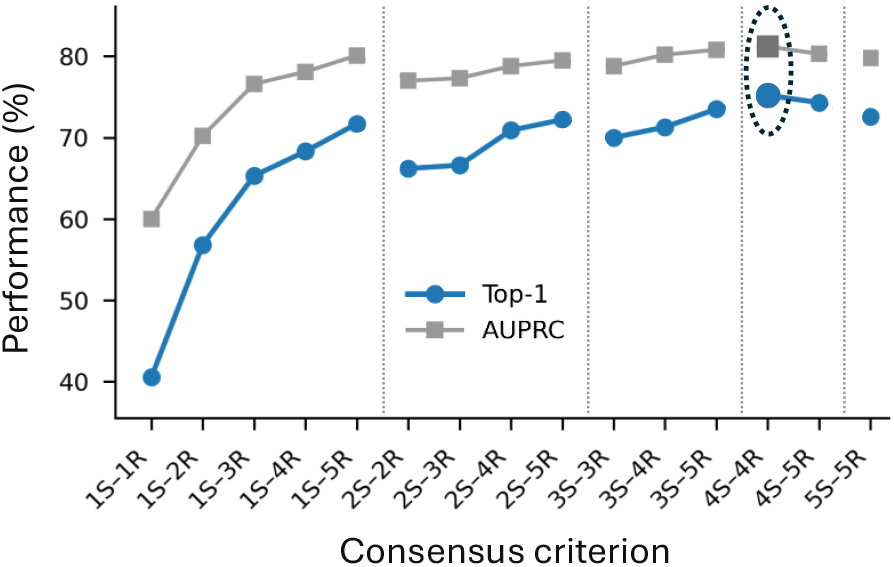
Consensus level of training labels affects model generalization. Performance of LigCys models trained on ABPP datasets across consensus levels (see also Supp. Table S2). Per-protein Top-1 recovery (blue) and AUPRC (gray) on the LC3D benchmark are shown. The 4S–4R model achieved the highest Top-1 and AUPRC (circled).

## Iterative data expansion to improve LigCys

Given the small 4S–4R training set, we asked whether selective data expansion could improve model generalization. We developed two iterative procedures that accept randomly selected ABPP data batches based on whether the new model improves LC3D Top-1 recovery or ABPP AUPRC in cross validation. A truncated 4S–4R set that excludes an ABPP hold-out set of 23 proteins was used to initiate the expansion. The LC3D-guided procedure expanded the training set to 1,099 proteins, and the resulting LigCys-S model increased the LC3D Top-1 score to 79.9% and AUPRC to 84.2% (Supp. Table S2). The ABPP-guided procedure expanded the training set to 1,744 proteins, and the resulting LigCys-A model increased the cross-validation AUPRC enrichment over random guess to 187% while slightly improving the LC3D Top-1 score to 71.3% (Supp. Table S2).

We tested whether adding structural features enhances predictions by training the structure-aware (SA) variants of the LigCys-S and LigCys-A models, which showed 0.9% and 1.3% lower Top-1 scores, respectively (Supp. Table S3). This result supports our hypothesis that ESMC per-residue representations capture the local biochemical, and implicitly, structural environment relevant to cysteine ligandability, consistent with our finding that cysteine p*K* _a_ values can be accurately predicted from sequence using an ESMC task head. ^40^ Hereafter, we will focus on a sequence-only blended model, which makes ensemble predictions using LigCys-A and LigCys-S (Supp. Methods). We will refer to it as the LigCys task head or production model.

## Performance of LigCys task head

Since the LC3D benchmark was used to guide data expansion for the LigCys-S model, we asked whether its performance deteriorates on unseen structural evidence. We therefore evaluated the LigCys task head on LC3Dts, a time-stamped structure dataset (see Methods for leakage prevention). LigCys achieved 73% Top-1 recovery and 81% AUPRC, comparable to the metrics obtained on LC3D (Fig. 2c), significantly outperforming the recent structure-based models TopCySPAL ^9^ (49% Top-1; 69% AUPRC) and CovCysPredictor ^12^ (16% Top-1; 58% AUPRC). The performance gap is likely underestimated, as the alternative models were evaluated using cysteine-liganded co-crystal structures and overlap with the training set was not verified. Recent analysis demonstrated that chemical modification at cysteine tends to increase its solvent exposure, inflating the performance of structure-based models. ^10^ A detailed model comparison is given in Supp. Data file 2.

We further evaluated LigCys on the ABPP holdout consisting of 23 proteins quantified by at least 10 studies. To account for multiple positives per protein, we used Top-4 recovery, which increases with the number of supporting sources (58% with 1 source, 66% with 4, and 78–100% with 7 or more), indicating that LigCys recovers a larger percentage of cysteines consistently liganded in ABPP experiments (Supp. Fig. S6a). Although LigCys does not use structural inputs, we asked whether cysteines in structured regions are better predicted. A stratified analysis showed that Top-4 recovery for cysteines in intrinsically disordered regions is 8% lower than in the structured regions, although this difference may be statistically insignificant given the low abundance of “disordered” cysteines (Supp. Fig. S6d).

Since the LigCys training set uses a 4S–4R consensus and the hold-out test showed higher recovery for cysteines consistently labeled across ABPP studies, we asked whether the model is biased toward heavily supported ABPP hotspots. To address this, we conducted a *post hoc* analysis of LigCys predictions for liganded cysteines with structural evidence (LC3D and LC3Dts), stratifying Top-1 recovery and AUPRC by the number of supporting ABPP sources. These sources were not used in training, as LC3D and LC3Dts neither overlap with nor are homologous to the ABPP training set (Methods). Across support levels from 0 to 4 or above, Top-1 and AUPRC values are similar and show no monotonic increase with greater ABPP support in the corresponding LatentLift clusters (Supp. Fig. S7), arguing against simple recapitulation of ABPP hotspots. Notably, most liganded proteins with structural evidence lack ABPP support, yet LigCys recovered them with Top-1 of 77% and AUPRC of 74% (Supp. Fig. S7), demonstrating model generalization beyond the ABPP data.

## LigBind task head predicts reversible binding residues

Since truly ligandable cysteines are located near a reversible binding pocket, we developed LigBind, a sequence-only ESMC task head for predicting reversible binding propensities of residues (Fig. 2b). LigBind was trained on a new dataset LigBind3D, comprising 687,712 residue-level ligand-binding annotations across 1,998 unique proteins derived from protein-ligand complex structures in the PDB. In the hold-out test, LigBind achieved AUROC, AUPRC and Top-10 recovery of 93.9%, 74.5% and 78.1%, respectively (Fig. 2c).

We hypothesized that LigBind predictions not only complement but may also validate cysteine ligandability assessment. To test this, we applied LigBind to LC3D proteins not in the training set and computed the radial distribution function (RDF) of predicted reversible binding residues surrounding true positive (TP), true negative (TN), false positive (FP), and false negative (FN) cysteines (Fig. 4). The RDF for TPs shows the highest peak near 6 Å, consistent with proximity to reversible pockets, whereas the RDF for TNs is flat, consistent with the lack of a nearby pocket. Notably, the RDF for FNs exhibits the second-highest peak near 4 Å, suggesting that LigBind-predicted reversible binding sites may flag overlooked ligandable cysteines. Conversely, the RDF for FPs is also relatively flat, suggesting that some spurious ligandable cysteines could be filtered by requiring a nearby reversible-binding residue. This analysis suggests that integrating LigCys and LigBind within a multitask platform would enhance the reliability of covalent ligandability assessment.

**Figure 4.**
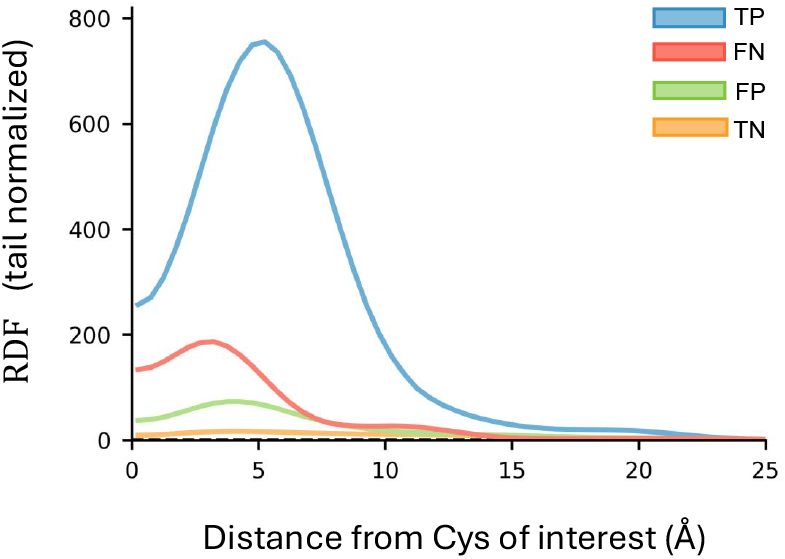
LigBind predictions can rescue false negatives and filter false positives in cysteine ligandability assessment. Radial distribution function (RDF) of LigBind-predicted reversible binding residues around cysteine sites, stratified by the LigCys prediction (TP, TN, FP, FN). Distance is the minimum all-atom Euclidean distance between the cysteine and any predicted ligand-binding residue. RDF is normalized by the mean shell density over the last 20% of the distance range.

## AiPP is a multitask protein-centric toolkit

To provide biological context for cysteine ligandability, we trained six additional ESMC task heads to identify cysteines likely to participate in metal coordination (Zn, Cu, Fe, Fe–S clusters), cofactor (heme) binding, or disulfide bonding. These sequence-only task heads were trained on structural evidence, following the same approach as LigBind (Methods). The hold-out tests demonstrated strong performance across all six task heads (Fig. 2c; Supp. Table S8).

We combined the six cysteine-context task heads with LigCys and LigBind into a unified, sequence-based AI Protein Profiling (AiPP) platform to identify and contextualize ligandable cysteines (Fig. 2b). To further enhance utility and interpretability, AiPP integrates two external sequence-based modules: KaML-ESM, ^40^ which predicts cysteine p*K* _a_ as a proxy for reactivity, and RIDA, ^43^ which predicts intrinsic disorder and molecular recognition features (MoRFs) that undergo folding upon binding. Together, the 8 task heads and 2 external modules provide a comprehensive, protein-centric toolkit to inform cysteine-directed covalent ligand discovery. Below, we apply AiPP to several challenging target classes, while providing further validation through comparison to experiment.

## AiPP maps ligandability of transcription factors

Transcription factors (TFs) are traditionally considered undruggable due to the lack of well-defined pockets and long disordered regions, which may contain MoRFs that mediate interactions with DNAs or other proteins. AiPP is particularly valuable for TFs, as it integrates cysteine ligandability with pocket and MoRF evaluations. We applied AiPP to the TFs highlighted in the DrugMap publication ^11^ and not in the LigCys training set, including FOXA1 (discussed later), FOXA2, SOX10, MYOD1, PAX8, MYB, IKZF1, IRF4, TFAP2C (Supp. Data file 3). FOXA2 C247, SOX10 C71, PAX8 C57, and TFAP2C C209 were predicted as Top-1 or Top-2 ligandable, consistent with the large number of covalent fragment hits (except for FOXA2 C247). ^11^ Notably, AiPP identified MYOD1 C251, MYB C322, IKZF1 C126, and IRF4 C99 as Top-1 ligandable, while these proteins were unliganded in the DrugMap study. ^11^ MYOD1 regulates the expression and circadian amplitude of Bmal1. ^44^ Since C251 lies in a predicted MoRF region, we propose a testable hypothesis that targeting this site may disrupt MYOD1 binding and modulate its function.

## AiPP prioritizes ligandable sites in FOXA1

FOXA1 is a pioneer TF playing critical roles in cancers. ^47^ C258 was predicted as the Top-1 ligandable and hyper-reactive cysteine, adjacent to a MoRF with a reversible binding residue (Fig. 5a). We therefore hypothesized that a C258-directed covalent ligand could engage the dynamic pocket and disrupt the interaction between FOXA1 and DNA. Supporting this hypothesis, an acrylamide-based stereoselective probe was recently discovered that site-specifically reacts with C258 in a DNA-dependent manner and remodels FOXA1’s pioneering activity in prostate cancer cells. ^45^

**Figure 5.**
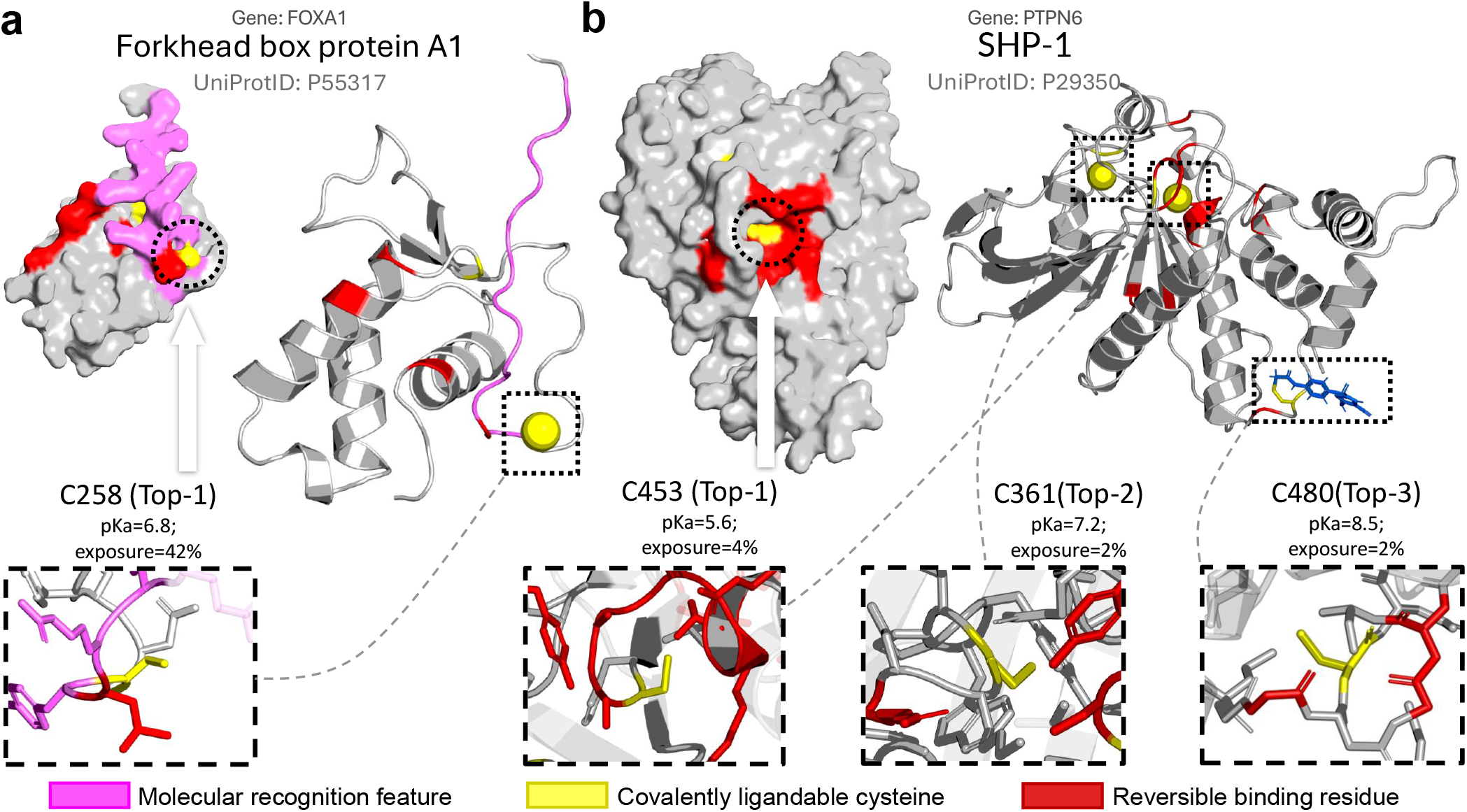
AiPP reveals ligandable cysteines and associated dynamic pockets in undruggable FOXA1 and PTPN6. **a.** The Top-1 ligandable cysteine (C258) in FOXA1 was recently targeted by a stereoselective acrylamide probe **WX-02-23**, modulating chromatin binding. ^45^ C258 is located in the predicted MoRF (magenta) containing a reversible binding residue (red). **b**. Three ligandable sites in PTPN6 were predicted. While C453 (Top-1) is in the active site, C361 (Top-2) and C480 (Top-3) are in cryptic pockets. These sites were undetected or unliganded by ABPP (LigCysABPP database and DrugMap ^11^). We recently developed a first-in-class allosteric covalent inhibitor **M029** targeting C480, demonstrating efficacy in immune cell activation. ^46^ The Top-5 and Top-20 predicted reversible binding residues are shown for FOXA1 and PTPN6, respectively. The ESM3-generated ^13^ structures are shown.

In the DrugMap study, ^11^ C227 was liganded with a modest signal comparable to C258. AiPP, however, ruled out C227, predicting it as a lowconfidence Top-3 site distal to the MoRF region (Supp. Data file 3). This contrast demonstrates that AiPP can help prioritize sites for ligand discovery. Collectively, these TF examples underscore the potential of AiPP to guide covalent ligand discovery against targets lacking well-defined binding pockets. Of note, all four cysteines in FOXA1 were predicted ligandable by TopCySPAL, ^9^ with C258 ranked third. This highlights a known limitation of structure-based models that rely strongly on solvent exposure (see discussion of PTPN6).

## AiPP guides inhibitor discovery for PTPN6

Despite being key cell signaling regulators implicated in several human diseases, protein tyrosine phosphatases (PTPs) have long been deemed undruggable due to challenges of isoform selectivity and bioavailability. ^48^ PTPs utilize a conserved cysteine for catalysis, which can be targeted by covalent inhibitors. ^49^ However, this catalytic cysteine (for example, in PTPN1, PTPN6, and PTPN11) was undetected or unliganded in ABPP experiments (LigCysABPP database and DrugMap, ^11^ Supp. Data file 3). Focusing on PTPN6, a novel intracellular target for small molecule immunotherapy, AiPP predicted three ligandable cysteines with nearby reversible binding residues (Fig. 5b). Notably, the catalytic C453 was Top-1 ligandable (undetected by ABPP according to CysDB, ^50^ LigCysABPP database, and DrugMap ^11^) and surrounded by numerous reversible-binding residues. However, its predicted extremely low p*K* _a_ (5.6) flags it as functional (catalytic nucleophile), ^40^ making it a low-priority site due to potential selectivity concerns.

C361 and C480 were predicted as Top-2 and Top-3 ligandable sites and lie in the vicinity of several reversible-binding residues (Fig. 5b). Note, C480 was undetected according to DrugMap ^11^ or CysDB. ^50^ These cysteines were predicted as hyper-reactive or reactive, with p*K* _a_ values of 7.2 and 8.5, respectively. These predictions guided our medicinal chemistry efforts, leading to the discovery of a first-in-class phenyl chloroacetamide-based covalent allosteric inhibitor **M029** through covalent fragment screening and medicinal chemistry optimization (Supp. Fig. S9). ^46^ **M029** selectively inactivates PTPN6 by covalently binding to non-conserved C480 located in a cryptic pocket far from the active site (Fig. 5b). **M029** is orally active and blocks tumor progression by activating natural killer cells (Supp. Fig. S9).

The prospective identification of both catalytic (positive control) and allosteric cysteines, together with their corresponding reversible binding pockets and distinct nucleophilicities (reflected by their p*K* _a_’s), supports AiPP as a practical toolkit for guiding therapeutic discovery. The fact that these cysteines in PTPN6 and other PTPs were undetected or unliganded by ABPP suggests that AiPP overcomes a limitation of chemical proteomics. To compare with structure-based models, we applied TopCySPAL ^9^ to PTPN6. The two allosteric cysteines were missed, while the catalytic cysteine was ranked fifth (Supp. Data file 2). The tendency of predicting buried cysteines as negatives (C361/C453/C480 in PTPN6 are deeply buried, Fig. 5b) and solvent-exposed cysteines as positives (e.g., false positives in FOXA1) reflects the dependence of structure-based models on solvent exposure. By contrast, LigCys leverages evolutionary information learned by ESMC to predict intrinsic ligandability encoded in protein sequence.

## AiPP illuminates the ligandable proteome

We applied AiPP to the entire human proteome, covering 19,486 cysteine-containing proteins annotated in UniProt – roughly tenfold the LigCys training set, including 9,580 proteins undetected and 3,362 unliganded by ABPP (Fig. 6a and Supp. Fig. S10). AiPP illuminates ABPP-undetected drug targets, including 304 GPCRs, 493 transporters, and 399 enzymes (Fig. 6a). Notably, 567 of the undetected/unliganded drug targets contain highly scored (≥ 0.9) Top-1 cysteines, including 78 GPCRs, 177 transporters, and 207 enzymes (Fig. 6b). In total, AiPP mapped 17,413 Top-1 ligandable cysteines across 17,379 proteins in the human proteome (Fig. 6c, some proteins have multiple Top-1 sites). As a proof of concept for guiding future ligand discovery, below we discuss two GPCRs undetected by ABPP.

**Figure 6.**
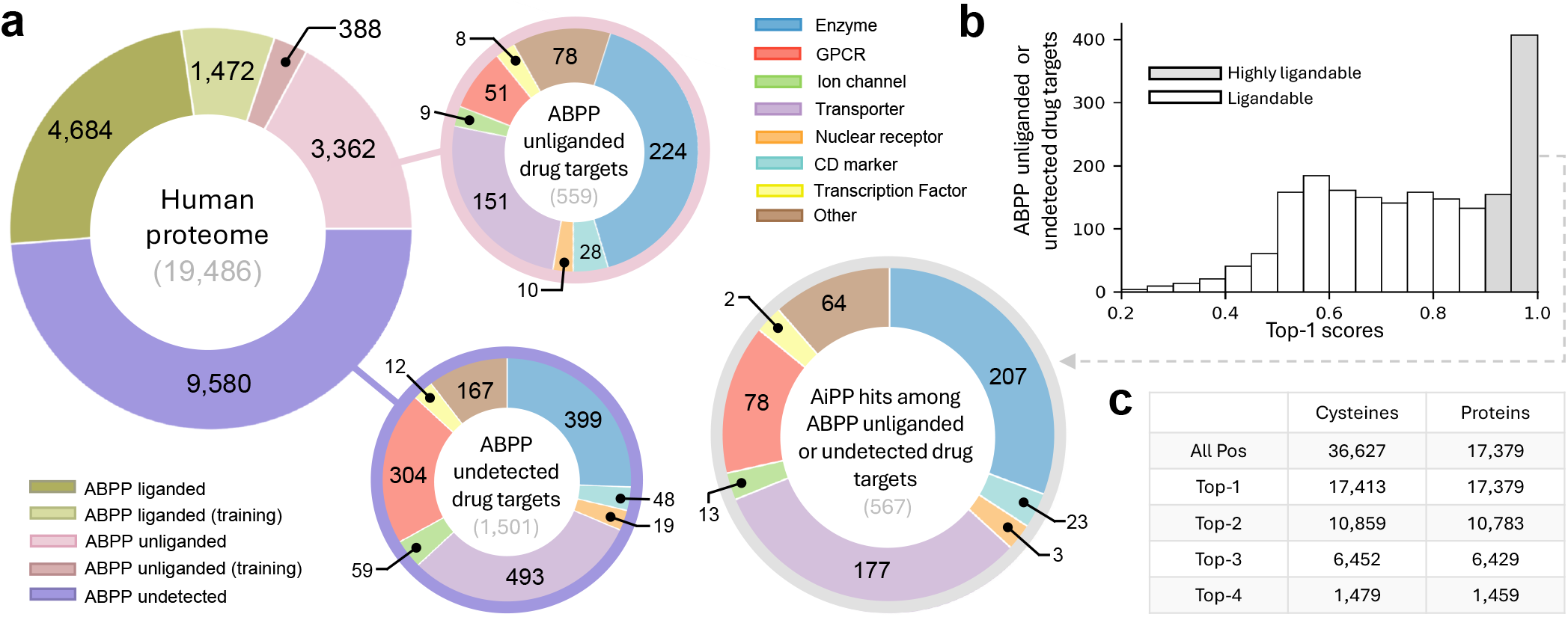
AiPP Atlas illuminates ligandable cysteine sites across the human proteome. **a.** The human cysteine-containing proteome is stratified by ABPP ligandability status and inclusion in LigCys training. ABPP-unliganded and undetected drug targets excluded from model training are stratified by functional class. **b**. Distribution of LigCys scores for the Top-1 cysteines in the ABPP-unliganded or undetected drug targets excluded from model training. Gray bars indicate drug targets with highly scored (≥ 0.9) Top-1 cysteines; the corresponding pie chart displays their functional classes. **c**. Summary of LigCys predictions across the human proteome. All Pos row reports the number of ligandable cysteines and the number of proteins containing at least one ligandable cysteine. The Top-K rows report the corresponding Top-K predictions. Cysteine counts exceed protein counts at each rank as multiple cysteines may share the same rank. Only ligandable sites with a confidence score ≥ 0.9 are included in **b** and **c**.

Orally available drugs for treatment of obesity are highly desirable. One emerging target in the space is melanocortin receptor 3 (MC3R), a regulator of feeding behavior. ^51^ AiPP identified a highly scored Top-1 cysteine, C237^6.30^, located at the cytoplasmic terminus of transmembrane helix 6 (Supp. Fig. S12). The analogous C262^6.29^ in another appetite-regulating receptor NMUR2^52^ was also predicted as the Top-1 ligandable site. Intriguingly, although missed by Lig-Cys, the analogous C347^6.31^ in GLP1R covalently engages a sulfonamide-based molecule, triggering allosteric modulation. ^53^ We hypothesized that covalent ligands targeting C237^6.30^ in MC3R and C262^6.29^ in NMUR2 could disrupt G-protein coupling and modulate receptor activity. We present this testable hypothesis for future experimental verification.

## Concluding Discussion

We developed AiPP, a multitask sequence-based platform for predicting and contextualizing ligandable cysteine sites. AiPP produced a humanproteome atlas that links predicted covalent ligandability with reversible binding pockets, cysteine functional context, predicted reactivity, intrinsic disorder and MoRF annotations. Case studies of historically undruggable proteins FOXA1 and PTPN6 demonstrated AiPP’s utility in guiding hypothesis-driven therapeutic discovery, complementing chemoproteomics. We expect the AiPP atlas to accelerate ligand and biological discovery targeting undruggable proteins and the dark proteome. Integration of protein-centric AiPP with ligand-centric approaches (e.g., molecular docking and generative models for ligand design) provides a new framework for advancing AI drug design. Finally, the LatentLift approach for harmonizing heterogeneous experimental labels is broadly applicable to proteomics-based machine learning.

Current work has several caveats. The LigCys task head focuses on intrinsic ligandability, which is an evolutionarily encoded property of the protein itself, independent of ligand chemistry. LigCys is also by design agnostic to cellular state. Recent studies demonstrated that the reactivity and/or ligandability of a subset of cysteines depends on phosphorylation ^32^ and redox state of the cell. ^11^ Developing cell-state-aware models would require not only understanding the underlying cellular factors (e.g., phosphorylation or redox), but also deconvoluting experimental conditions from true cellular signals in datasets. Such efforts may benefit from the accelerating pace in human proteoform discoveries. ^54^ A related caveat is limited predictive power for mutation sites. For example, AiPP recovered C12 in KRAS-G12C as ligandable and highly reactive (Supp. Data File 3) but missed C220 in p53-Y220C as a false negative. This is likely because KRAS G12C lies proximal to a preexisting dynamic pocket, whereas p53 Y220C destabilizes the protein structure to create a new pocket. ^55^ Future work could incorporate protein mutational stability data, molecular dynamics simulations, or both, to help models recognize cryptic pockets that arise through structural destabilization. AiPP can be improved in the future, e.g., by incorporating continuously growing ABPP data and by adding task heads for predicting other types of covalently ligandable sites such as lysines and tyrosines.

## Methods

### Overview of the AiPP development

AiPP was developed as a sequence-based platform composed of eight task heads and two auxiliary modules. For training and evaluation of the primary LigCys task head, we used the label-harmonized datasets based on LigCysABPP, a new chemoproteomic database. For benchmarking and external validation of LigCys, we curated the LC3D and LC3Dts datasets, which contain structural evidence of covalent cysteine ligandability. For training and evaluation of the LigBind task head, we curated the LigBind3D database extracted from the BioLiP2 database. ^56^ For training and evaluation of the cysteine-context task heads, we curated datasets from the PDB structures containing Zn, Cu, Fe, Fe-S cluster, heme, and disulfide sites. All datasets were converted into a unified site-level record scheme containing a protein identifier (`UID`), a residue position (`ROI`), a binary label (`EXP_BIN`), a label source (`SOURCE`), optional source-specific quantitative metadata, and optional notes (`NOTE`).

### LigCysABPP construction and cysteine label definitions

LigCysABPP, a new chemoproteomic database, was constructed from peptide-level measurements reported in 15 cysteine-directed ABPP studies ^4,8,17–29^ published between 2016 and 2025. Peptide records were expanded to site-level cysteine records, mapped to one or more protein identifiers when required, and validated against full-length protein sequences retrieved from UniProt. We then added undetected cysteines from ABPP-quantified proteins as provisional negative records. After filtering to remove errors, redundancies, and proteins exceeding 2,046 residues in length, the LigCysABPP database contained 703,135 cysteine records for 140,459 cysteine sites across 10,649 proteins (Supp. Fig. S1). Among these cysteines, 46,222 are quantified (liganded or unliganded).

### LatentLift and consensus criterion for label assignment

LatentLift reconciled heterogeneous ABPP-derived labels and enforced leakage-controlled data partitioning. Each unique cysteine site was represented by the per-token embedding from layer 76 of ESMC, a 6-billion-parameter pLLM. Pairwise similarity between embedding vectors was computed as a composite score combining cosine similarity and inverse L1 and L2 distances. Sites with similarity scores above the clustering cutoff were grouped into connected components, and a representative site was selected for each cluster. Cluster labels were assigned from the underlying ABPP records using predefined *n*S–*m*R consensus criteria, where *n*S and *m*R denote the minimum number of supporting sources and supporting records, respectively. For example, under the operating 4S–4R criterion, a cluster was labeled positive if it contained at least four positive records from four distinct sources. Conversely, a cluster was labeled negative if it contained no positive records and at least four negative records from distinct sources. Clusters that did not meet either criterion were left unlabeled. To increase coverage, undetected cysteines contributed provisional negative votes and inherited cluster-level labels. To avoid overweighting large clusters, only cluster representatives and their assigned cluster labels were used for baseline model training. To prevent leakage, clusters were never split across training, validation or test sets, and clusters linked by a shared protein identifier were assigned to the same partition. Clusters containing LC3D records, together with clusters from the same proteins, were excluded from the LigCys training pool. Because embedding similarity captures local biochemical and structural similarity beyond global sequence similarity, LatentLift provides leakage control complementary to sequence-similarity methods such as CD-HIT, ^57^ which may miss proteins with divergent sequences but convergent site-level features.

Application of LatentLift and consensus labeling to LigCysABPP produced training datasets, including 4S–4R, which comprises 384 positive and 642 negative cysteines from 389 proteins (Supp. Table S2). The 4S–4R consensus criterion was chosen as the operational criterion based on a comparison of model performance (Fig. 3 and Supp. Table S2) and Shannon entropy analysis (Supp. Fig. S3). A truncated 4S–4R subset (4S–4R^*t*^) was generated by removing an ABPP hold-out set composed of 23 proteins, each quantified by at least 10 sources. The resulting 4S–4R^*t*^ set contains 354 positive and 574 negative cysteines from 366 proteins and served as the initial training dataset for the iterative data expansion procedures (Supp. Table S3). The ABPP holdout set was then employed to assess the LigCys model across different consensus thresholds and to probe any potential dependence on structural order (Supp. Fig. S6).

### Developing the LigCys task head

We built LigCys as a three-layer MLP trained on frozen 2,560-dimensional per-token embeddings extracted from ESMC layer 76 and label-harmonized ABPP data. To validate the task head architecture, we compared the simple MLP against six alternative architectures: three gated-MLP variants, CNN, dilated CNN, and bidirectional LSTM. Top-1 recovery was comparable between the MLP and the best alternative (gated MLP variant, Supp. Data File 1, SD11). To validate the choice of ESMC, we compared the LigCys performance against models trained with two alternative pLLMs, Ankh3-XL ^58^ and Dayhoff Atlas, ^59^ which reduced Top-1 recovery by 8% and 13%, respectively (Supp. Data File 1, SD12). To assess the effect of pLLM fine-tuning, we compared ESM2 15B (which matched ESMC performance) and its fine-tuned version, which yielded similar results (Supp. Data File 1, SD12). Note, ESMC itself could not be fine-tuned as its weights are not publicly available.

We used two iterative data-expansion procedures to expand the 4S–4R^*t*^ training set (Supp. Table S3). Candidate cysteines were drawn from all 4S–4R^*t*^ clusters, including non-representative members and proteins with only positive or negative labels. The first expansion strategy was guided by the LC3D benchmark. At each iteration, batches of randomly selected cysteines were incorporated into the training set, after which the model was trained and benchmarked against LC3D. A batch was accepted if Top-1 recovery exceeded the previous iteration’s baseline. Across six accepted iterations, Top-1 recovery increased from 70.9% to 79.9% and AUPRC from 77.8% to 84.2%, with no further improvement at iteration 7 (Supp. Table S2). Cross-validated ABPP performance also improved over the baseline: AUPRC enrichment over random guess (PRCE) increased from 74% to 141% (Supp. Table S3). The resulting model, termed LigCys-S, was trained on 1,099 proteins, including 441 positive and 1,345 negative cysteines—nearly three times the size of the starting set. The second strategy used crossvalidation PRCE to guide expansion, accepting only the top-performing batch at each iteration. This increased PRCE to 187%, while LC3D Top-1 recovery initially reached 74.4% before settling at 71.3% (Supp. Table S3). The resulting model, termed LigCys-A, was trained on 2,643 cysteines from 1,744 proteins. Detailed data expansion protocols are given in the Supp. Methods.

Proteins from the LC3D, LC3Dts, and ABPP hold-out sets were excluded from all training and validation datasets during data expansion. The production LigCys model integrated the LigCys-S and LigCys-A predictions.

### Benchmarking and evaluation of LigCys

As structure-derived evidence for cysteine ligandability, we curated the LC3D benchmark and the time-stamped LC3Dts external test set. The initial LC3D set was derived from the LigCys3D database ^10^ by retaining structures with small-molecule covalent ligands of molecular weight ≥ 200 Da, followed by manual inspection and sequence reconstruction from the deposited coordinate files. Covalently modified cysteines were labeled positive, whereas all other cysteines in the same protein were labeled provisional negatives. After LatentLift leakage-control filtering relative to the LigCys training pool, the final LC3D benchmark contained 231 positive and 1,350 negative cysteines in 223 protein-chain identifiers. The LC3Dts test set was constructed using the same procedure, but was restricted to PDB structures deposited between January 1, 2024 and November 11, 2025, and further filtered to remove overlap with the LigCys training pool and LC3D. The final LC3Dts set contained 68 protein-chain identifiers with 77 positive cysteines, and was reserved as a time-stamped external test set for final LigCys evaluation and comparison with structure-based models (Fig. 2c and Supp. Data file 2). For both LC3D and LC3Dts, the maximum global and local sequence similarities to LigCys training proteins were below 21% and 41%, respectively (Supp. Fig. S5).

LigCys was benchmarked on the LC3D dataset and evaluated on the ABPP hold-out and timestamped external LC3Dts dataset. Performance was quantified using ranked recovery metrics, including Top-1 and Top-*k* recovery, together with per-protein AUROC, AUPRC, precision, recall and F1 score. Top-*k* recovery was defined as the fraction of recoverable positive sites appearing among the top *k* predictions within each protein, normalized by the maximum number of recoverable positives at that *k*. We emphasized Top-1 recovery because it directly reflects the practical site-prioritization setting in which only one candidate cysteine may be selected for experimental followup. AUROC, AUPRC and threshold-dependent metrics were interpreted cautiously because negative labels in ABPP and structural datasets are provisional, reflecting absence of observed reactivity, ligation or binding rather than confirmed non-ligandability.

LigCys performance on the external LC3Dts test set and on the experimental PTPN6, FOXA1, and MC3R targets was compared against existing structure-based cysteine ligandability models. (Supp. Data File 2). To evaluate whether Top1 recovery was preferentially enriched at ABPP-supported hotspots, a post hoc analysis of LigCys performance stratified by the degree of ABPP support was conducted (Supp. Fig. S7).

### LigBind and cysteine functional-context task heads

For training the LigBind task head for predicting reversible ligand-binding propensities, we assembled the LigBind3D dataset using the BioLiP2 database, ^56^ which provides protein–ligand complex structures. We selected single-chain protein structures bound to small-molecule ligands with molecular weight between 150 and 600 Da, while filtering out nucleic acids, peptides, ions, crystallization additives, and cofactors. Ligand-contacting residues were identified using a 4.5 Å cutoff based on heavy-atom distances, and retained binding pockets were required to include at least three protein residues in spatial proximity. Ligands covalently attached to cysteine were kept solely for the purpose of defining the surrounding noncovalent pocket contacts. The resulting LigBind3D database contains 687,712 site-level entries spanning 1,998 distinct protein sequences (Supp. Methods and Table S6).

For training task heads for predicting cysteine biological functions, we assembled structure-derived datasets for Zn, Cu, Fe, Fe-S, heme and disulfide-associated cysteines derived from PDB entries. Metal and heme contacts were assigned from residue-target-atom distances using context-specific cutoffs, whereas disulfide bonds were assigned from cysteine sulfur-sulfur geometry and annotation in the PDB file (Supp. Methods and Table S6).

LigBind is a single-layer MLP task head trained as a sequence-based residue classifier, using frozen ESMC layer 76 embeddings as input. A single-layer MLP was chosen over deeper variants based on comparable performance, avoiding unnecessary complexity (Supp. Data File, SD13). The ensemble predictions were obtained by averaging predicted probabilities across independently trained models. The functional-context task heads were trained using the same sequence-only architecture on all eligible residue types. Evaluation and reporting in AiPP were restricted to cysteines because the outputs are used to annotate LigCys predictions. LigBind and context labels were reconciled using LatentLift clustering with permissive propagation of direct structural evidence, and leakage-free partitions were enforced by keeping embedding-similar clusters and shared protein identifiers within the same data partition. LigBind and context outputs were not used to define LigCys ligandability labels; instead, they were reported alongside LigCys scores as orthogonal pocket and functional-context annotations to prioritize cysteine-directed covalent ligand discovery. For additional details, refer to Supp. Methods.

### LigCys scoring and confidence calculation

While the Top-1 voting scheme was applied to the LigCys predictions during model development (including LC3D benchmarking throughout dataset expansion), a consensus-gated residual voting (CGRV) strategy was employed for LC3Dts evaluation and final reporting. CGRV is an ensemble inference scheme designed to suppress isolated outlier “nominations” while preserving consistent secondary and tertiary site-level support. For each protein, ensemble members (i.e., individual models) first nominate their highest-scoring cysteine; when a dominant rank-1 emerges, isolated dissenting rank-1 votes are reassigned to the dominant site. The LigCys score for each cysteine is defined as the total number of CGRV votes received across voting stages divided by the number of ensemble members. Because one ensemble member may contribute votes at more than one stage, LigCys scores may exceed 1. A cysteine that has a nonzero LigCys score is considered ligandable, while cysteines lacking any CGRV evidence are considered unligandable. For ligandable cysteines, confidence scores are derived *post hoc* by applying Benjamini-Hochberg correction to within-protein ensemble nominations, with confidence defined as 1 − *q*. As with the CGRV scheme, confidence scores were applied only in the external LC3Dts evaluation and in reporting LigCys predictions for the human proteome, where highconfidence predictions were defined as those with confidence scores ≥ 0.90, unless otherwise noted.

### AiPP predictions for the human proteome

The final AiPP platform was applied to all cysteine-containing human proteins annotated in UniProt to generate AiPP Atlas 1.0. For each protein, AiPP reported LigCys scores and confidence values, LigBind probabilities, cysteine-context annotations for metal, heme and disulfide-associated sites, KaML-ESM cysteine p*K* _a_ predictions, and RIDA predictions of intrinsic disorder and molecular recognition features. Protein-level summaries were generated by ranking cysteines within each sequence according to LigCys score and annotating each site with predicted binding-pocket proximity, reactivity, disorder and functional-context information. Case-study proteins were analyzed using the same AiPP Atlas outputs by combining within-protein LigCys rank, LigCys confidence, proximity to LigBind-predicted residues, cysteine-context annotations, KaML-ESM p*K* _a_ predictions and RIDA disorder/MoRF predictions. Unless otherwise stated, atlas analyses and case studies used high-confidence LigCys nominations, defined as confidence ≥ 0.90.

## Data availability

The datasets used to train and evaluate the LigCys models are available at Zenodo https://doi.org/10.5281/zenodo.17193766. The LigBind3D datasets used to train and evaluate the LigBind task head are available at Zenodo https://doi.org/10.5281/zenodo.17204148.

The structure-derived cysteine-context datasets used to train and evaluate the metal, heme and disulfide-associated task heads are available at Zenodo https://doi.org/10.5281/zenodo.19863565. Three Supplementary Data files are provided with this paper and mirrored at https://github.com/JanaShenLab/AiPP/. A web server provides access to the AiPP Atlas https://aipp.computchem.org/#explore, a sequence-based prediction engine https://aipp.computchem.org/#predict, and searchable LigCysABPP database https://aipp.computchem.org/#inspect. Additional data and resources are described in the repository documentation.

## Code availability

The AiPP code, including scripts for database construction, LatentLift clustering, model training, model evaluation and proteome-scale prediction, is available at https://github.com/JanaShenLab/AiPP/. Trained model weights are available at https://doi.org/10.5281/zenodo.17210111. Embeddings and supporting files are also archived in Zenodo, links are available in the repository documentation.

## Acknowledgments

We thank Benjamin Cravatt for comments and discussion. We thank Neil Thomas, Chetan Mishra and Marius Wiggert from Evolutionary Scale for facilitating use of ESM Cambrian and ESM3.

## Author contributions

J.S. conceived the idea, designed the research, and oversaw the AiPP development. G.W.D. developed the AiPP platform architecture, LatentLift, the databases, generated the AiPP Atlas, trained and evaluated the models, integrated the external modules, prepared the figures, tables and data files, built and maintained the web server, and prepared the repository releases. D.K. trained the LigCys model, conducted analyses and evaluations, and created figures, data files, and wrote portions of the initial draft. M.S. contributed to analyses, model evaluation, and preparation of figures and data files. R.L. built the initial ABPP database and analyzed the ABPP data. J.L. performed experimental work related to the PTPN6 inhibitor discovery and validation. Z.-Y.Z. oversaw the experimental work and provided figures as well as sections of the original draft. G.W.D. and J.S. prepared the original draft and the revised manuscript. All authors reviewed and approved the final manuscript.

## Additional information

Supplementary Information is available for this paper.

## References

(1) Aebersold, R. et al. How Many Human Proteoforms Are There? Nat. Chem. Biol. 2018, 14, 206–214.

(2) Wishart, D. S. et al. DrugBank 5.0: A Major Update to the DrugBank Database for 2018. Nucleic Acids Res. 2018, 46, D1074–D1082.

(3) Weerapana, E.; Wang, C.; Simon, G. M.; Richter, F.; Khare, S.; Dillon, M. B. D.; Bachovchin, D. A.; Mowen, K.; Baker, D.; Cravatt, B. F. Quantitative Reactivity Profiling Predicts Functional Cysteines in Proteomes. Nature 2010, 468, 790–795.

(4) Backus, K. M.; Correia, B. E.; Lum, K. M.; Forli, S.; Horning, B. D.; González-Páez, G. E.; Chatterjee, S.; Lanning, B. R.; Teijaro, J. R.; Olson, A. J.; Wolan, D. W.; Cravatt, B. F. Proteome-Wide Covalent Ligand Discovery in Native Biological Systems. Nature 2016, 534, 570–574.

(5) Chan, W. C.; Sharifzadeh, S.; Buhrlage, S. J.; Marto, J. A. Chemoproteomic Methods for Covalent Drug Discovery. Chem. Soc. Rev. 2021, 50, 8361–8381.

(6) Abegg, D.; Frei, R.; Cerato, L.; Prasad Hari, D.; Wang, C.; Waser, J.; Adibekian, A. Proteome-Wide Profiling of Targets of Cysteine Reactive Small Molecules by Using Ethynyl Benziodoxolone Reagents. Angew. Chem. Int. Ed. 2015, 54, 10852–10857.

(7) White, M. E.; Gil, J.; Tate, E. W. Proteome-Wide Structural Analysis Identifies Warhead- and Coverage-Specific Biases in Cysteine-Focused Chemoproteomics. Cell Chem. Biol. 2023, 30, 828–838.e4.

(8) Biggs, G. S. et al. Robust Proteome Profiling of Cysteine-Reactive Fragments Using Label-Free Chemoproteomics. Nat. Commun. 2025, 16, 73.

(9) Bonus, M.; Greb, J.; Majmudar, J. D.; Boehm, M.; Korczynska, M.; Nazemi, A.; Mathiowetz, A. M.; Gohlke, H. TopCysteineDB: A Cysteinome-wide Database Integrating Structural and Chemoproteomics Data for Cysteine Ligandability Prediction. J. Mol. Biol. 2025, 437, 169196.

(10) Liu, R.; Clayton, J.; Shen, M.; Bhatnagar, S.; Shen, J. Machine Learning Models to Interrogate Proteome-Wide Covalent Ligandabilities Directed at Cysteines. JACS Au 2024, 4, 1374–1384.

(11) Takahashi, M. et al. DrugMap: A Quantitative Pan-Cancer Analysis of Cysteine Ligandability. Cell 2024, 187, 2536–2556.e30.

(12) Reimer, B. M.; Awoonor-Williams, E.; Golosov, A. A.; Hornak, V. CovCysPredictor: Predicting Selective Covalently Modifiable Cysteines Using Protein Structure and Interpretable Machine Learning. J. Chem. Inf. Model. 2025, 65, 544–553.

(13) Hayes, T. et al. Simulating 500 Million Years of Evolution with a Language Model. Science 2025, 387, 850–858.

(14) ESM Team. ESM Cambrian: Revealing the Mysteries of Proteins with Unsupervised Learning. https://www.evolutionaryscale.ai/blog/esm-cambrian.

(15) Cravatt, B. F. Activity-Based Protein Profiling – Finding General Solutions to Specific Problems. Isr. J. Chem. 2023, 63, e202300029.

(16) Weerapana, E.; Speers, A. E.; Cravatt, B. F. Tandem Orthogonal Proteolysis-Activity-Based Protein Profiling (TOP-ABPP)—a General Method for Mapping Sites of Probe Modification in Proteomes. Nat. Protoc. 2007, 2, 1414–1425.

(17) Bar-Peled, L. et al. Chemical Proteomics Identifies Druggable Vulnerabilities in a Genetically Defined Cancer. Cell 2017, 171, 696–709.e23.

(18) Vinogradova, E. V. et al. An Activity-Guided Map of Electrophile-Cysteine Interactions in Primary Human T Cells. Cell 2020, 182, 1009–1026.e29.

(19) Cao, J.; Boatner, L. M.; Desai, H. S.; Burton, N. R.; Armenta, E.; Chan, N. J.; Castellón, J. O.; Backus, K. M. Multiplexed CuAAC Suzuki–Miyaura Labeling for Tandem Activity-Based Chemoproteomic Profiling. Anal. Chem. 2021, 93, 2610–2618.

(20) Kuljanin, M.; Mitchell, D. C.; Schweppe, D. K.; Gikandi, A. S.; Nusinow, D. P.; Bulloch, N. J.; Vinogradova, E. V.; Wilson, D. L.; Kool, E. T.; Mancias, J. D.; Cravatt, B. F.; Gygi, S. P. Reimagining High-Throughput Profiling of Reactive Cysteines for Cell-Based Screening of Large Electrophile Libraries. Nat. Biotechnol. 2021, 39, 630–641.

(21) Yan, T.; Desai, H. S.; Boatner, L. M.; Yen, S. L.; Cao, J.; Palafox, M. F.; Jami-Alahmadi, Y.; Backus, K. M. SP3-FAIMS Chemoproteomics for High-Coverage Profiling of the Human Cysteinome**. Chem-BioChem 2021, 22, 1841–1851.

(22) Tao, Y.; Remillard, D.; Vinogradova, E. V.; Yokoyama, M.; Banchenko, S.; Schwefel, D.; Melillo, B.; Schreiber, S. L.; Zhang, X.; Cravatt, B. F. Targeted Protein Degradation by Electrophilic PROTACs That Stereoselectively and Site-Specifically Engage DCAF1. J. Am. Chem. Soc. 2022, 144, 18688–18699.

(23) Yang, F.; Jia, G.; Guo, J.; Liu, Y.; Wang, C. Quantitative Chemoproteomic Profiling with Data-Independent Acquisition-Based Mass Spectrometry. J. Am. Chem. Soc. 2022, 144, 901–911.

(24) Yan, T.; Boatner, L. M.; Cui, L.; Tontonoz, P. J.; Backus, K. M. Defining the Cell Surface Cysteinome Using Two-Step Enrichment Proteomics. JACS Au 2023, 3, 3506–3523.

(25) Burton, N. R.; Polasky, D. A.; Shikwana, F.; Ofori, S.; Yan, T.; Geiszler, D. J.; Veiga Leprevost, F. D.; Nesvizhskii, A. I.; Backus, K. M. Solid-Phase Compatible Silane-Based Cleavable Linker Enables Custom Isobaric Quantitative Chemoproteomics. J. Am. Chem. Soc. 2023, 145, 21303–21318.

(26) Koo, T.-Y.; Lai, H.; Nomura, D. K.; Chung, C. Y.-S. N-Acryloylindole-alkyne (NAIA) Enables Imaging and Profiling New Ligandable Cysteines and Oxidized Thiols by Chemoproteomics. Nat. Commun. 2023, 14, 3564.

(27) Burton, N. R.; Backus, K. M. Functionalizing Tandem Mass Tags for Streamlining Click-Based Quantitative Chemoproteomics. Commun. Chem. 2024, 7, 80.

(28) Njomen, E.; Hayward, R. E.; DeMeester, K. E.; Ogasawara, D.; Dix, M. M.; Nguyen, T.; Ashby, P.; Simon, G. M.; Schreiber, S. L.; Melillo, B.; Cravatt, B. F. Multi-Tiered Chemical Proteomic Maps of Tryptoline Acrylamide–Protein Interactions in Cancer Cells. Nat. Chem. 2024, 16, 1592–1604.

(29) Tian, C.; Sun, L.; Liu, K.; Fu, L.; Zhang, Y.; Chen, W.; He, F.; Yang, J. Proteome-Wide Ligandability Maps of Drugs with Diverse Cysteine-Reactive Chemotypes. Nat. Commun. 2025, 16, 4863.

(30) Shannon, D. A.; Weerapana, E. Covalent Protein Modification: The Current Landscape of Residue-Specific Electrophiles. Curr. Opin. Chem. Biol. 2015, 24, 18–26.

(31) Long, M. J. C.; Aye, Y. Privileged Electrophile Sensors: A Resource for Covalent Drug Development. Cell Chem. Biol. 2017, 24, 787–800.

(32) Kemper, E. K.; Zhang, Y.; Dix, M. M.; Cravatt, B. F. Global Profiling of Phosphorylation-Dependent Changes in Cysteine Reactivity. Nat. Methods 2022, 19, 341–352.

(33) Mann, M.; Kulak, N. A.; Nagaraj, N.; Cox, J. The Coming Age of Complete, Accurate, and Ubiquitous Proteomes. Mol. Cell 2013, 49, 583–590.

(34) Gao, M.; Moumbock, A. F. A.; Qaseem, A.; Xu, Q.; Günther, S. CovPDB: A High-Resolution Coverage of the Covalent Protein–Ligand Interactome. Nucleic Acids Res. 2022, 50, D445–D450.

(35) Guo, X.-K.; Zhang, Y. CovBinderInPDB: A Structure-Based Covalent Binder Database. J. Chem. Inf. Model. 2022, 62, 6057–6068.

(36) Zhang, W.; Pei, J.; Lai, L. Statistical Analysis and Prediction of Covalent Ligand Targeted Cysteine Residues. J. Chem. Inf. Model. 2017, 57, 1453–1460.

(37) Du, H.; Jiang, D.; Gao, J.; Zhang, X.; Jiang, L.; Zeng, Y.; Wu, Z.; Shen, C.; Xu, L.; Cao, D.; Hou, T.; Pan, P. Proteome-Wide Profiling of the Covalent-Druggable Cysteines with a Structure-Based Deep Graph Learning Network. AAAS Research 2022, 2022, 9873564.

(38) Boatner, L. M.; Eberhardt, J.; Shikwana, F.; Holcomb, M.; Lee, P.; Houk, K. N.; Forli, S.; Backus, K. M. CIAA: Integrated Proteomics and Structural Modeling for Understanding Cysteine Reactivity with Iodoacetamide Alkyne. ACS Chem. Biol. 2025, 20, 1669–1682.

(39) Uhlen, M.; Oksvold, P.; Fagerberg, L.; Lundberg, E.; Jonasson, K.; Forsberg, M.; Zwahlen, M.; Kampf, C.; Wester, K.; Hober, S.; Wernerus, H.; Björling, L.; Ponten, F. Towards a Knowledge-Based Human Protein Atlas. Nat. Biotechnol. 2010, 28, 1248–1250.

(40) Shen, M.; Dayhoff, G. W.; Shen, J. Protein Electrostatic Properties Are Fine-Tuned Through Evolution. 2025.

(41) Liu, R.; Yue, Z.; Tsai, C.-C.; Shen, J. Assessing Lysine and Cysteine Reactivities for Designing Targeted Covalent Kinase Inhibitors. J. Am. Chem. Soc. 2019, 141, 6553–6560.

(42) Liu, R.; Verma, N.; Henderson, J. A.; Zhan, S.; Shen, J. Profiling MAP Kinase Cysteines for Targeted Covalent Inhibitor Design. RSC Med. Chem. 2022, 13, 54–63.

(43) Dayhoff II, G. W.; Uversky, V. N. Rapid Prediction and Analysis of Protein Intrinsic Disorder. Protein Sci. 2022, 31, e4496.

(44) Hodge, B. A.; Zhang, X.; Gutierrez-Monreal, M. A.; Cao, Y.; Hammers, D. W.; Yao, Z.; Wolff, C. A.; Du, P.; Kemler, D.; Judge, A. R.; Esser, K. A. MYOD1 Functions as a Clock Amplifier as Well as a Critical Co-Factor for Downstream Circadian Gene Expression in Muscle. eLife 2019, 8, e43017.

(45) Won, S. J.; Zhang, Y.; Reinhardt, C. J.; Hargis, L. M.; MacRae, N. S.; DeMeester, K. E.; Njomen, E.; Remsberg, J. R.; Melillo, B.; Cravatt, B. F.; Erb, M. A. Redirecting the Pioneering Function of FOXA1 with Covalent Small Molecules. Mol. Cell 2024, 84, 4125–4141.e10.

(46) Qu, Z. et al. Discovery of a First-in-Class Covalent Allosteric SHP1 Inhibitor with Immunotherapeutic Activity. Angew. Chem. Int. Ed. 2026, 65, e25126.

(47) Liu, N.; Wang, A.; Xue, M.; Zhu, X.; Liu, Y.; Chen, M. FOXA1 and FOXA2: The Regulatory Mechanisms and Therapeutic Implications in Cancer. Cell Death Discov. 2024, 10, 1–15.

(48) Zhang, Z.-Y. Drugging the Undruggable: Therapeutic Potential of Targeting Protein Tyrosine Phosphatases. Acc. Chem. Res. 2017, 50, 122–129.

(49) Abdo, M.; Liu, S.; Zhou, B.; Walls, C. D.; Wu, L.; Knapp, S.; Zhang, Z.-Y. Seleninate in Place of Phosphate: Irreversible Inhibition of Protein Tyrosine Phosphatases. J. Am. Chem. Soc. 2008, 130, 13196–13197.

(50) Boatner, L. M.; Palafox, M. F.; Schweppe, D. K.; Backus, K. M. CysDB: A Human Cysteine Database Based on Experimental Quantitative Chemoproteomics. Cell Chem. Biol. 2023, 30, 683–698.e3.

(51) Patel, T. P. et al. Melanocortin 3 Receptor Regulates Hepatic Autophagy and Systemic Adiposity. Nat. Commun. 2025, 16, 1690.

(52) Botticelli, L.; Micioni Di Bonaventura, E.; Del Bello, F.; Giorgioni, G.; Piergentili, A.; Quaglia, W.; Bonifazi, A.; Cifani, C.; Micioni Di Bonaventura, M. V. The Neuromedin U System: Pharmacological Implications for the Treatment of Obesity and Binge Eating Behavior. Pharmacol.Res. 2023, 195, 106875.

(53) Cong, Z. et al. Molecular Insights into Ago-Allosteric Modulation of the Human Glucagon-like Peptide-1 Receptor. Nat. Commun. 2021, 12, 3763.

(54) Smith, L. M.; Agar, J. N.; Chamot-Rooke, J.; Danis, P. O.; Ge, Y.; Loo, J. A.; Paša-Tolić, L.; Tsybin, Y. O.; Kelleher, N. L.; The Consortium for Top-Down Proteomics The Human Proteoform Project: Defining the Human Proteome. Sci. Adv. 2021, 7, eabk0734.

(55) Mavridi, D.; Funk, J. S.; Balourdas, D.-I.; Krämer, A.; Khan Tareque, R.; Timofeev, O.; Spencer, J.; Stiewe, T.; Joerger, A. C. Targeting the P53 Cancer Mutants Y220C, Y220N, and Y220S with the Small-Molecule Stabilizer Rezatapopt. Cell Death Dis. 2026,

(56) Zhang, C.; Zhang, X.; Freddolino, L.; Zhang, Y. BioLiP2: An Updated Structure Database for Biologically Relevant Ligand– Protein Interactions. Nucl. Acids Res. 2024, 52, D404–D412.

(57) Fu, L.; Niu, B.; Zhu, Z.; Wu, S.; Li, W. CD-HIT: Accelerated for Clustering the next-Generation Sequencing Data. Bioinformatics 2012, 28, 3150–3152.

(58) Alsamkary, H.; Elshaffei, M.; Elkerdawy, M.; Elnaggar, A. Ankh3: Multi-Task Pretraining with Sequence Denoising and Completion Enhances Protein Representations. 2025.

(59) Yang, K. K.; Alamdari, S.; Lee, A. J.; Kaymak-Loveless, K.; Char, S.; Brixi, G.; Domingo-Enrich, C.; Wang, C.; Lyu, S.; Fusi, N.; Tenenholtz, N.; Amini, A. P. The Dayhoff Atlas: Scaling Sequence Diversity for Improved Protein Generation. 2025,

