## Supplemental Methods, Tables, and Figures for "Illuminating the Ligandable Human Proteome with AI Protein Profiling"

**6 Benchmarking LigCys against prior methods** 37

**7 External modules and annotations** 39

**Supplementary Figures** 41

Figure S1. Overview of proteins and cysteines in LigCysABPP database . . . 42

Figure S2. Analysis of cysteine ligandability across ABPP sources and records . . . 42

Figure S3. Source threshold effects on label variability and coverage . . . . . . . . . . . . . 44
Figure S4. Similarity metrics used for LatentLift clustering . . . . . . . . . . . . . . . . . . . 45
Figure S5. Sequence identity leakage checks . . . . . . . . . . . . . . . . . . . . . . . . . 46
Figure S6. Evaluation of LigCys on the ABPP hold-out set and disorder analysis . . . . . 47
Figure S7. Post hoc analysis of LigCys performance versus off-training ABPP support . . 48
Figure S8. RDF of LigBind-predicted binding residues around cysteine sites . . . . . . . . 49
Figure S9. Characterization and efficacy of the first-in-class covalent allosteric inhibitor of SHP-1 . . . . . . . . . . . . . . . . . . . . . . . . . . . . . . . . . . . . . . . . 50
Figure S10. Functional classification and LigCys Top-1 prediction scores . . . . . . . . . 51
Figure S11. Reversible binding propensity of residues in MC3R and GLP-1R . . . . . . . 52

**Supplementary Tables** 52
Table S1. Literature sources and experimental conditions in LigCysABPP . . . . . . . . . 52
Table S2. Evaluation of consensus thresholds for ABPP data selection in LigCys model training . . . . . . . . . . . . . . . . . . . . . . . . . . . . . . . . . . . . . . . . 55
Table S3. Model performance in the iterative training set expansion . . . . . . . . . . . . . 56
Table S4. Features used in training the structure-aware variants of LigCys . . . . . . . . . 57
Table S5. Performance of the LigBind task head . . . . . . . . . . . . . . . . . . . . . . . 59
Table S6. Data and clustering details for training LigBind and functional-context task heads 60
Table S7. Task-specific AiPP hyperparameters . . . . . . . . . . . . . . . . . . . . . . . 60
Table S8. Performance of functional-context task heads . . . . . . . . . . . . . . . . . . . 60

**References** 62

### Supplementary Methods

### 1 LigCys and LigBind databases and record schema

To train and evaluate AiPP, we assembled three complementary databases capturing covalent ligandability and reversible ligand-binding signals. Specifically, we constructed: (i) LigCysABPP, a cysteine ligandability database derived from peptide-level measurements from 15 cysteine-directed activity-based protein profiling (ABPP) studies published between 2016–2025;[1–15] (ii) LC3D, a LigCys3D-derived subset of proteins with covalently liganded cysteines supported by co-crystal structures;[16] and (iii) LigBind3D, residue-level annotations of residues involved in reversible ligand binding derived from BioLiP2.[17]

**Definition of records in the databases.** All databases curated in this work were converted into a unified schema of structured, site-level records to enable consistent downstream processing across modules. Each record includes: (1) a source-specific protein unique identifier (`UID`); (2) a residue of interest (`ROI`) corresponding to a 1-based sequence position; (3) a binary site label (`EXP_BIN`; `POS` or `NEG`); (4) source-specific quantitative metadata (i.e., the raw competition ratio `R` and the study-defined threshold `EXP_THR`); (5) a source identifier (`SOURCE`); and (6) an optional free-text note (`NOTE`; e.g., ligand identity, molecular weight, probe identity). Fields not applicable to a given source were set to sentinel values for harmonization and were never interpreted as quantitative measurements. For structure-derived records (LC3D and LigBind3D), we set `R` = 999 and `EXP_THR` = 0 to denote direct structural evidence. For undetected sites (cysteines in ABPP-quantified proteins with no detectable probe signal and therefore no competition measurement; Section 1.1), we set `R` = -1 and `EXP_THR` = 0.

#### 1.1 LigCysABPP: curation of chemoproteomic data

**Curation of an ABPP database.** We manually curated peptide-level results from 15 ABPP studies published between 2016–2025.[1–15] Cysteine-directed ABPP provides indirect, chemoproteomic measurements of cysteine ligandability at scale. However, ABPP readouts depend on experimental context (e.g., cell type, redox state, probe/warhead chemistry, and competition design). Accordingly, site labels can vary across studies and should be treated as weak supervision. We therefore retain per-record provenance and assay metadata (`SOURCE` and `NOTE`). AiPP later aggregates evidence across studies via embedding-based clustering and multi-source consensus labeling (Section 3.1).

For each study, we extracted peptide-level entries and converted them into the structured, site-level records described above. Entries referencing multiple cysteine positions or multiple UIDs were expanded into one record per (`UID`, `ROI`) pair while preserving the original study metadata. When a single ROI mapped to multiple UIDs, the ROI was duplicated across those UIDs. This yielded 683,192 records spanning 58,704 unique cysteine sites (UID, ROI pairs) across 14,417 distinct proteins (UIDs).

**Protein sequence reconstruction and record validation.** To ensure consistency between sequences and experimental annotations, all site-level records were subject to source-specific sequence reconstruction and cysteine identity validation. Each UID was mapped to its full-length canonical sequence using a local UniProt FASTA cache or the UniProt REST API (`https://rest.uniprot.org`). Records were excluded if the reported ROI did not correspond to a cysteine in the retrieved sequence, if the sequence contained ambiguous residues (e.g., ‘X’), or if the accession was unavailable. In UniProt, ‘X’ denotes an unknown amino acid; we removed such records to ensure complete, biologically coherent sequences for modeling. This protocol yielded 671,515 validated records (out of 683,192 initial records), covering 53,867 unique cysteine sites across 12,745 proteins.

**Consolidation of UIDs and filtering of derived records.** To eliminate redundancy, we collapsed UIDs referring to identical sequences or strict subsequences, selecting a single representative UID per group. This step yielded 650 UID groups in total (89 identical sequences and 561 subsequences), most of which corresponded to UniProt isoform or redundant accessions. In UniProt, isoform accessions (e.g., "-1", "-2") can reflect distinct transcript annotations even when the translated protein sequence is identical; we therefore grouped records by amino-acid sequence rather than accession. All records were then updated to reference the representative UID as the canonical identifier.

Next, we filtered records derived from ambiguous peptide mappings: any record whose canonical UID did not appear in at least one unambiguous record (single UID, single ROI) was discarded, removing 1,607 records corresponding to 1,078 UIDs.

**Treatment of undetected cysteine sites in quantified proteins.** We added 114,568 undetected records, representing cysteine sites in otherwise ABPP-quantified proteins that showed no detectable probe signal and therefore lacked a quantitative competition measurement. These were encoded with `EXP_BIN` = `NEG` and `SOURCE` = `UNDETECTED` for downstream consensus processing, but were treated as provisional negative evidence rather than hard negatives. They reflect the absence of observed reactivity rather than definitive evidence of non-ligandability and were retained only for downstream consensus labeling (Section 3.1).

**Final records in the LigCysABPP database.** Finally, we removed records from proteins shorter than 30 residues or longer than 2,046 residues. This cutoff reflects the 2,048-token input limit of the protein large language model (pLLM) used in this work, which includes the beginning-of-sequence (BOS) and end-of-sequence (EOS) tokens. This removed 81,341 records describing 25,485 unique cysteines (UID, ROI pairs) from 1 short protein and 367 long proteins. The final ABPP database contained 703,135 records: 608,898 validated peptide-derived records and 94,237 undetected records (after filtering). Together, these records span 140,459 unique cysteine sites (UID, ROI pairs) across 10,649 representative UIDs (unique sequences after consolidation). Of these sites, 46,222 were ABPP-quantified (15,559 liganded and 30,663 unliganded). The remaining 94,237 sites were undetected and were retained as provisional negative evidence for downstream consensus labeling. At the protein level, 6,502 ABPP-quantified proteins (UIDs) contained at least one liganded cysteine.

#### 1.2 LC3D and LC3Dts: curation of covalent co-crystal structures

**Curation of the LC3D dataset.** LC3D was constructed as a curated subset of LigCys3D, a database previously reported by our group,[16] providing a direct structural complement to the ABPP data. LigCys3D compiles cysteines observed to form covalent bonds with ligands in experimentally resolved structures from the worldwide Protein Data Bank (wwPDB). For LC3D, we retained only structures with covalent ligands of molecular weight $\geq 200$ Da. Each structure was manually inspected, and entries deemed incorrect (e.g., wrong residue ID, missing ligand, terminal cysteines) or unsuitable (e.g., cysteines forming disulfide bonds or helical staples, or cysteines linked to peptides or proteins) were removed or corrected.

**Sequence reconstruction and cysteine validation.** Rather than relying on LigCys3D-provided annotations, we extracted the referenced PDB identifiers, downloaded the corresponding coordinate files, and independently derived cysteine annotations. Sequences were reconstructed directly from atomic coordinates using our in-house tool (`gSeq`), which derives a per-chain construct sequence from modeled residues in file order and inserts residues annotated as unobserved in the wwPDB record (mmCIF `_pdbx_unobs_or_zero_occ_residues` and, when available, legacy PDB `REMARK 465`) to improve consistency between the reconstructed sequence and the coordinate model. When both mmCIF and legacy PDB files were available in the local wwPDB mirror, gSeq reconciled the two reconstructions by favoring cross-format agreement and internal consistency with the corresponding polymer sequence (SEQRES/SEQRES-like), defaulting to mmCIF on ties. Site-level annotations were validated against these reconstructed sequences to confirm residue identity and positional accuracy. We did not remap sequences to UniProt—in contrast to our original LigCys3D processing[16]—because remapping can discard experimental details (e.g., engineered mutations or truncations). Our goal was to preserve the exact protein sequence used in the structural experiment rather than revert to a potentially mismatched canonical reference. After UID consolidation and validation, the curated LC3D collection contained 316 POS LigCys3D-derived cysteine records across 275 PDB chains (UIDs).

**Provisional labeling of unliganded cysteines in co-crystal structures.** To achieve complete cysteine coverage per structure, we added all other cysteine residues in the validated sequences as provisional negatives (EXP_BIN = NEG). As in LigCysABPP, these negatives reflect the absence of observed covalent modification rather than definitive evidence and were retained for downstream label reconciliation (Section 3.1). After addition of these negatives, LC3D comprised 1,643 site-level cysteine records across 275 PDB chains. To ensure each PDB chain contained both positive and negative sites, we retained only chains with at least one negative cysteine.

**Leakage prevention.** To prevent leakage with the LigCys training set, we first applied the LatentLift leakage-control procedure described in Section 3.1 and in the paragraph “Pruning LC3D-containing clusters to prevent data leakage”. Specifically, LC3D-containing clusters were excluded from the ABPP-derived training pool, and any remaining clusters linked to the same proteins were removed through the transitive UID-based partitioning scheme. As an additional conservative failsafe for the structural benchmark, we then pruned a small number of LC3D proteins that lay slightly above our prespecified global or local sequence-identity cutoffs relative to the LigCys training set. The final LC3D benchmark set contained 223 unique UIDs and 1,581 unique UID–ROI pairs: 231 positives (14.6%) and 1,350 negatives, with no masked sites. The global and local sequence similarities between LC3D and LigCys training proteins are below 21% and 41%, respectively (Fig. S5).

**Records in LC3D dataset.** All entries were converted into the structured record format used for the LigCysABPP database. Each record uses a UID in the format `PDBID-CHAINID`, with ROI values referencing sequence positions from the corresponding PDB structure. Ligand-specific metadata (e.g., molecular weight, three-letter PDB ligand code) were stored in the `NOTE` field. ABPP-specific fields such as `EXP_THR` and `R` were assigned sentinel values (`EXP_THR` = 0, `R` = 999) to denote direct structural evidence.

**Curation of time-stamped LC3Dts external test set.** To assess generalization to covalently liganded proteins supported by newly deposited structures, we assembled an additional external test set, LC3Dts, using the same LC3D inclusion criteria and processing pipeline but restricting the PDB pull to structures with covalently liganded cysteines deposited between January 1, 2024 and November 11, 2025. After validating the pull-down, deduplicating entries within the subset, we ensured no leakage by removing proteins overlapping with the LigCys training pool or the final LC3D benchmark set. As an additional conservative failsafe, we also removed the small number of proteins whose local or global sequence identity lay slightly above the corresponding prespecified cutoffs relative to the LigCys training proteins. The global and local sequence similarities between LC3Dts and LigCys training proteins are below 21% and 41%, respectively (Fig. S5). The final LC3Dts contained 69 unique sequences spanning 68 UIDs and comprising 77 covalently liganded cysteines. Note, there are two sequences (each containing a different liganded cysteine) related to a single UID. LC3Dts was constructed after model development, serving as an external test set for further evaluating LigCys and comparison to structure-based models.

#### 1.3 LigBind3D: reversible binding-site curation

**Curation of the LigBind3D database.** To develop models that predict reversible ligand-binding pockets, we curated an orthogonal structural database of reversible ligand-binding residues from BioLiP2,[17] a manually curated resource of biologically relevant protein–ligand complexes derived from wwPDB structures. Here, "reversible" refers to noncovalent pocket recognition. When a ligand is covalently linked to a cysteine in the deposited structure, we retain the ligand only to define the surrounding reversible pocket contacts that precede covalent capture. We selected single-chain systems—wwPDB entries whose deposited coordinate model contains exactly one protein polymer chain—containing small molecules with molecular weights between 150–600 Da. We excluded entries associated with nucleic acids, peptides, ions, crystallization agents, or cofactors. Structures containing multiple protein chains (homomers, heteromers, or antibody fragments), even when the ligand contacts only one chain, were excluded. This restriction reduces conformational confounding: because our models take only a single amino-acid sequence as input, they cannot condition on which complex is present or distinguish complex-specific conformations. It also avoids ambiguity from inter-chain binding sites and matches our sequence-only modeling setup, which operates on one protein chain at a time. Each retained chain was recorded as the UID (`PDBID-CHAINID`). For each selected PDB entry, we applied a custom structural annotation pipeline (`PickPocket`) to validate biologically relevant protein–ligand interactions. `PickPocket` resolves `LINK` records, merges "multi-residue" ligands, detects missing atoms, renumbers residues, and filters out artifacts. Ligands annotated in mmCIF but missing from the coordinate file (e.g., covalently attached fragments) were reconciled when appropriate. Ligands were retained only if they (i) remained within the target molecular weight range after reconstruction, (ii) were not covalently linked to protein atoms—*except when the covalent bond involved a cysteine residue, in which case we retained the adduct as a pocket-defining ligand because covalent capture requires prior localization of the ligand in the binding pocket*—and (iii) formed coherent binding pockets involving at least three spatially proximal ($\leq$ 4.5 Å) residues.

**Records in LigBind3D database.** For each structure that passed all filtering steps, we reconstructed the full protein sequence directly from the atomic coordinate file, as in building the LC3D dataset. Next, ligand-contacting residues were labeled based on a 4.5 Å heavy-atom distance threshold. This operational definition captures reversible pocket contacts in resolved complexes. For cysteine-linked adducts, we defined pocket residues by the same 4.5 Å proximity rule. These residues were converted into structured, site-level records with `EXP_BIN` = `POS`. Each record uses a UID in the format `PDBID -CHAINID`, with ROI values referencing sequence positions from the corresponding PDB structure. Ligand-specific metadata (e.g., 3-letter ligand code, molecular weight) were stored in the `NOTE` field. As with the LC3D database, sentinel values (`EXP_THR` = 0, `R` = 999) were used to denote direct structural evidence. All other residues in the same chain were added as negatives with `EXP_BIN` = `NEG` and are distinguishable via the `NOTE` field. As in other databases, these negatives are weakly labeled, reflecting the absence of observed interaction rather than confirmed non-binding, and were retained as provisional for downstream label reconciliation (Section 3.2).

Overall, the final LigBind3D database comprised 687,712 site-level records across 1,998 unique sequences (UIDs). This database provides a direct structural perspective on reversible ligand recognition, complementing the chemoproteomic ABPP data and the covalent structural data in LC3D. However, unlike LigCys3D, which restricts ROIs to cysteines, LigBind3D treats every residue in each protein as a potential ROI, reflecting the broader diversity of reversible ligand interactions.

### 2 Functional-context annotations for cysteines

To support biological interpretation of LigCys outputs, we derived residue-level annotations from wwPDB structures and trained sequence-only models to reproduce these labels. Although these structure-derived labels are defined over all residue types in each parsed chain, evaluation and reporting in this work are restricted to cysteines because these predictors are used to annotate LigCys outputs. We report per-cysteine model outputs alongside LigCys scores as an interpretability layer. All annotations in this section were generated from a local mirror of wwPDB mmCIF coordinate files cloned on November 11, 2025. Record fields and sentinel conventions follow those used for structure-derived databases in Section 1 (LC3D and LigBind3D). Below we describe the context-specific labeling rules. These labels were clustered and partitioned using the embedding-based framework described in Section 3.2 prior to training the corresponding sequence-only models (Section 4.1).

#### 2.1 Metal and cofactor coordination

**ZNBind3D, CUBind3D, FEBind3D, FESBind3D, and HEMBind3D databases.** To annotate residues involved in metal and cofactor coordination, we implemented an automated mmCIF parsing pipeline (`metalMiner`) that uses atomic coordinates and standard wwPDB annotations to identify residue–metal/cofactor contacts.

For each structure, we restricted candidate atoms to protein polymer chains (with automatic fallbacks when polymer annotations were missing or incomplete) and considered only heavy-atom donors (O, N, or S). Hydrogen atoms were excluded, and atoms with occupancy $< 0.5$ were ignored. For entries containing multiple coordinate models (e.g., NMR ensembles), contacts were retained only if supported by at least half of the models.

We defined a residue–target contact when any qualifying donor atom lay within a fixed distance cutoff of a specified target atom, identified by wwPDB chemical component identifiers (e.g., a Zn/Cu/Fe ion, or the Fe atom within a heme or Fe–S cluster component). Distance cutoffs were 3.5 Å for Zn and Cu, and 3.0 Å for Fe, Fe–S clusters, and heme. Using this procedure, we generated five structure-derived metal/cofactor contact databases: (i) **ZNBind3D** (Zn; component `ZN`); (ii) **CUBind3D** (Cu; components `CU` and `CU1`); (iii) **FEBind3D** (Fe; component `FE`); (iv) **FESBind3D** (Fe–S clusters; Fe atoms in the canonical Fe–S components `SF4`, `FS4`, `F3S`, and `FES`, using the Fe atom annotations in each component); and (v) **HEMBind3D** (heme; Fe atoms in heme components `HEM`, `HEC`, `HEA`, and `HEB`).

For each database, positives were residues in contact with the corresponding target atom under the geometric rule above; residues in the same parsed protein chains that were not in contact were treated as negatives. We also recorded whether each contact was explicitly annotated by depositors as a coordination link in mmCIF connection records, and used this flag as a consistency check on automated contact calls.

#### 2.2 Disulfide bonding

**SSBind3D database.** To annotate whether predicted cysteine sites are likely constrained by disulfide formation, we curated a structure-derived disulfide database (SSBind3D) from the wwPDB mmCIF mirror described above. Disulfide bonds were identified by an automated parsing pipeline (`disulfideMiner`) that uses atomic coordinates and standard wwPDB annotations to detect disulfide-linked cysteine pairs.

We used the same chain filtering described in Section 1.2. Candidate sulfur atoms were taken from cysteine `SG` atoms and from modified cysteine residues whose wwPDB chemical components list `CYS` as the parent residue. Selenocysteine-derived disulfides were not included. Atoms with occupancy $< 0.5$ were excluded, and alternate locations were handled with a compatibility rule that permits pairing when altloc identifiers match or when one or both atoms lack an altloc identifier.

Putative disulfide bonds were called when two qualifying sulfur atoms satisfied an `SG`–`SG` distance window of 1.90–2.20 Å. For entries containing multiple coordinate models (e.g., NMR ensembles), we scanned all models and retained only bonds supported in at least 50% of models. To ensure consistent structural quality, we excluded structures whose best reported resolution was poorer than 3.0 Å (using the best available resolution value in the wwPDB record).

In addition to geometry-based calls, we recorded whether each disulfide pair was explicitly annotated by depositors in mmCIF connection records as an independent consistency check on the automated assignments. We did not discard structures with no disulfide evidence under the criteria above (by either geometry or depositor annotation). Instead, cysteines from these structures were used as negative examples. This prevents the model from implicitly assuming that disulfides are ubiquitous and better reflects the strong class imbalance in disulfide prevalence.

### 3 LatentLift: clustering, partitioning, and label reconciliation

At the core of AiPP is **LatentLift**, a pLLM embedding-based framework for clustering residue sites and reconciling heterogeneous labels across sources. We represent each residue by a 2,560-dimensional token embedding extracted from layer 76 of ESM Cambrian (ESMC),[18] a 6-billion-parameter protein large language model computed from sequence alone. LatentLift uses these embeddings to (i) group biochemically similar sites, (ii) assign cluster-level labels and reconcile conflicts, and (iii) enforce leakage-free partitioning by keeping highly similar sites (and proteins linked by shared identifiers) in the same data split. We use the resulting clusters and partitions to construct training and evaluation datasets for LigCys and LigBind, and we apply the same clustering and UID-exclusive partitioning policy to the functional context databases (Section 2) to define leakage-free splits for training the corresponding cysteine context predictors (Section 4.1).

#### 3.1 LatentLift for LigCys

**LatentLift clustering.** To group biochemically similar cysteine sites, we clustered validated site-level records using their ESMC-derived embeddings. Each unique cysteine site—defined by a (UID, ROI) pair—was represented as a single node in embedding space, encoded by a 2,560-dimensional vector. All records corresponding to the same site were collapsed into a single node for clustering, ensuring that redundant evidence did not influence cluster formation or representative selection.

We computed a composite similarity score $S$ between embedding vectors, defined as $S = \frac{1}{3}\big(C + D_1 + D_2\big)$, where $C$ is the cosine similarity between vectors, and $D_1$ and $D_2$ are the inverse L1 and inverse L2 distances: $D_1 = 1/(1 + L_1)$, $D_2 = 1/(1 + L_2)$. This composite score requires agreement in both direction (cosine similarity) and magnitude (L1/L2 distances), and equals 1 only when two embeddings are identical. We used a clustering cutoff of $S \geq 0.3$, which corresponds approximately to cosine similarity $\sim 0.9$ (Supplementary Fig. S4b). The distribution of $S$ and the cutoff location are shown in Supplementary Fig. S4a.

An undirected edge was drawn between two nodes if $S \geq 0.3$, and clusters were formed as connected components using a union–find algorithm. Within each cluster, a representative site was selected as the node with the highest average similarity $S$ to all other members. Any node with $S < 0.3$ relative to the representative was pruned. Pruned nodes were iteratively re-clustered using the same procedure until convergence. Any remaining unclustered nodes were assigned as singleton clusters.

After clustering, all original records associated with each node were fully re-expanded so that each cluster included the complete set of source-level records corresponding to its constituent sites. This ensured that downstream cluster-level label assignment could integrate all available evidence during label derivation. The final result was a set of compact, non-overlapping clusters of cysteine sites sharing high representation-level similarity, each anchored by a representative node used for consensus labeling and model training.

LatentLift uses fixed thresholds (the clustering cutoff $S \geq 0.3$ and the $n$S–$m$R consensus criteria, where $n$S denotes the minimum number of distinct supporting sources and $m$R denotes the minimum number of supporting records). For labeling, we analyze the coverage–consensus trade-off, label stability, and benchmark impact across $n$S–$m$R settings (Supplementary Fig. S3 and Supplementary Table S2), motivating 4S–4R as the fixed operating point used throughout.

**Leakage-free partitioning in LatentLift.** To ensure that evaluation reflects true generalization, LatentLift enforces leakage-free partitioning using the same pLLM embedding-space similarity used for clustering. Conventional global sequence identity filtering (e.g., CD-HIT[19]) can reduce redundancy, but it does not control for functionally convergent and/or structurally similar ligandable sites embedded within otherwise dissimilar proteins. Consistent with this, the maximum global sequence identity between training-set proteins and proteins in the external LC3D and LC3Dts sets is 21.9% and 21.2%, respectively (Supplementary Fig. S5a,c), below commonly used CD-HIT thresholds (e.g., 30–40%).

LatentLift clustering operates on contextual ESMC embeddings, which capture local biochemical, structural, and evolutionary information. As a result, residues with similar ligandability profiles tend to cluster together—even across proteins lacking detectable sequence homology, yielding a higher-resolution view than traditional methods based on sequence alignment.

To prevent leakage, LatentLift applies two complementary safeguards during dataset partitioning. First, each cluster is treated as an indivisible unit and is never split across training, validation, or test sets. Second, LatentLift enforces protein-level (UID) exclusivity: all clusters containing records from the same protein are assigned to the same partition. This constraint was applied transitively—if any two clusters shared a UID (even through different residues), both were placed in the same partition. Operationally, we treat

UID–cluster membership as a graph and assign data splits at the level of its connected components, ensuring that any proteins and clusters linked directly or indirectly through shared membership are indivisible across train/validation/test partitions. As a result, no protein appears in more than one dataset partition, and no structurally or biochemically similar residues are shared between training and evaluation sets. Together, these safeguards prevent both representation-level and protein-level leakage, ensuring that reported performance metrics reflect true predictive power on unseen proteins and local contexts.

**Label assignment for cysteine clusters.** To reconcile heterogeneous ABPP-derived cysteine ligandability labels while preserving a strictly external structural benchmark, we assigned consensus labels to ABPP-derived clusters within the full set of LatentLift clusters. To avoid data leakage, clustering was performed jointly on all validated ABPP and LC3D site-level records, defining a single embedding-space similarity graph used both for leakage prevention and for identifying ABPP sites that are embedding-similar to LC3D benchmark proteins. We then applied conservative pruning to prevent leakage from LC3D into ABPP-derived training labels (see *Pruning LC3D-containing clusters to prevent data leakage* below). In addition, we used ABPP records within the same embedding clusters as LC3D sites only for post hoc analyses (e.g., stratifying LC3D performance by the number of ABPP sources supporting a site; Supplementary Fig. S7); this stratification does not modify LC3D labels or the leakage-free train/validation/test splits.

LC3D labels were taken directly from the curated co-crystal structures and were treated as fixed, per-site annotations at the level of the specific PDB chain and residue position. Clustering was not used to propagate labels among LC3D sites: if multiple LC3D records fell in the same cluster, each retained its original `POS` or `NEG` label, and LC3D evaluation always used these input labels. Likewise, we did not use LC3D labels to assign labels to ABPP-only sites. Operationally, we flagged any cluster containing at least one LC3D record as LC3D-containing and excluded such clusters from ABPP consensus labeling and from the training pool (see *Pruning LC3D-containing clusters to prevent data leakage* below).

Clusters that did not contain any LC3D records were evaluated using a consensus voting scheme, in which positive or negative labels were assigned only if they met predefined thresholds across both record counts and sources (e.g., 1S–1R, 1S–2R, . . . , 4S–4R, 4S–5R, . . . ; see Supplementary Table S2). As described in the Supplementary Fig. S3 and Supplementary Table S2, we evaluated the coverage–stringency trade-off and benchmark performance across consensus levels and selected 4S–4R as a conservative operating threshold; this choice was then held fixed for all training, evaluation, and expansion procedures reported in this work. For positives, a cluster was labeled positive once the required threshold was met (e.g., at the 1S–2R level, at least two `POS` records from one source; at the 4S–4R level, at least four `POS` records from different sources). Negatives followed the same thresholds but required the stricter condition that *all* records in the cluster were labeled `NEG` (i.e., no positive votes). Clusters that did not meet these criteria—or exhibited conflicting or insufficient evidence—were masked and excluded from training. Applying this rule, for example, at the 4S–4R level, 940 clusters were labeled `POS` and 5,170 clusters labeled `NEG`, covering 162,809 site-level records describing 11,473 distinct cysteines across 11,118 unique proteins. The remaining clusters, including 52,010 singletons, were masked. For baseline LigCys training, we trained on one representative site per labeled cluster; thus, the “4S–4R” counts quoted in the main text refer to cluster representatives rather than all underlying site-level records. Training on one representative per cluster prevents repeatedly measured sites from being overweighted during optimization while preserving broad biochemical coverage through cluster expansion.

**Pruning LC3D-containing clusters to prevent data leakage.** Following cluster-label assignment, we applied a conservative pruning step to eliminate potential data leakage. Specifically, all clusters containing any LC3D records were removed from the training pool after label assignment. Additionally, we identified all UIDs represented in these clusters and removed any remaining clusters that shared a UID with them—even if those clusters did not contain LC3D records themselves. This strict pruning ensured that no ABPP-derived cluster contained cysteine sites from proteins already seen in the LC3D-resolved set, eliminating indirect leakage via shared sequence context. Importantly, LC3D was withheld as an orthogonal benchmark set due to its high-confidence structural origin and minimal redundancy with ABPP-derived data. Excluding all LC3D-associated proteins from training ensures that subsequent model evaluation on this benchmark set reflects true generalization to unseen, structure-supervised ligandable sites.

The remaining labeled clusters comprised a distilled, nonredundant training set derived exclusively from ABPP records. This set preserved biochemical and experimental diversity while enforcing rigorous separation between structure-derived and sequence-derived labels. All records within each retained cluster inherited the resolved cluster label, providing consistent and leakage-free supervision for model training.

**Pruning an ABPP hold-out set.** We created an ABPP hold-out set to provide a second, independent check during LigCys training-data expansion (Section 5). As defined in the main text, it contains 23 proteins that were each measured by at least 10 independent ABPP sources and that have at least one positive and one negative cysteine. All site-level records for these proteins were flagged as hold-out and never used for model training or validation. In all steps (clustering, label assignment, and expansion), any cluster that contained a record from a hold-out protein—and any other cluster sharing that protein identifier (UID)—was placed entirely in the hold-out partition.

We used this hold-out set in two ways: (i) to monitor the LC3D-guided expansion for signs of overfitting to LC3D, and (ii) to evaluate how the ABPP-guided cross-validation performs on unseen, well-measured proteins. Because cysteines within these proteins have different numbers of supporting measurements, we further stratified the hold-out cysteines into nine $n$S–$m$R subsets ($n = 1, \ldots, 9$) to examine how heterogeneous labeling affects fair performance assessment and error analysis. These diagnostics were purely observational and did not influence batch acceptance, hyperparameter choices, or model selection. A complete list of hold-out UIDs and cysteine records is provided in Supplementary Data SD6–SD7.

**Robustness to heterogeneous ABPP sources.** Because ABPP-derived labels vary across experimental contexts (e.g., cell type, redox state, probe/warhead chemistry, and competition design), we assessed whether LigCys performance depends disproportionately on any single ABPP study. We performed a leave-one-source-out analysis in which we rebuilt the baseline 4S–4R training set after excluding one ABPP source at a time, retrained LigCys under the same protocol, and evaluated on the leakage-free LC3D benchmark. Across all 15 ablations, performance varied modestly (LC3D Top-1 recovery: 72.22%–77.35%; range = 5.13 percentage points), indicating that no single ABPP context is required for strong generalization. Full per-source ablation results are provided in Supplementary Data SD4.

**Independent heterogeneity analysis using DrugMap.** To interpret disagreements that can arise under heterogeneous ABPP supervision, we compared AiPP consensus labels and LigCys scores against an independent set of "heterogeneous cysteines" from the DrugMap study.[20] We intersected DrugMap heterogeneous cysteines with LigCysABPP (excluding DrugMap as a source) and assigned each overlapping site an AiPP consensus label under the same 4S–4R criterion used throughout this work (Supplementary Fig. S10a). Separately, for prediction analysis (Supplementary Fig. S10b), we scored all DrugMap heterogeneous cysteines with valid protein sequences using LigCys ($N = 237$), including the $N = 228$ sites that overlap LigCysABPP and additional DrugMap sites outside the curated LigCysABPP universe. Among the 228 overlapping sites, most were masked (144) or positive (39), with 45 labeled negative, consistent with our design choice that high-confidence negatives should reflect sites that are consistently non-ligandable across sources rather than sites that are context-variable. We then compared LigCys ensemble classifications for AiPP 4S–4R negatives versus DrugMap heterogeneous cysteines (Supplementary Fig. S10b). LigCys predictions were derived from the final production ensemble using CGRV (Section 4.3): a LigCys score of 0 indicates the site received no votes across CGRV stages, whereas any nonzero LigCys score indicates it received at least one vote and is classified as positive under the LigCys decision rule. We used this comparison to assess whether sites annotated as heterogeneous in DrugMap are preferentially classified as ligandable by LigCys under the same decision rule.

#### 3.2 LatentLift for LigBind and functional-context databases

For LigBind3D and the functional context databases, we used the same embedding-space clustering and leakage-free partitioning as in Section 3.1, but resolved cluster labels using a permissive propagation rule (i.e., 1S–1R) rather than the $n$S–$m$R consensus thresholds used for LigCys.

**Label assignment for LigBind clusters.** LigBind3D records were processed independently from ABPP and LC3D records to derive cluster-level labels for residues involved in reversible ligand binding. All LigBind3D records were clustered using the same embedding-based similarity metric and threshold as in the LigCysABPP workflow.

Clusters were resolved using a permissive propagation rule: if any record in a cluster was labeled `POS` (i.e., within 4.5 Å of a ligand in a resolved structure), the entire cluster was assigned a positive label. Otherwise—if no records were labeled `POS`—the cluster was labeled `NEG`. No vote thresholds, source requirements, or masking logic were applied. This approach treats ligand proximity as direct structural evidence and enables propagation of contact labels to structurally or biochemically similar residues across isoforms, homologs, or convergent domains.

From the clustering step, we obtained 581,493 clusters (515,751 singletons; 65,742 multi-member) spanning 1,998 UIDs and 687,712 LigBind3D records. Of these, 29,158 clusters were uniformly positive and 549,028 uniformly negative, while 3,307 contained mixed labels. Uniform clusters were defined as those in which all records shared the same label. At the record level, labels totaled 39,044 positive and 648,668 negative prior to reconciliation (5.7% positive; $N : P = 16.6 : 1$). To enforce within-cluster consistency, we performed a single-pass reconciliation that converted $\texttt{NEG}\rightarrow\texttt{POS}$ assignments, yielding 45,617 positive and 642,095 negative records overall (6.6% positive; $N : P = 14.1 : 1$). These cluster-level labels constitute the high-confidence structural annotations used for LigBind model training and benchmarking.

**Label assignment for functional context clusters.** The functional context databases (ZNBind3D, CUBind3D, FEBind3D, FESBind3D, HEMBind3D, and SSBind3D; Section 2) were clustered using the same embedding-based similarity metric, threshold, and leakage-free partitioning policy used for LigBind3D (i.e., identical representation-based clustering and UID-exclusive splits). Cluster-level labels were resolved using the same permissive propagation rule: if any record in a cluster is labeled POS (direct structural contact or bond annotation), the cluster is assigned POS; otherwise, it is assigned NEG. To enforce within-cluster consistency, we applied the same single-pass reconciliation used for LigBind, converting $\texttt{NEG}\rightarrow\texttt{POS}$ assignments in mixed-label clusters when required by the cluster label. A quantitative summary of database scale, clustering outcomes, and label prevalence is provided in Supplementary Table S7.

### 4 Models: architecture, training, and evaluation

**Shared partitioning and evaluation protocol across all models.** Unless otherwise stated, all residue-level models in AiPP—LigCys, LigBind, and the functional context annotation models (ZNBind, CUBind, FEBind, FESBind, HEMBind, and SSBind)—share the same leakage-free partitioning policy defined by LatentLift (Section 3.1, *Leakage-free partitioning in LatentLift*), which enforces cluster-level indivisibility in pLLM embedding space and protein-level (UID) exclusivity across training, validation, and evaluation sets. Ensemble inference is also shared across tasks (Section 4.3).

#### 4.1 Sequence-only models

**LigCys-seq models.** LigCys-seq models were trained to perform per-site inference given the input sequence. The input protein sequence was tokenized and encoded by ESMC,[18] and per-token embeddings were extracted from layer 76, yielding a 2,560-dimensional vector for each cysteine. ESMC per-token embeddings were used in frozen form as input to a three-layer feedforward network with hidden dimensions 1,024, 512, and 256, each followed by GELU activation. We applied layer normalization to the input embeddings and dropout ($p = 0.5$) before the final projection to a scalar ligandability logit (sigmoid). Embeddings were obtained via the hosted ESM Forge encoder and were not updated during training; only the task-specific MLP head was trained.

Training was formulated as binary classification with residue labels 1 (consensus positive), 0 (consensus negative), and 2 (masked). Masked residues were excluded from loss and metric computation; when a binary label was required (e.g., ranked recovery within a protein), masked residues were treated as negative. We used AdamW with an initial learning rate of $1 \times 10^{-5}$ and weight decay of $1 \times 10^{-5}$. The learning rate schedule multiplied the learning rate by 0.9 after epoch 1 and by 0.7 after epoch 2; the learning rate was unchanged after epoch 3, and then multiplied by 0.5 after each epoch from 4 through 9. Each model was trained for 10 epochs without early stopping. For each of 10 distinct train/validation splits, we trained 20 independent replicates (different random seeds) for ensemble evaluation and variance estimation. We also explored additional readout-head variants in ablation studies (see *Head-depth and local-context ablations*; Supplementary

Data SD11).

Mini-batches were constructed per protein, initially containing all cysteines from a single input sequence. To address class imbalance, a 10:1 sampling ratio of negative to positive residues was enforced using a custom sampler. If a given protein contained too few residues to populate a full batch, additional residues were randomly sampled from other proteins in the training set to meet the batch size requirement. The loss function combined binary cross-entropy with focal modulation ($\alpha = 0.66$, $\gamma = 1.0$). Model checkpoints were selected based on maximum validation AUPRC and subsequently evaluated on held-out LC3D benchmark data using ensemble inference (Section 4.3). To benchmark LigCys-seq against alternative protein language models, we trained the same LigCys readout head using final-layer per-token embeddings from multiple PLMs (including ESM2-15B, Ankh3XL, and Dayhoff-3b-UR90) under the same protocol (trained on the baseline 4S–4R dataset prior to any expansion and evaluated on LC3D with the same per-protein metrics and ensembling approach). Results are summarized in Supplementary Data SD12.

**LigBind-seq models.** LigBind-seq models share the same backbone embedding procedure as LigCys-seq models, using fixed 2,560-dimensional per-token representations from ESMC layer 76. A simpler single-layer feedforward network was applied, consisting of a layer normalization, a GELU-activated linear transformation (2,560 $\rightarrow$ 2,560), dropout ($p = 0.1$), and a final projection to a scalar logit. We retained this minimal readout head for LigBind-seq and did not perform an extensive task-head sweep.

Classification is binary, with residue-level labels of 1 (binding) or 0 (non-binding). No masked residues were present in the LigBind3D database. The loss function was adaptive focal loss with 20 confidence bins, initial $\gamma = 1.0$, bin updates every 5 epochs, a calibration-penalty term $\lambda = 0.1$, and $\gamma$ clamped to [0.5, 5.0]. As with LigCys-seq models, all ESMC embeddings were frozen. Models were trained for up to 100 epochs using AdamW (learning rate $1 \times 10^{-5}$, weight decay $1 \times 10^{-5}$), with early stopping triggered after

20 epochs of no improvement in validation AUPRC. For each of 10 distinct train/validation splits, 20 models with different random seeds were trained, yielding a total of 200 LigBind-seq models.

Training batches were constructed per protein, enforcing a 15:1 ratio of negative to positive residues using the same sampling strategy as for LigCys-seq models. If a single protein did not contain enough residues to meet the minimum batch size of 256, additional samples were drawn randomly from other proteins. Evaluation metrics were computed on ensemble predictions (Section 4.3) and included per-protein AUROC, AUPRC, F1 score, and top-$k$ recovery.

**Functional context sequence-only models.** The functional context annotation models (ZNBind, CUBind, FEBind, FESBind, HEMBind, and SSBind) share the same sequence-only architecture and training protocol as LigBind-seq. Fixed per-token ESMC layer 76 embeddings (2,560 dimensions) were passed through the same LigBind-seq readout head to produce a per-residue probability for the corresponding context label (e.g., metal coordination, cofactor coordination or disulfide bonding). All ESMC embeddings were frozen and only the readout head was trained. During training and validation, these models were trained on all residue types, preserving the native label prevalence and class imbalance of the underlying structural annotations. These models follow the same LatentLift leakage-free split policy described above (cluster indivisibility and UID exclusivity) and use the same ensemble inference strategy as LigBind-seq, with probability averaging across ensemble members. Task-specific differences were limited to (i) adaptive focal-loss hyperparameters (including $\alpha$ and $\gamma$) and (ii) the negative-to-positive sampling ratio, set based on label prevalence in each training dataset. A summary of per-task dataset scale, clustering outcomes, and label prevalence is provided in Supplementary Table S6. Task-specific sampling ratios and loss settings are summarized in Supplementary Table S7, and per-task benchmark performance is reported in Supplementary Table S8.

**Model-complexity ablations.** To assess whether LigCys-seq performance was limited by the simplicity of the readout architecture, we evaluated higher-capacity sequence-only models trained on the baseline 4S–4R dataset prior to any expansion and evaluated on the LC3D benchmark using the same ensembling and per-protein evaluation protocol. First, we tested alternative task-head architectures while keeping the pLLM embeddings frozen, comparing the production MLP head to gated-MLP variants, a 1D CNN head, a dilated CNN/temporal convolutional network (TCN) head, and a bidirectional LSTM head.

Second, we tested whether adapting the pLLM backbone itself improved LigCys-seq performance. Because the production LigCys-seq model uses frozen ESMC embeddings obtained through the hosted Forge encoder, we performed this backbone-adaptation ablation using qLoRA on the locally trainable ESM2_15B backbone (`esm2_t48_15B_UR50D`). LoRA adapters were inserted throughout the ESM2_15B transformer using rank $r = 64$, scaling factor $\alpha = 128$, and adapter dropout $p = 0.1$. QLoRA used a quantization block size of 32, bfloat16 compute precision, double quantization, gradient checkpointing, and a paged 8-bit AdamW optimizer. The pretrained ESM2_15B backbone weights were not directly updated.

The PEFT model used a single-hidden-layer MLP task head with dimensions $5120 \rightarrow 1024 \rightarrow 1$ and dropout $p = 0.5$. Training used the same residue-level binary classification framework and focal-loss formulation as LigCys-seq, with a 6:1 negative-to-positive sampling ratio. Both the qLoRA adapter parameters and task head were optimized with learning rate $1 \times 10^{-5}$ and weight decay $1 \times 10^{-4}$. Models were trained for 15 total epochs: during the first 5 epochs, adapters were disabled and only the task head was optimized; during the remaining 10 epochs, the qLoRA adapters and task head were optimized jointly. Checkpoints were selected by validation AUPRC and evaluated on the held-out LC3D benchmark.

Performance from these model-complexity controls, including AUROC, AUPRC, precision, recall, F1 score, and Top-1 recovery, is reported in Supplementary Data SD11 and was used to assess whether increased readout-head complexity or PEFT-based backbone adaptation justified replacing the lightweight frozen-embedding LigCys-seq model.

#### 4.2 Structure-aware variants

Sequence-only models use ESMC embeddings. We also tested structure-aware (SA) extensions that incorporate explicit structural information. We trained SA variants only as ablations for comparison to LigCys-seq and LigBind-seq; all reported results in this work use sequence-only models. For LigCys, we used a disjoint adaptive gating unit (DAGU) to combine sequence embeddings with engineered structural features. For LigBind, we concatenated geometric embeddings with ESMC vectors.

**ESM3-generated structural models.** For SA ablations and case-study visualizations, we generated structures with ESM3[21] in sequence-conditioned mode and decoded the structure-track outputs to Cartesian coordinates using the provided utilities. We did not fine-tune or modify ESM3. ESM3 outputs were treated as fixed input structures for feature computation and visualization.

**LigCys-SA models.** LigCys-SA augments LigCys-seq with residue-level structural features computed from ESM3-generated coordinates. We generated solvent-excluded surface meshes with MSMS[22] (probe radius 1.4 Å) and derived features capturing solvent accessibility/topology (FreeSASA[23]), geometric binding-site proximity (distance to LigBind-seq–predicted binding residues), structural confidence (pLDDT), and local chemical environment (KaML-ESM p$K_a$ metrics; RIDAO disorder metrics). Features were clipped, log-transformed where appropriate, and zero-imputed, yielding $\sim$70 candidates; after removing near-zero-variance features and pruning correlated pairs ($|r| > 0.9$; z-scored), 45 nonredundant features remained (Supplementary Table S4) and were concatenated with frozen 2,560-dimensional ESMC embeddings for training. To combine frozen ESMC embeddings with engineered structural features, we used a disjoint adaptive gating unit (DAGU) that learns feature-wise gates to weight and combine the ESMC and structural-feature blocks. The concatenated input was split into an ESMC block and a structural-feature block; each block was layer-normalized and passed through a lightweight learned gate (linear layer followed by sigmoid) that outputs a same-dimensional mask over features. The normalized features are modulated by element-wise multiplication with their masks; the gated blocks are concatenated and jointly normalized, and the result is passed to the LigCys-seq task head.

**LigBind-SA models.** LigBind-SA augments LigBind-seq with residue-level structural embeddings computed from ESM3-generated coordinates. Structural embeddings were produced by an ensemble of eight pretrained geometric transformers (PeSTo)[24] and pooled to residue-level vectors. These embeddings were concatenated with frozen 2,560-dimensional ESMC representations and passed to the LigBind-seq prediction head (one-layer MLP; GELU; dropout $p = 0.1$); the DAGU block was not used. Training and ensembling follow LigBind-seq. LigBind-SA was evaluated only as an ablation and was not used in the production pipeline because it requires generated structures at inference time.

#### 4.3 Ensembling and ranked recovery

To improve prediction stability and generalization, all models were evaluated using ensemble inference. All ensembles used leakage-free partitions defined by LatentLift (Section 3.1), enforcing cluster-level indivisibility and UID exclusivity. For LigCys, proteins in LC3D, LC3Dts, and the ABPP hold-out set were excluded from all training and validation splits; LC3D was used for dataset expansion (Section 5.1), whereas LC3Dts and the ABPP hold-out set were used strictly for evaluation. For LigBind and the functional context models, leakage-free splits were enforced within their respective structure-derived datasets using the same cluster-indivisibility and UID-exclusive policy.

For each task (LigCys, LigBind, and the functional context annotation models) and model variant (seq and SA, where applicable), we constructed 10 distinct data splits and trained 20 independent models per split, yielding 200 trained models per setting. All replicates used identical configurations but distinct random seeds and training shuffles. As a diagnostic, Supplementary Data SD10 summarizes, for each split, the performance of individual replicates (mean $\pm$ s.d. across random seeds) alongside the corresponding 200-member ensemble (10 splits $\times$ 20 replicates).

To form an ensemble, we selected a single checkpoint from each replicate corresponding to the epoch with the highest validation AUPRC. These 200 best-performing models were then used to generate per-site predictions for evaluation and ranked recovery (see *Model evaluation and ranked recovery*).

For LigBind models and functional context models, ensemble predictions were aggregated by averaging predicted probabilities. For LigBind, evaluation included all residue types. For the functional context models, which annotate LigCys predictions, evaluation was restricted to cysteines.

**LigCys reporting: Top-1 voting during development and CGRV for final results.** In this subsection, “Top-1” refers to the ensemble voting scheme, whereas “Top-1 recovery” refers to the ranked-retrieval evaluation metric defined below. During model development (benchmarking, ablations, and dataset expansion), we summarized LigCys ensemble behavior using a Top-1 voting scheme: for each protein, each ensemble member cast one Top-1 vote for the cysteine with the highest ligandability probability. For each cysteine, the fraction of ensemble members voting for that site (its Top-1 vote fraction) served as the **LigCys score** under this scheme. However, because the LigCys decision rule used throughout this work is that a cysteine is predicted to be ligandable if and only if its LigCys score is nonzero, Top-1 voting has two coupled limitations. First, isolated Top-1 nominations from a small number of outlier models can create false-positives even when the remaining models provide no support. Second, Top-1 voting does not capture consistent near-miss sites that are repeatedly ranked just below the winner (e.g., consistently rank-2) but rarely win Top-1; such sites can also exhibit low Top-1 vote fractions that are difficult to distinguish from true outliers.

To address these coupled failure modes in the production pipeline, we developed **consensus-gated residual voting (CGRV)** for LigCys reporting. Unless otherwise stated, all reported LigCys results in this work (including main-text results from the production model) use CGRV rather than Top-1 voting. CGRV operates per protein using per-cysteine probabilities and is designed to suppress outlier-driven nominations while retaining consistent runner-up signal. To start, each ensemble member casts a rank-1 vote for its highest-probability cysteine. When one cysteine is a dominant rank-1 winner (receiving at least 50% of rank-1 votes), dissenting rank-1 votes are treated as spurious and are reassigned to the dominant winner.

To improve robustness to near-miss consensus, CGRV also allows limited additional voting for runner-up sites. A site is eligible for rank-2 voting only if it is rank-2 by at least 20% of ensemble members and does not primarily track overall model confidence on that protein (defined by an $R^2 \leq 0.60$ when regressing the site's probabilities across ensemble members against each member's maximum per-protein probability). For each ensemble member, among eligible sites we selected the one with the largest positive residual from this regression and cast one rank-2 vote if that residual was $> 0$ (otherwise the model abstained). Rank-3 voting was analogous, requiring that a site be exactly rank-3 in at least 40% of ensemble members and excluding the primary winner and the global rank-2 winner. We used stricter consensus requirements for lower-ranked votes because ensemble agreement typically weakens beyond the top prediction; tightening these thresholds reduces outlier-driven secondary and tertiary nominations and helps control false positives.

Total CGRV support was summarized as the total number of votes a site received across rank-1, rank-2, and rank-3 stages, divided by the number of ensemble members; we refer to this quantity as the **LigCys score**. Because a single ensemble member may contribute votes in multiple stages, LigCys scores can exceed 1. Under the LigCys decision rule used throughout this work, any cysteine with a nonzero LigCys score is predicted to be ligandable.

**Confidence score for LigCys outputs.** In addition to the LigCys score, we report a confidence score derived from false-discovery control[25] over LigCys nominations. Conceptually, we estimate an empirical null distribution for each protein by breaking the correspondence between residues and ensemble-member predictions within that protein, compute per-cysteine significance values, and convert them to $q$-values using Benjamini–Hochberg correction[25] across cysteines in the same protein. For nominated cysteines, we report confidence as $\mathrm{Conf} = 1 - q$, so higher confidence indicates a lower expected false-discovery rate; cysteines that are not nominated are assigned $\mathrm{Conf} = 0$. Confidence is reported post hoc and does not affect model training, ensemble construction, or which sites are nominated.

**Model evaluation and ranked recovery.** Unless otherwise stated, all performance metrics were computed from ensemble outputs. For LigCys, the LigCys score served as the continuous ensemble score. Models were evaluated on held-out sets using standard classification metrics, including area under the receiver operating characteristic curve (AUROC) and area under the precision–recall curve (AUPRC). To provide an interpretable breakdown of prediction behavior, we computed per-protein precision and recall at a single fixed decision threshold (per model setting) and averaged these quantities across proteins. For LigCys, predictions were binarized as positive if the LigCys score was $> 0$ and negative otherwise. For probability-based models, binary predictions were obtained by thresholding the ensemble-averaged residue probability using a fixed evaluation threshold derived from validation data. For each ensemble member, we selected the checkpoint with the highest validation AUPRC, identified the probability threshold that maximized validation F1 for that checkpoint, and defined the ensemble evaluation threshold as the mean of these per-member thresholds. The resulting fixed evaluation thresholds for LigBind and the functional context models are reported in Supplementary Data SD14.

We also evaluated ranked retrieval performance using top-$k$ recovery. For each test protein, residues were ranked by the ensemble score (LigCys score for LigCys; averaged probability for LigBind and context models). Given a set of $N$ proteins and ligandable residues $L_i$ for protein $i$ ($1 \leq i \leq N$), and the set of residues with top-$k$ predictions $R_{k,i}$, top-$k$ recovery was defined as the fraction of recoverable true positives that appear in the top-$k$ predictions:

$$\text{Top-}k \text{ Recovery} = \frac{1}{M_k} \sum_{i=1}^{N} |L_i \cap R_{k,i}| \,,$$

where

$$M_k = \sum_{i=1}^{N} \min \{k, |L_i|\}$$

is the maximum number of recoverable positives at threshold $k$. For $k > 1$, this formulation is equivalent to a per-protein average weighted by $\min\{k, |L_i|\}$ (proteins with more recoverable sites contribute proportionally more); for $k = 1$, $\min\{1, |L_i|\} \in \{0, 1\}$, so Top-1 recovery reduces to the mean hit rate across proteins with at least one true positive.

We emphasize Top-1 recovery (i.e., $k = 1$) as the primary evaluation metric, reflecting practical constraints in experimental screening where only a single site can be tested. Top-1 recovery directly answers the operational question: when limited to testing one predicted site per protein, how often is the model's top recommendation truly ligandable?

Ranked recovery is also more robust under the labeling assumptions used in this work because negative labels in our benchmark sets are provisional—they reflect the absence of observed ligandability rather than confirmed inactivity. This applies both to structure-derived negatives (i.e., residues without observed ligand binding in available structures) and to ABPP-derived negatives (i.e., sites that are unliganded or undetected in the ABPP sources considered). As a result, false-positive counts (and derived metrics such as precision) should be interpreted cautiously. Ranked recovery instead emphasizes whether experimentally validated positives are prioritized near the top of each protein's ranking, aligning with the platform's goal of guiding residue selection for follow-up validation. Accordingly, Top-1 recovery is reported as the primary evaluation criterion across all benchmark comparisons.

All classification metrics (e.g., AUROC, AUPRC, precision, recall, and F1 score) were computed per protein and then averaged across proteins. This formulation is more stringent than global aggregation, as it mitigates bias from protein length or site count and emphasizes consistency across targets. Reporting per-protein classification metrics therefore provides a more realistic estimate of the performance an end user can expect when applying the model to individual proteins in proteome-wide screens.

### 5 Training-data expansion for LigCys

We developed two iterative training-data expansion procedures that differ in their acceptance criterion. The LC3D-guided procedure (Section 5.1) defines the LigCys-S training set by accepting batches that improve LC3D Top-1 recovery, whereas the ABPP-guided procedure (Section 5.2) defines the LigCys-A training set by accepting batches that improve cross-validated ABPP AUPRC. In both procedures, candidate cysteines were drawn from the ABPP-derived data after LatentLift clustering and consensus labeling (Section 3.1). Throughout expansion, LC3D proteins and the ABPP hold-out proteins remained excluded from the training pool, preserving LatentLift-defined leakage-free evaluation partitions.

#### 5.1 LC3D-guided expansion

To increase training-set size and coverage while preserving high-confidence supervision, we applied an iterative expansion procedure guided by LC3D Top-1 recovery. This LC3D-guided procedure underlies the LigCys-S training set reported in Supplementary Table S3, which is approximately two- to threefold larger than the distilled 4S–4R core in labeled cysteines and proteins.

The protocol started from the truncated 4S–4R dataset (one representative cysteine per labeled LatentLift cluster; Section 3.1), restricted to proteins with both positive and negative cysteines.

**The candidate pool.** The candidate pool included cysteines meeting the same 4S–4R consensus threshold, drawn from (i) proteins containing exclusively positive cysteines, (ii) proteins containing exclusively negative cysteines, and (iii) non-representative cysteines within each cluster.

**Expansion protocol.** In each iteration, we randomly sampled 100 candidate batches, each containing 175 cysteines (except iterations 1–3, which used batch sizes of 80, 100, and 150 cysteines, respectively). Within each batch, cysteines were randomly split into training and validation subsets (9:1). For each batch, we trained 24 models (6 random splits $\times$ 4 random seeds) and selected checkpoints by validation AUPRC. These reduced-size ensembles were used only for batch selection during expansion; final reported benchmarking uses the 200-member ensembles described in Section 4.3. Throughout expansion, LigCys scores were computed using the Top-1 voting scheme (Section 4.3) rather than CGRV. The resulting 24-model ensemble was evaluated on the LC3D benchmark using Top-1 recovery (Section 4.3), and the batch achieving the highest LC3D Top-1 recovery was accepted. Because LC3D Top-1 recovery is the acceptance criterion, LC3D serves as a development benchmark during this expansion; generalization was additionally monitored on the ABPP hold-out set described in Section 3.1, which was never used for batch acceptance or model selection. All cysteines from the accepted batch (with their labels) were then transferred from the candidate pool to the training pool. Iterations continued until LC3D Top-1 recovery no longer increased. The final expanded set (4S–4R-S) was used to train a production model; the ABPP hold-out set was used only for evaluation. The production model was then benchmarked on LC3D, and performance across iterations is reported in Supplementary Table S3.

#### 5.2 ABPP-guided expansion

This ABPP-guided expansion procedure underlies the LigCys-A training set reported in Supplementary Table S3. As an alternative to LC3D-guided expansion, we expanded training data based on cross-validated ABPP AUPRC. We started from the same LatentLift-labeled 4S–4R core dataset (one representative cysteine per labeled LatentLift cluster; Section 3.1), restricted to proteins with both positive and negative cysteines.

**Batch construction.** Candidate batches were assembled with exactly 25 positives and the remainder negatives. During early iterations, the batch size was 100 cysteines (25 positives, 75 negatives); as the pool expanded, the batch size increased to 175 cysteines (25 positives, 150 negatives). Cysteines were drawn from the candidate set in order of highest ensemble uncertainty, defined as ascending $|\,\overline{p}-0.5\,|$ (closest to 0.5 first), where $\overline{p}$ is the mean predicted ligandability probability across ensemble members at the selected checkpoints, so that the most uncertain cysteines were evaluated earliest.

**Cross-validation.** We used group-aware evaluation over $K = 10$ prespecified UID-based train/validation splits, each formed by holding out 20% of UIDs for validation and using the remaining 80% for training; the splits were selected to minimize overlap between validation UID sets (low pairwise Jaccard similarity). For each split, we trained three replicate models (different random seeds), for a total of $K \times R = 30$ models for each baseline and candidate evaluation. As above, these ensembles were used for tournament scoring; final reported benchmarking uses the 200-member ensembles described in Section 4.3. The UID-to-split mapping was generated once at the start of each tournament, with each validation subset constrained to match the global positive fraction within 1%. This mapping was cached and reused throughout the tournament to minimize variance from resampling. At the start of each iteration, a baseline ensemble was trained on the current training pool using the cached splits; its checkpoints provided (i) a reference for scoring candidate batches and (ii) per-cysteine ensemble uncertainty scores used to prioritize cysteines during batch construction.

**Controlling variance.** Several safeguards ensured that observed performance changes reflected the added cysteines rather than randomness. A single UID-to-split mapping was generated at the start of each tournament and cached for all evaluations within that tournament; validation subsets were constrained to match the global positive fraction within 1%. After each acceptance step, newly added UIDs were deterministically assigned to splits to maintain balance while preserving existing assignments. To reduce long-term partitioning bias, the tournament was periodically restarted with freshly generated splits (after iterations 4, 8, and 12). All models within an iteration (baseline and candidates) were initialized from the same saved weight snapshot and trained with identical architectures, losses, and optimization schedules (see Section 4.1). Finally, three random-seed replicates were trained per split, and performance was summarized as the mean across all $K \times R$ split–replicate scores, with a 95% confidence interval computed over those values. Together, these design choices minimized variance from partitioning and training stochasticity, isolating performance gains attributable to the added cysteines.

**Tournament evaluation.** For each candidate batch, we formed a provisional augmented training pool by unmasking the corresponding cysteines in the candidate pool, and evaluated performance on the same cached splits used for the iteration's baseline ensemble. Performance was measured as AUPRC enrichment, $\Delta = \mathrm{AUPRC} - \mathrm{prevalence}$, which quantifies improvement over the random baseline expected at the observed class balance and prevents shifts in prevalence from spuriously changing apparent performance. Each batch was scored by the mean across all $K \times R$ split–replicate scores with a 95% confidence interval (CI) computed over those values; batches were ranked by the lower CI bound.

**Recombination and selection.** Each tournament iteration comprised up to four rounds. In round 1, all candidate batches were scored against the baseline ensemble. In subsequent rounds, we applied a hybrid genetic-algorithm and successive-halving strategy: the top fraction of batches (20% in early iterations; 50% in later iterations) was retained and the remainder discarded. Retained batches were recombined via one-point crossover (swapping segments of cysteines between two parent batches) and mutation (randomly altering a small fraction of cysteines to maintain diversity), with an exploration fraction $\epsilon = 0.1$ controlling the mutation rate. The best-performing batch from the previous round was always preserved.

**Acceptance criteria and pool updates.** At the end of each tournament iteration, the overall best-performing batch was accepted if its lower CI bound or its mean enrichment exceeded the baseline ensemble. Accepted cysteines were merged into the training pool (updating existing UIDs or creating new records as needed) and masked in the candidate pool to prevent reselection. For each iteration, a detailed acceptance log was recorded and both checkpointed and augmented training pools were saved. The process terminated when no candidate batch met the acceptance threshold.

### 6 Benchmarking LigCys against prior methods

To contextualize LigCys predictions relative to prior structure-based cysteine ligandability predictors, we benchmarked LigCys against TopCysPAL (https://topcysteinedb.hhu.

de/Prediction) and CovCysPred using the strictly external, time-stamped co-crystal test set LC3Dts (Section 1.2). LC3Dts comprises wwPDB structures with covalently liganded cysteines deposited between January 1, 2024 and November 11, 2025 and was constructed after model development; it was not used for training, hyperparameter tuning, or model selection. LC3Dts contains 69 unique sequences spanning 68 proteins because one protein is represented by two distinct, non-overlapping PDB constructs that each contain a covalently liganded cysteine. Throughout this benchmarking, the unit of evaluation is the PDB chain (UID `PDBID-CHAINID`).

Both TopCysPAL and CovCysPred were trained on structure-derived features and require structures as input, whereas LigCys is sequence-only. Accordingly, to provide a best-case comparison for the structure-based baselines, we supplied each method with experimentally resolved wwPDB coordinate files (i.e., real PDB structures rather than predicted structures), which is a favorable setting for these baselines.

TopCysPAL predictions were obtained by querying the public TopCysPAL web application for each LC3Dts entry by providing the corresponding PDB identifier and chain ID. We note that TopCysPAL reports a wwPDB snapshot date of June 26, 2024 for its training-set construction; consequently, 25 LC3Dts evaluated chains (UID `PDBID-CHAINID`, each corresponding to a unique UniProt entry) were deposited between January 1, 2024 and June 26, 2024 and thus could overlap with TopCysPAL training data. LC3Dts entries deposited after June 26, 2024 are necessarily out-of-snapshot.

CovCysPred predictions were generated locally using the authors' implementation (https://github.com/BrynMarieR/CovCysPredictor). Coordinate files for LC3Dts entries were fetched directly from the wwPDB and processed under the authors' recommended settings. For CovCysPred, the available documentation does not specify the wwPDB snapshot date used to assemble their structure dataset; therefore, we cannot directly quantify training-set overlap with LC3Dts, although overlap is plausible given the publication timeframe.

Benchmark labels were taken from LC3Dts: cysteines observed to be covalently liganded in the curated co-crystal structures were treated as positives. As in other structure-derived datasets used in this work, remaining cysteines in each evaluated chain are weakly labeled negatives, reflecting the absence of observed covalent modification in the available structural context rather than definitive evidence of non-ligandability. To directly compare methods in a practical experimental prioritization setting under weakly labeled negatives, we emphasize ranking-based evaluation. LigCys scores were generated by the final production ensemble using CGRV (Section 4.3).

For each method, we report (i) Top-1 recovery on LC3Dts, defined per protein chain as whether the highest-scoring cysteine above the method cutoff (TopCysPAL: 0.5; Cov-CysPred: 0.14) is an LC3Dts `POS` site, and (ii) AUROC and AUPRC computed from each method's continuous scores against the LC3Dts labels. If a method returned no cysteine above its cutoff for a chain, that chain was counted as a miss for Top-1 recovery. All benchmarking results and per-site prediction discrepancies are provided in Supplementary Data file 2.

### 7 External modules and annotations

All tools in this section were used only as auxiliary annotators to generate engineered inputs for the structure-aware LigCys-SA ablations (Section 4.2) and to stratify analyses; they were not used for supervision, LatentLift clustering/partitioning, or inference in the production sequence-only models.

**RIDAO.** The Rapid Intrinsic Disorder Analysis Online (RIDAO) platform aggregates six intrinsic-disorder predictors to produce per-residue annotations of intrinsic disorder and molecular recognition features (MoRFs).[26] For each input sequence, RIDAO reports per-residue predictions from its constituent tools and a mean disorder profile. Additionally,

RIDAO provides ANCHOR2[27] predictions of MoRF propensity. MoRFs are short disordered segments that can undergo binding-induced folding.

RIDAO outputs are used in two places in this work: (i) as sequence-derived auxiliary inputs for disorder/MoRF feature construction for the structure-aware LigCys-SA model (Section 4.2), and (ii) to stratify and report LigCys performance by disorder class (Supplementary Fig. S11).

**KaML-ESM.** KaML-ESM is a sequence-only model for residue-level p$K_a$ prediction. It uses per-token embeddings from protein language models (ESM2 or ESMC) as inputs to a small feedforward task head to predict residue-specific p$K_a$ values for titratable amino acids (Asp, Glu, His, Lys, Cys, and Tyr).

KaML-ESM was used only to generate engineered electrostatics features for the structure-aware LigCys-SA ablations (Section 4.2). Within AiPP, it is run in inference mode as an auxiliary, sequence-derived annotator to obtain per-cysteine predicted p$K_a$ values and derived quantities, including cysteine p$K_a$ and p$K_a$ shift relative to the solution model value.

#### Supplementary Figures

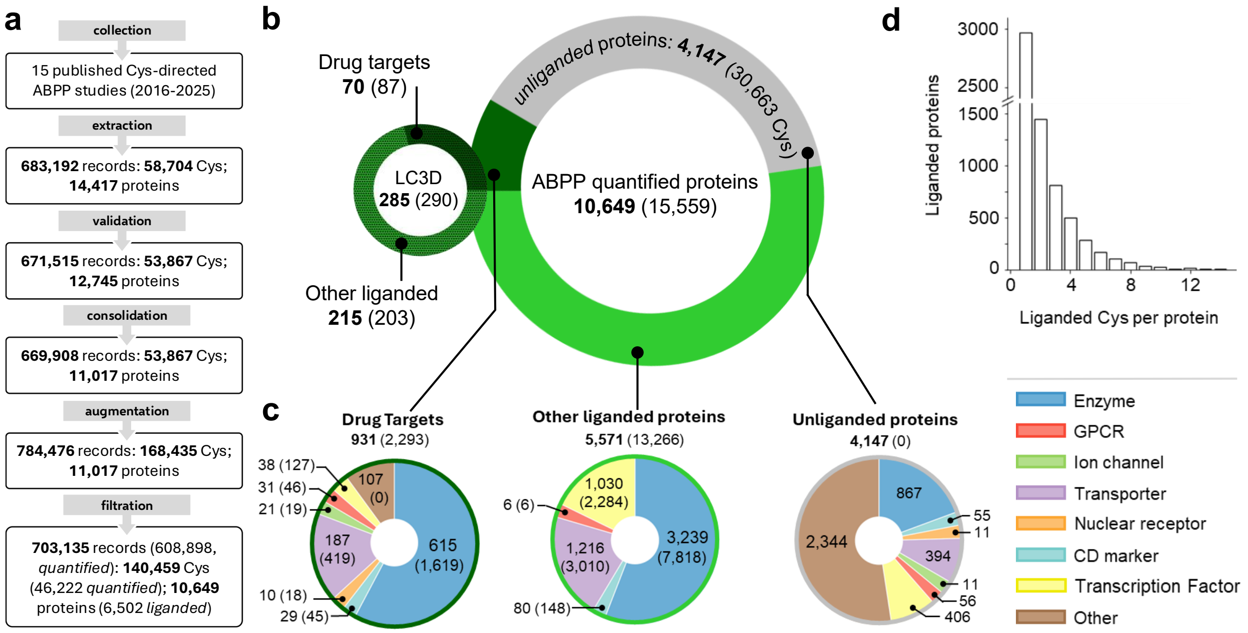

Figure S1: **Overview of the ABPP-quantified proteins and cysteines in LigCysABPP database. a.** The LigCysABPP database is manually curated from a collection of 15 published cysteine-directed ABPP studies.[1–15] Peptide-derived records are extracted from each publication, validated against UniProt, consolidated across identical sequences and subsequences, augmented with unquantified (unseen) cysteine-sites on otherwise quantified proteins, and filtered for compatibility with downstream processing steps. **b.** LigCysABPP contains the cysteine ligandability records of 10,649 ABPP-quantified proteins, of which 6,502 are liganded and 4,147 are unliganded. A liganded cysteine is defined as having at least 1 pos ABPP record while a liganded protein is defined as having at least 1 liganded cysteine. Among the liganded proteins, 931 are known drug targets (with 2,293 liganded cysteines), while the rest of 5,571 other liganded proteins contain 13,266 liganded cysteines. The LC3D database contains 285 unique proteins (290 cysteines liganded in the co-crystal structures), among which only 70 proteins (87 drug targets) are also identified as liganded by ABPP. **c.** Functional classes of drug targets, other liganded proteins, and unliganded proteins according to The Human Protein Atlas classifications[28](https://www.proteinatlas.org/). **d.** Histogram of the number of ABPP liganded cysteines per protein shows that most liganded proteins contain 1-3 liganded cysteines.

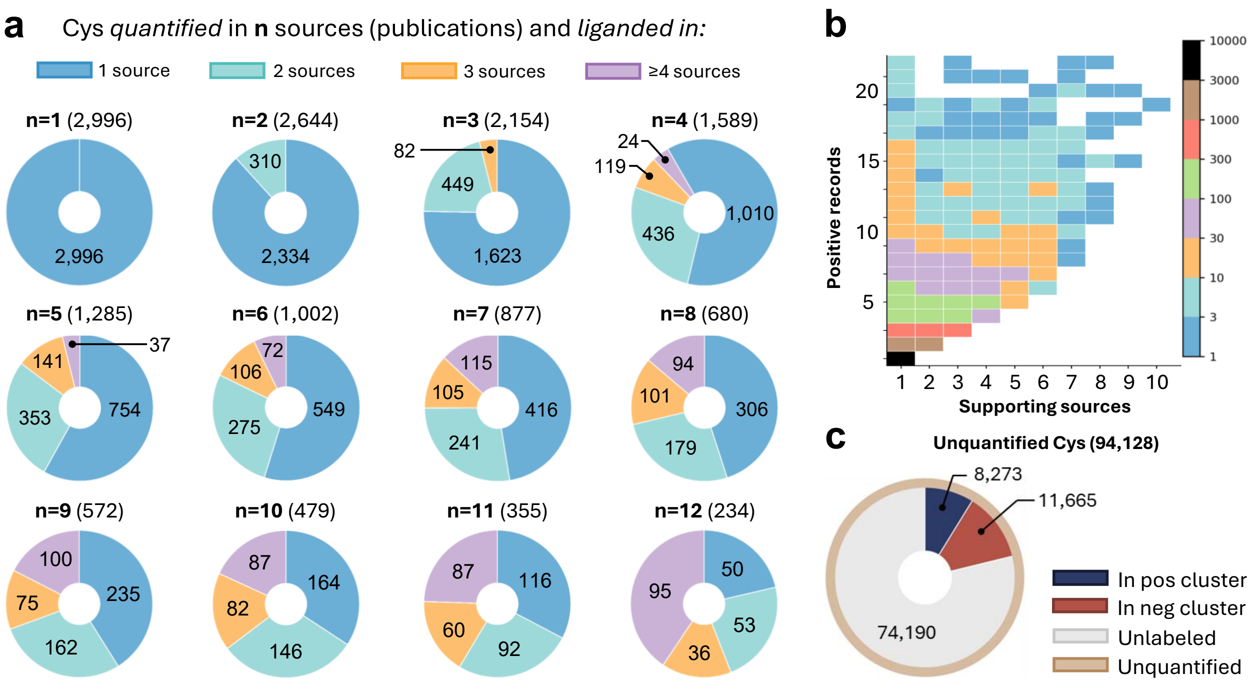

Figure S2: **Analysis of cysteine ligandability across ABPP sources and records. a.** Each pie chart displays the number of liganded cysteines quantified by $n$ sources. A liganded cysteine is defined as one that has at least one pos ABPP record. Each pie segment represents the number of cysteines supported by 1 (blue), 2 (green), 3 (yellow), and $\geq$4 (purple) sources. **b.** The number of liganded cysteines with the number of supporting sources and positive records. A supporting source for a specific cysteine is defined as one that contains at least one pos record for that cysteine. A total of 7,787 cysteines are labeled pos by only 1 record (black). No liganded cysteine is supported by $>$10 sources. Liganded cysteines with $>$25 records are negligible and therefore excluded from the plot. **c.** Following LatentLift clustering (Supplementary Methods), 8,273 unquantified cysteines are labeled pos, 11,665 labeled neg, while 74,190 cysteines remain unlabeled.

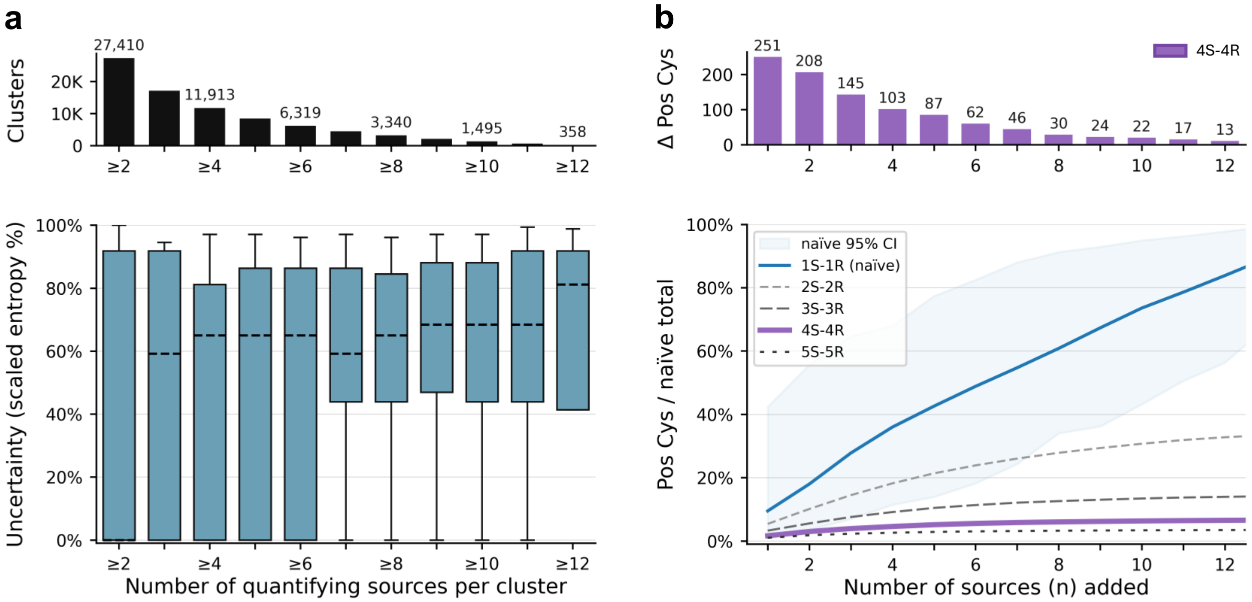

Figure S3: **Analysis of how source threshold affects label variability and data coverage. a.** *Top:* Bar plot showing the number of cysteine clusters supported by at least $n$ independent sources, regardless of the number of underlying records. For clarity, bins corresponding to odd values of $n$ are unlabeled. *Bottom:* Label variability within each cluster, stratified by the number of supporting sources. Variability is quantified using the Shannon entropy of the binary label distribution ($p$, the fraction of sources labeling a cysteine as positive), scaled to a 0–100% range. Boxes denote the interquartile range; medians are shown as horizontal dashed lines; whiskers span the 10th–90th percentiles. A minimum in entropy is observed at $n \geq 4$, suggesting this threshold maximizes labeling consistency across sources. **b.** *Top:* Incremental gain in the number of positive cysteines when increasing from $n-1$ to $n$ sources. *Bottom:* For each consensus threshold (e.g., 1S–1R through 5S–5R), we randomly sample $n$ sources from a pool of 15, repeat 200 times, and compute the average number of positive cysteines. This is normalized by the naïve total (i.e., all cysteines reported as positive in any single record). The 95% confidence interval is shown for the naïve curve. The 4S–4R consensus curve (highlighted in purple) plateaus after four sources, supporting its use as a data-driven threshold for robust labeling.

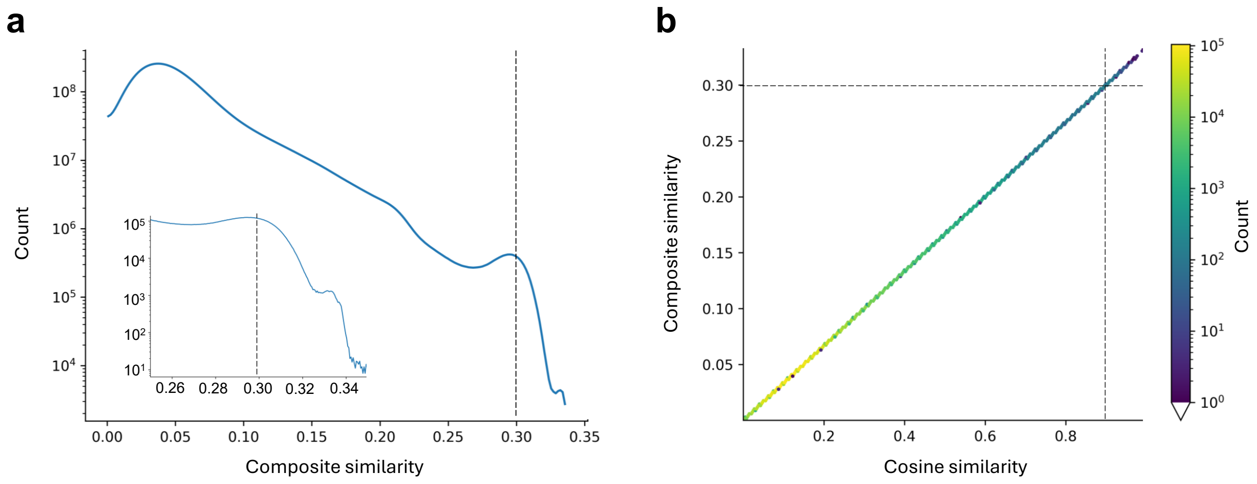

Figure S4: **Similarity metrics used for LatentLift clustering. a.** Distribution of the composite similarity score $S = \frac{1}{3}(C + D_1 + D_2)$, where $C$ is cosine similarity and $D_1 = 1/(1+L_1)$ and $D_2 = 1/(1+L_2)$ are inverse $L_1/L_2$ embedding distances. Counts are shown on a log scale. The LatentLift clustering cutoff $S = 0.3$ is indicated (dashed line); inset shows a zoom around the cutoff. **b.** Joint distribution of composite similarity $S$ and cosine similarity $C$ (hexbin density; colour indicates pair counts on a log scale), estimated from a random sample of $n = 5{,}000{,}000$ pairs. Dashed lines mark $S = 0.3$ and the corresponding cosine-similarity value obtained by linear mapping between $S$ and $C$ in the sampled pairs.

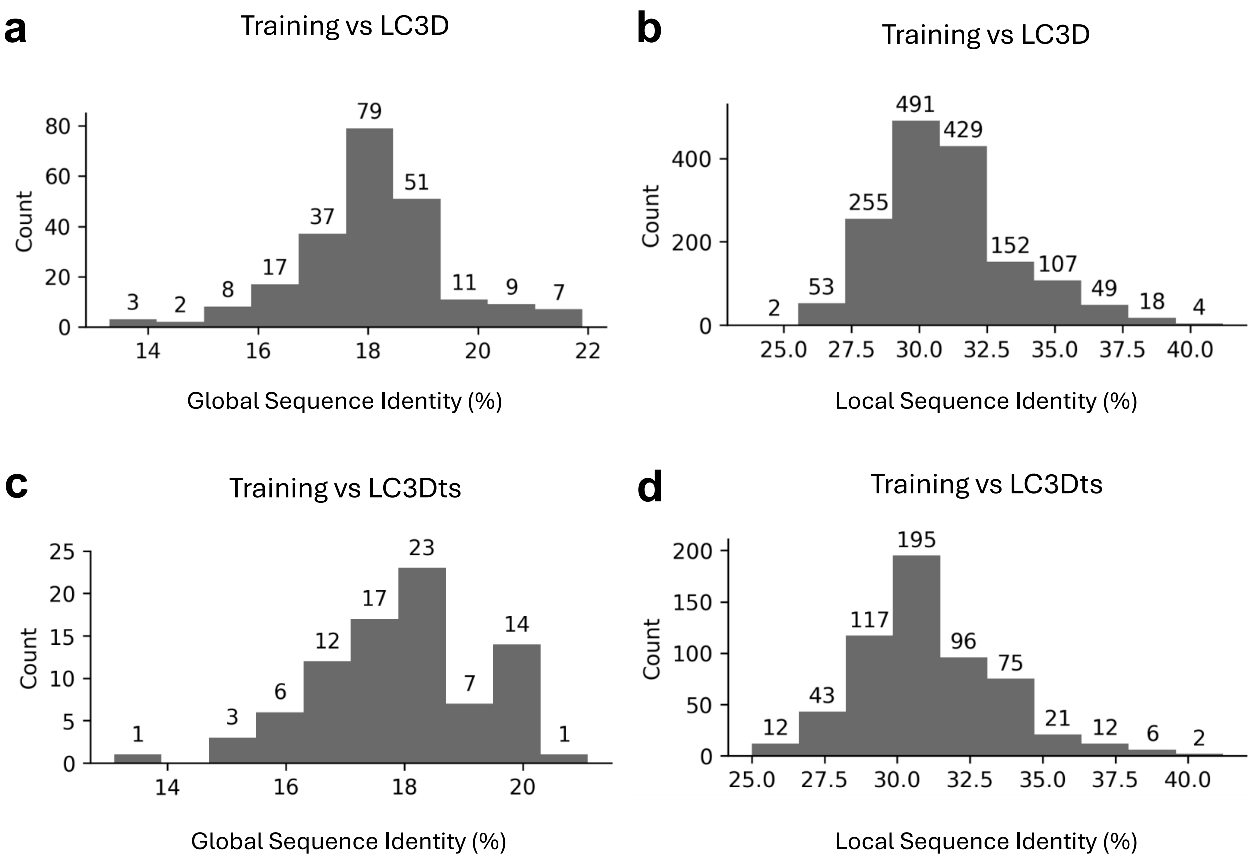

Figure S5: **Sequence identities between the LigCys training set and the LC3D benchmark and LC3Dts external test set. a,c.** Protein sequence identities between proteins in the training set and in the LC3D (a) or LC3Dts (c) test set. For each test-set protein, we computed the maximum global sequence identity to any training-set protein using EMBOSS[29] and plotted the histogram for all proteins. The maximal sequence identity is 21.9% and 21.2% for LC3D and LC3Dts, respectively. **b,d.** Local sequence identities around the cysteine of interest between the training set and LC3D (b) or LC3Dts (d) test set. For each cysteine of interest in the test set, we extracted 30 amino acids centered on that cysteine (excluding sites within 15 residues of either terminus) and computed the maximum identity of this sequence to the corresponding one in the training set using EMBOSS.[29] The distribution shows a maximal identity of 41.2% for both test sets.

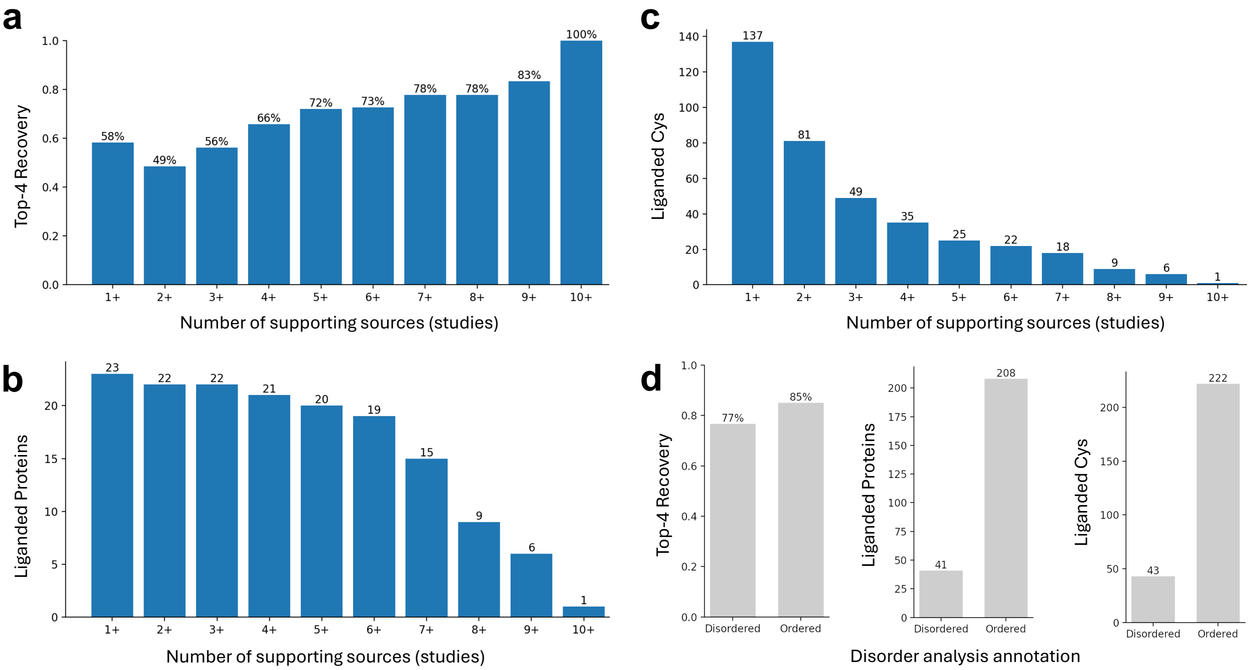

Figure S6: **Evaluation of the production LigCys task head on the ABPP hold-out set across source support levels and stratified analysis by structure disorder. a-c.** The blended, production LigCys ensemble was evaluated on an ABPP hold-out set of 23 proteins quantified by $\geq$10 independent ABPP sources, each containing at least one liganded and one unliganded cysteine. Positive support denotes the number of independent ABPP sources reporting a liganding event for a given cysteine site. At $t+$ supporting sources (threshold), sites with positive support $\geq t$ were treated as liganded. **a.** Top-4 recovery of liganded sites at each threshold, computed from per-protein LigCys score rankings (Supplementary Methods). **b.** Number of proteins with at least one liganded cysteine meeting the threshold. **c.** Number of liganded cysteines meeting the threshold. **d.** Disorder-stratified Top-4 recovery analysis and the counts of liganded proteins and liganded cysteines for each category. To increase data coverage, ABPP hold-out (at 4+ support level) was combined with the LC3D benchmark set. Cysteines were assigned to intrinsically disordered or ordered regions using the external module RIDAO (Supplementary Methods; a residue was labeled disordered if any predictor in RIDAO classified it as disordered).

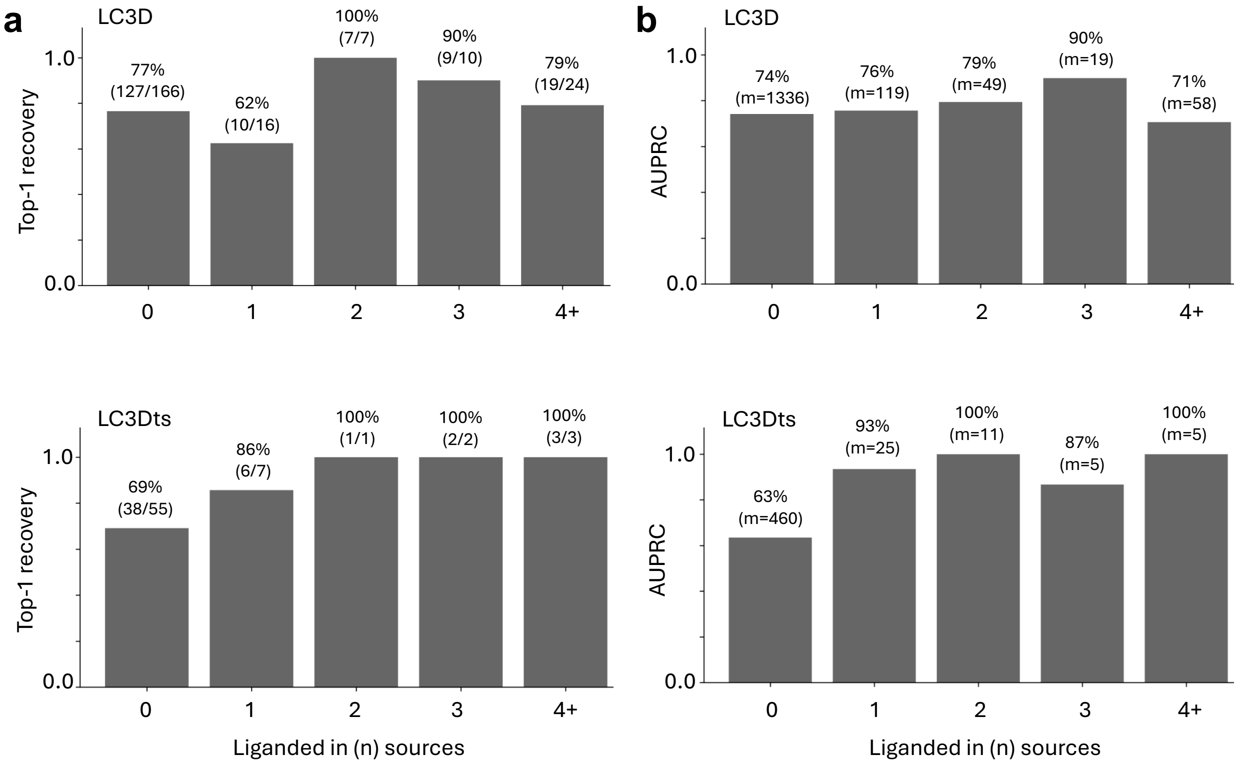

Figure S7: **A *post hoc* analysis of LigCys performance vs. number of supporting ABPP sources not in the LigCys training set. a.** Top-1 recovery on LC3D (top) and LC3Dts (bottom) stratified by the number of independent ABPP studies supporting the top-ranked site for each protein. For each cysteine in the respective set, we count the number of distinct ABPP sources contributing at least one positive record to its LatentLift cluster. Numbers above bars show the Top-1 recovery and the number of correct Top-1 hits over recoverable proteins in each bin. **b.** Site-level AUPRC for LC3D (top) and LC3Dts (bottom) stratified by the same ABPP source-support bins, computed over all cysteines in each bin; numbers above bars show AUPRC and the number of unique cysteine sites ($m$).

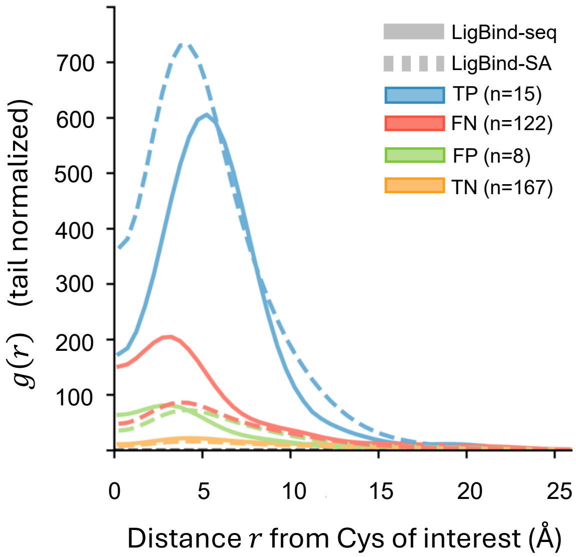

Figure S8: **LigBind predictions can rescue false negatives and filter false positives in cysteine ligandability assessment.** Radial distribution function (RDF) of LigBind-predicted reversible binding residues around cysteine sites, stratified by the LigCys prediction outcome (TP, TN, FP, FN). Solid and dashed lines represent the LigBind-seq and LigBind-SA predictions, respectively, on the pruned LC3D benchmark set that excludes proteins in the LigBind training set. Distance is the minimum all-atom Euclidean distance between the cysteine of interest and each predicted ligand-binding residue. RDF is normalized by the mean shell density over the last 20% of the distance range.

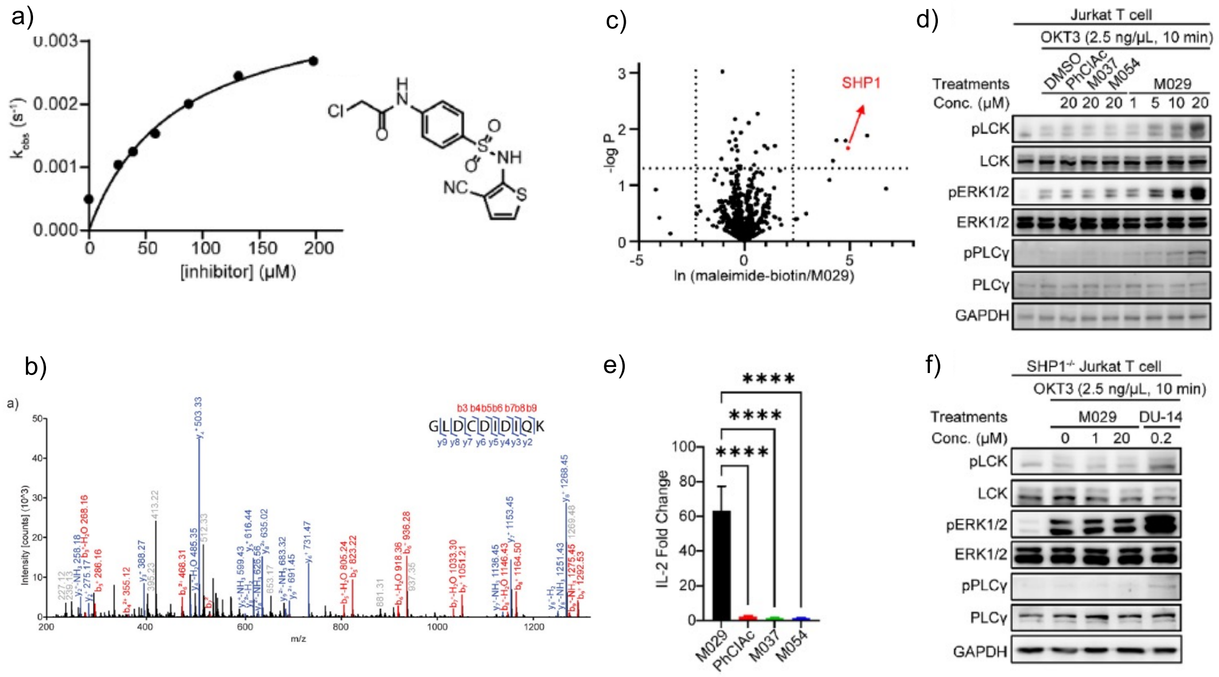

Figure S9: **Characterization and efficacy of the covalent allosteric inhibitor of SHP-1. a, b.** Characterization of the covalent allosteric inhibitor (**M029**) of SHP-1. **c-f. M029** engaged SHP-1 in cellulo and showed strong efficacy in immune activation. Adapted from Ref.[30]

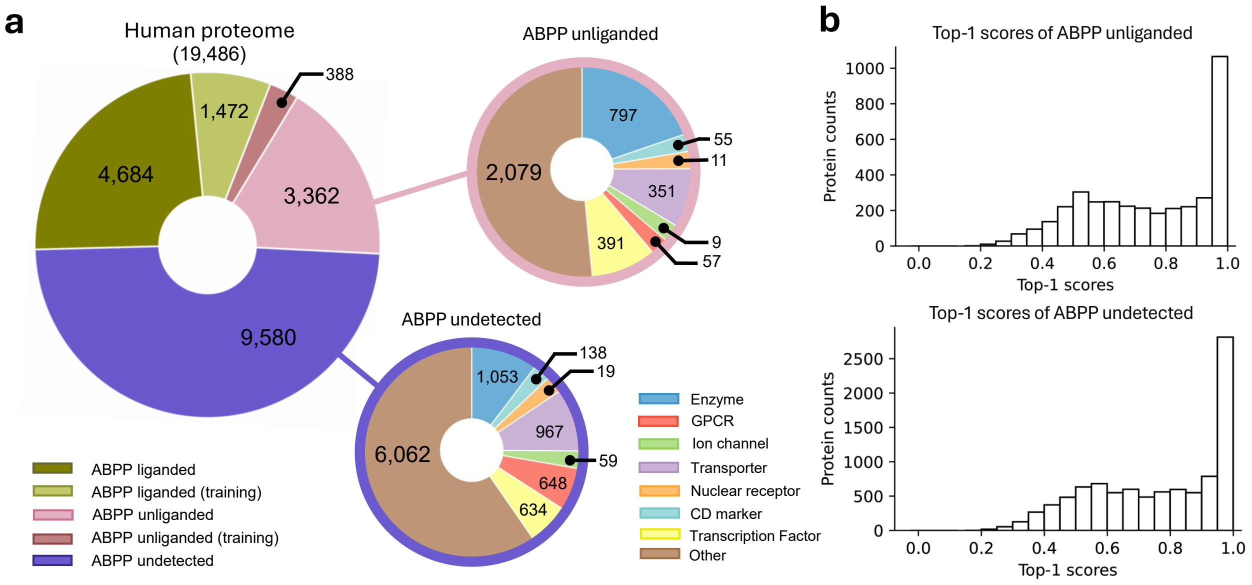

Figure S10: Functional family classification and LigCys Top-1 prediction scores for the ABPP-unliganded and ABPP-undetected proteins in the human proteome that were excluded from model training. This figure is to accompany Figure 6.

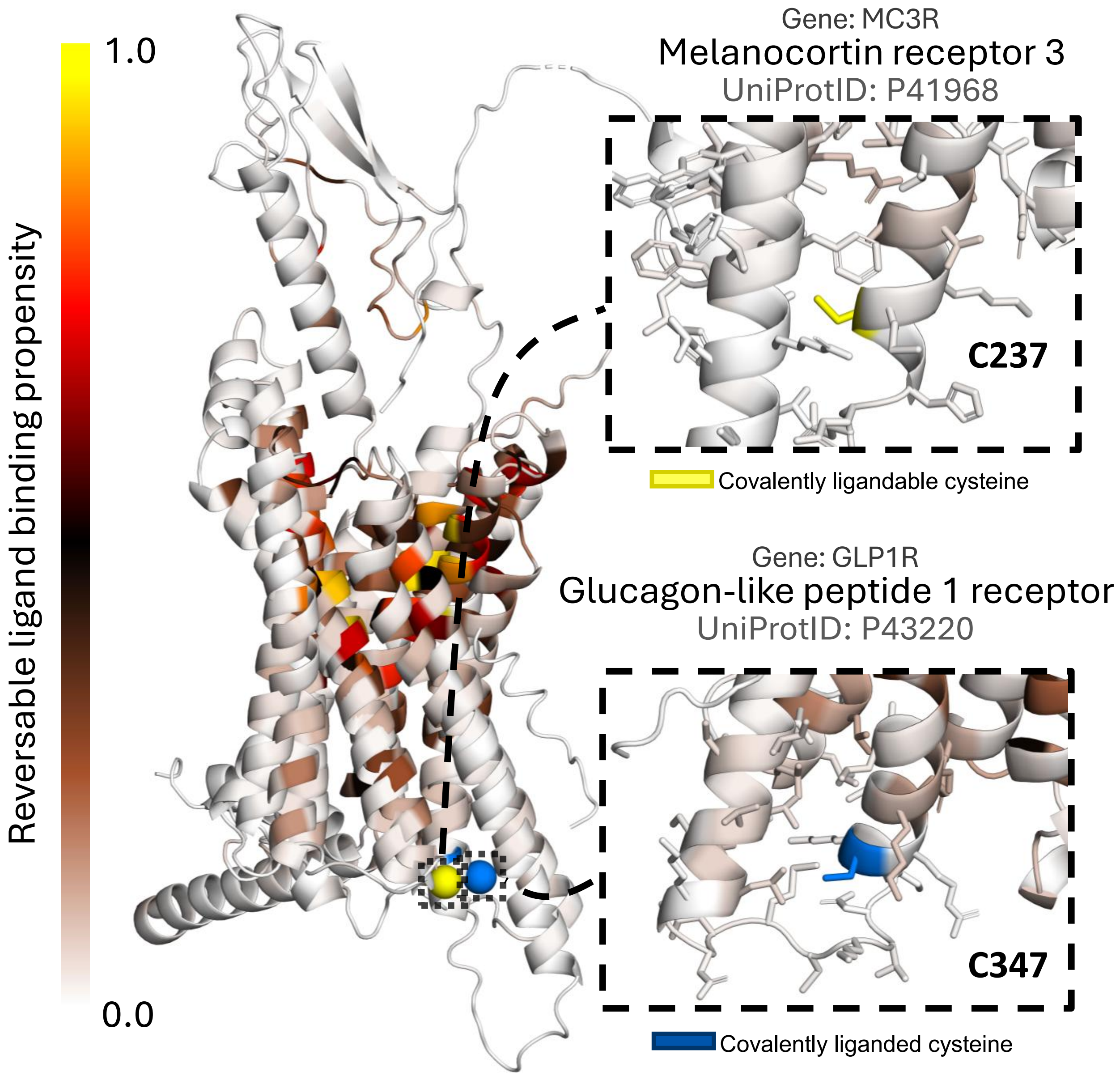

Figure S11: Reversible binding propensity scores for MC3R and GLP-1R mapped onto the ESM3 predicted (inactive) structures. The LigCys-predicted ligandable cysteine (C237$^{6.30}$) in MC3R and the analogous C347$^{6.31}$) in GLP-1R are colored yellow and blue, respectively. There nearby residues display low propensities (light pink color) for reversible binding.

### Supplementary Tables

Table S1: **Literature sources and experimental conditions in LigCysABPP.**

| Publication | Probe chemistry | # frag. | Cell line(s) | Note | [probe] conc. | [frag] conc. |
|---|---|---|---|---|---|---|
| Backus et al., Nature 2016 | IA-alkyne (IAA) | 65 | MDA-MB-231 and Ramos | isoTOP-ABPP | $100\,\mu\mathrm{M}$ | $500\,\mu\mathrm{M}$ or $250\,\mu\mathrm{M}$ (f4) |
| Bar-Peled et al., Cell 2017 | IAA | 2 | H2122, H460, A549, H1975, H358, H1792, H2009, HEK293T | isoTOP-ABPP | $100\,\mu\mathrm{M}$ | $500\,\mu\mathrm{M}$ |
| Vinogradova et al., Cell 2020 | IA-desthiobiotin (IA-DTB) | 10 | Primary human T cells | TMT-ABPP and isoTOP-ABPP | $100\,\mu\mathrm{M}$ | $500\,\mu\mathrm{M}$ |
| Cao et al., Anal. Chem. 2021 | Iodophenyl IA | 7 | HEK293T; Jurkat T lymphocytes | Multiplexed CuAAC–Suzuki–Miyaura chemoproteomic profiling | $20\,\mathrm{mM}$ | $200\,\mu\mathrm{M}$ |
| Kuljanin et al., Nature Biotech 2021 | Desthiobiotin IA (DBIA) | 285 | HCT116, HEK293T, PaTu-8988T | SLC-ABPP | $500\,\mu\mathrm{M}$ | $25\,\mu\mathrm{M}$ |
| Yan et al., ChemBioChem 2021 | IAA | 1 | Jurkat T lymphocytes | SP3-FAIMS isoTOP-ABPP | $200\,\mu\mathrm{M}$ | $500\,\mu\mathrm{M}$ |
| Yang et al., J. Am. Chem. Soc. 2022 | – | 21 | Ramos cells and MDA-MB-231 | DIA-ABPP | $100\,\mu\mathrm{M}$ | $500\,\mu\mathrm{M}$ |

Continued on next page

| Publication | Probe chemistry | # frag. | Cell line(s) | Note | [probe] conc. | [frag] conc. |
|---|---|---|---|---|---|---|
| Tao et al., J. Am. Chem. Soc. 2022 | IA-DTB | 4 | HEK293T; primary human T cells | TMT-ABPP | $100\,\mu\mathrm{M}$ | $500\,\mu\mathrm{M}^{a}$ |
| Koo et al., Nat. Commun. 2023 | NAIA-5 and NAIA-4 | 1 | 231MFP, MDA-MB-231, HepG2 | NAIA-ABPP | $10\,\mu\mathrm{M}$ | $500\,\mu\mathrm{M}$ |
| Yan et al., JACS Au 2023 | IAA | 1 | Jurkat and primary T cells | Cys-Surf / cell-surface cysteine chemoproteomics | $2\,\mathrm{mM}$ | $50\,\mu\mathrm{M}$ |
| Burton et al., J. Am. Chem. Soc. 2023 | IAA | 3 | HEK293T, HCT-15 | sCIP | $1\,\mathrm{mM}$ | $500\,\mu\mathrm{M}$ |
| Njomen et al., Nat. Chem. 2024 | IA-DTB | 28 | 22Rv1, Ramos | isoTOP-ABPP | $10\,\mu\mathrm{M}$ | $20\,\mu\mathrm{M}$ |
| Burton et al., Commun. Chem. 2024 | IA-DTB, IAA | 4 | HEK293T | sCIP-TMT | $1\,\mathrm{mM}$ | $500\,\mu\mathrm{M}$ |
| Biggs et al., Nat. Commun. 2025 | IA-DTB | 80 | HEK293T and Jurkat | High-throughput label-free DIA chemoproteomics (HT-LFQ) | $25\,\mu\mathrm{M}$ | Appropriate concentration |
| Tian et al., Nat. Commun. 2025 | IPM | 70 | HEK293T | CuAAC chemoproteomic profiling | $100\,\mu\mathrm{M}$ | $5\,\mu\mathrm{M}$ or $500\,\mu\mathrm{M}$ |

[a] Fragment concentration not explicitly stated in Tao et al.; table lists the concentration used in Vinogradova et al. (Cell 2020) as referenced.

Table S2: Evaluation of consensus thresholds for ABPP data selection in LigCys model training[a]

| ABPP training data | | | | LC3D benchmark evaluation (%) | | | | |
|---|---|---|---|---|---|---|---|---|
| **Criteria** | **Pos** | **Neg** | **Prot** | **PRC** | **ROC** | **Prec** | **Rec** | **Top-1** |
| 1S–1R | 13430 | 55441 | 7051 | 54.9 | 68.2 | 41.7 | 59.1 | 40.4 |
| 1S–2R | 6078 | 16791 | 4216 | 70.8 | 81.5 | 45.3 | 85.9 | 57.4 |
| 1S–3R | 3396 | 8600 | 2829 | 77.2 | 86.3 | 48.8 | 87.9 | 66.4 |
| 1S–4R | 2094 | 5048 | 2004 | 78.5 | 86.5 | 51.6 | 87.7 | 69.1 |
| 1S–5R | 1418 | 3354 | 1495 | 80.6 | 87.7 | 56.5 | 89.7 | 72.6 |
| 2S–2R | 3467 | 6838 | 2617 | 77.8 | 84.8 | 52.8 | 87.0 | 67.7 |
| 2S–3R | 2198 | 4765 | 1883 | 78.1 | 85.9 | 55.8 | 87.7 | 68.2 |
| 2S–4R | 1450 | 3326 | 1404 | 79.8 | 87.2 | 58.8 | 86.3 | 72.2 |
| 2S–5R | 988 | 2258 | 1041 | 80.2 | 87.0 | 61.3 | 85.0 | 73.1 |
| 3S–3R | 1115 | 1920 | 997 | 79.5 | 87.0 | 58.9 | 87.4 | 71.3 |
| 3S–4R | 867 | 1583 | 837 | 80.7 | 88.6 | 55.6 | 89.5 | 72.2 |
| 3S–5R | 649 | 1244 | 669 | 81.4 | 88.1 | 62.9 | 86.6 | 74.0 |
| **4S–4R** | **384** | **642** | **389** | **81.8** | **87.6** | **63.1** | **87.7** | **75.8** |
| 4S–5R | 347 | 590 | 358 | 80.9 | 86.5 | 64.3 | 85.0 | 74.9 |
| 5S–5R | 142 | 236 | 150 | 80.4 | 86.1 | 55.3 | 86.1 | 73.1 |

[a] LigCys models trained with ABPP datasets derived from increasingly strict consensus criteria are evaluated on the external LC3D dataset. At a consensus threshold $n$S–$m$R, label assignment requires at least $m$ records from $n$ different sources (see main text). Each model is an ensemble of 200 models (see Supplementary Methods). Performance is measured on the orthogonal LC3D benchmark set on a per-protein basis. Cysteines without covalent ligation are provisionally labeled neg. Precision and recall for predicting ligandable cysteines are calculated based on the decision rule used throughout this work, i.e., any cysteine with a nonzero LigCys score is predicted to be ligandable. Top-1 recovery refers to the probability that the cysteine with the highest LigCys score is a true pos given that the protein has at least one pos cysteine.

Table S3: Performance metrics for iterative training data expansion guided by LC3D-based evaluation (top block) and ABPP-based cross validation (bottom block).

| ABPP training data[a] | | | | ABPP valid. (%)[b] | | LC3D benchmark [c] (%) | | | | |
|---|---|---|---|---|---|---|---|---|---|---|
| **LC3D-guided expansion** | | | | | | | | | | |
| **Iter** | **Pos** | **Neg** | **Prot** | **PRCE** | **ROC** | **PRC** | **ROC** | **Prec** | **Rec** | **Top-1** |
| 0 | 354 | 574 | 366 | 74 | 74.6 | 79.0 | 85.6 | 62.9 | 85.4 | 70.9 |
| 1 | 359 | 649 | 442 | 92 | 78.2 | 79.7 | 85.6 | 64.5 | 84.8 | 73.1 |
| 2 | 369 | 739 | 534 | 93 | 75.9 | 81.5 | 87.6 | 65.2 | 86.1 | 75.3 |
| 3 | 384 | 874 | 674 | 91 | 74.2 | 83.6 | 89.2 | 60.7 | 88.8 | 77.1 |
| 4 | 401 | 1032 | 827 | 116 | 77.5 | 83.8 | 88.6 | 64.3 | 88.3 | 78.5 |
| 5 | 425 | 1186 | 966 | 139 | 79.3 | 83.3 | 88.2 | 65.2 | 87.4 | 78.0 |
| **6** | **441** | **1345** | **1099** | **141** | **78.2** | **84.9** | **89.0** | **67.6** | **87.9** | **80.7** |
| 6-SA[f] | 441 | 1345 | 1099 | 128 | 76.7 | 85.3 | 90.3 | 61.6 | 91.9 | 79.8 |
| **ABPP-guided expansion** | | | | | | | | | | |
| **Tour[d]** | **Pos** | **Neg** | **Prot** | **PRCE** | **ROC** | **PRC** | **ROC** | **Prec** | **Rec** | **Top-1** |
| 0 | 354 | 574 | 366 | 74 | 74.6 | 79.0 | 85.6 | 62.9 | 85.4 | 70.9 |
| 1 | 378 | 643 | 452 | 93 | 78.8 | 80.1 | 86.5 | 63.4 | 85.4 | 73.5 |
| 2 | 468 | 967 | 817 | 122 | 82.1 | 81.7 | 88.1 | 63.8 | 87.4 | 74.9 |
| 3 | 555 | 1432 | 1258 | 147 | 82.5 | 81.3 | 88.3 | 60.2 | 89.2 | 73.5 |
| **4** | **645** | **1998** | **1744** | **187** | **83.5** | **80.4** | **88.2** | **60.1** | **88.8** | **71.3** |
| 4-SA[f] | 645 | 1998 | 1744 | 178 | 80.9 | 79.8 | 87.2 | 59.7 | 89.5 | 70.0 |
| **Blended[e]** | - | - | - | - | - | 84.0 | 88.9 | 59.2 | 90.1 | 77.6 |

[a] In the LC3D-guided data expansion, Top-1 recovery of LC3D liganded cysteines was used to select candidate data batches (complete data in Supplemental Data). The total number of pos, neg cysteines, and distinct proteins in the training set is given after each iteration. The start of the expansion (Iter 0) used the 4S–4R$^t$ dataset, which excludes 23 proteins quantified by 10 sources from the original 4S–4R dataset. [b] Model validation metrics are AUROC (ROC) and AUPRC enrichment (PRCE), defined as $(\text{AUPRC} - \text{random})/\text{random}$. Here random represents the AUPRC of a random classifier (% positive samples in the dataset). [c] For evaluation on the LC3D benchmark set, precision (Prec) and recall (Rec) were calculated based on the decision rule used throughout this work, i.e., any cysteine with a nonzero LigCys score is predicted to be ligandable. Top-1 was calculated using LC3D ligandability labels as defined in Supplementary Methods. [d] In the ABPP-guided data expansion, AUPRC enrichment (PRCE) was used to select candidate data batches. Tour denotes the number of successive tournaments (effectively iterations 1, 5, 9, and 13; see Supplemental Data). All other details follow the notes above. [e] Blended denotes ensemble prediction using both Iter 6 and Tour 4 models, combining the LC3D-guided model selected for orthogonal generalization with the ABPP-guided model selected for improved PRCE on ABPP validation data. [f] Structure-aware variant (see Supp. Methods, section 4.2).

Table S4: Features used for training the structure-aware LigCys–SA task head[a]

| Feature name | Definition |
| --- | --- |
| **Sequence-structure alignment** | |
| pKa_shift | KaML residue $pK_a$ deviation |
| norm_posn | Normalized sequence position along chain (0–1) |
| plddt | Per-residue confidence score (0–100) |
| **Surface topology** | |
| surf_patch_area | Mesh area in 5 Å geodesic patch |
| surf_patch_concave_frac | Fraction of patch vertices with negative curvature |
| **Cysteine exposome** | |
| d_sg_exposure_depth | Offset of SG atom to nearest surface vertex |
| dir_sg_dist | Ray-traced outward SG distance |
| mean_hydrophobicity_8A | Mean Kyte–Doolittle scale within 8 Å |
| density_net_charge_8A | Net formal charge per residue within 8 Å |
| bin1.density.class | Fraction of residues by class, 0–3 Å shell |
| bin2.density.class | Fraction of residues by class, 3–6 Å shell |
| bin3.density.class | Fraction of residues by class, 6–9 Å shell |
| **Binding proximity** | |
| bind_euc_std | Std. of Euclidean distances to binding residues |
| bind_geo_mean | Mean geodesic distance to binding residues |
| bind_count_6A | Number of binding residues within 6 Å |
| **KaML summaries** | |
| rad_abs_shift_mean_3A | Neighbor average $\|pK_a \text{ shift}\|$, 3 Å |
| radial_avg_norm_sasa_3A | Neighbor average normalized SASA, 3 Å |

*Continued on next page*

| Feature name | Definition |
|---|---|
| rad_abs_shift_mean_6A | Neighbor average $|\text{pK}_a$ shift$|$, 6 Å |
| **RIDA summaries** | |
| anchor_geo_min | Minimum geodesic distance to ANCHOR2 residues |
| anchor_geo_mean | Mean geodesic distance to ANCHOR2 residues |
| vsl2b_label | Binary indicator of VSL2B disorder region |
| vsl2b_euc_min | Minimum Euclidean distance to VSL2B residues |
| vsl2b_geo_min | Minimum geodesic distance to VSL2B residues |
| vsl2b_count_6A | Number of VSL2B residues within 6 Å |
| mdp_label | Binary indicator of MDP disorder region |
| mdp_euc_min | Minimum Euclidean distance to MDP residues |
| mdp_euc_med | Median Euclidean distance to MDP residues |
| mdp_euc_std | Std. of Euclidean distances to MDP residues |
| mdp_geo_min | Minimum geodesic distance to MDP residues |
| mdp_count_3A | Number of MDP residues within 3 Å |

[a]Features were computed from ESM3-generated structures (MSMS surfaces; freesasa SASA), pre-processed by clipping to finite bounds, $\log_{1p}$ transforms where appropriate, zero-imputation for missing values, and z-score normalization for correlation screening. Redundancy was reduced by removing near-zero-variance features and pruning one of each pair with $|r| > 0.9$. Class "density" features denote fractions of residue classes within 0–3 Å, 3–6 Å, and 6–9 Å shells. Units: distances in Å, pLDDT on 0–100, charge unitless, counts as integers, $\text{pK}_a$ shifts in standard chemical units.

Table S5: Performance of the sequence-only and structure-aware LigBind models in predicting reversible ligand-binding residues[a]

| Metrics (%) | LigBind | LigBind-SA |
|---|---|---|
| **AUROC** | 93.9 | 95.1 |
| **AUPRC** | 74.5 | 77.5 |
| **Top-5** | 79.2 | 81.2 |
| **Top-10** | 78.1 | 80.1 |
| **Top-20** | 76.6 | 79.5 |

[a] The sequence-only LigBind and structure-aware LigBind-SA models were trained and evaluated using LatentLift-derived cluster labels (see Supplementary Methods). The ESM3 predicted structures were used in training and testing the LigBind-SA model. Performance metrics reflect an ensemble of 200 models. Per-residue ligandability probabilities were averaged across all 200 models. All metrics were computed on a per-protein basis using the held-out test set of 100 proteins. Test-split POS(%) (site-level prevalence; random AUPRC baseline) = 7.2%.

Table S6: Data for training LigBind and functional context models[a]

| Dataset | #UIDs | #Sites (UID–ROI) | #Clusters (repr.) | #POS | POS (%) |
|---|---|---|---|---|---|
| LigBind3D | 1998 | 687712 | 581493 | 45617 | 6.6 |
| ZNBind3D | 3209 | 960709 | 508408 | 20280 | 2.1 |
| CUBind3D | 896 | 283957 | 96201 | 6701 | 2.4 |
| FEBind3D | 1089 | 352546 | 156089 | 5488 | 1.6 |
| FESBind3D | 1167 | 449955 | 180538 | 6662 | 1.5 |
| HEMBind3D | 2467 | 762263 | 192486 | 5350 | 0.7 |
| SSBind3D | 83159 | 417152 | 137010 | 91812 | 22.0 |

[a]Summary of dataset size, LatentLift clustering, and positive-site prevalence for LigBind3D and the functional-context datasets. #UIDs denotes unique protein identifiers; #Sites denotes unique UID–ROI pairs; #Clusters denotes LatentLift cluster representatives; and #POS and POS (%) denote positive UID–ROI pairs after cluster-level label reconciliation. Clustering, label reconciliation, and leakage-free partitioning are described in the Supplementary Methods.

Table S7: Summary of hyperparameters that differ among the various AiPP models[a]

| Model | Loss | $\alpha$ | $\gamma$ | Batch Neg:Pos |
|---|---|---|---|---|
| LigCys | binary focal (fixed) | 0.66 | 1.0 | 10:1 |
| LigCys-SA | binary focal (fixed) | 0.66 | 1.0 | 10:1 |
| LigBind | adaptive binary focal | 0.80 | init 1.0 | 15:1 |
| LigBind-SA | adaptive binary focal | 0.80 | init 1.0 | 15:1 |
| ZNBind | adaptive binary focal | 0.80 | init 2.0 | 49:1 |
| CUBind | adaptive binary focal | 0.90 | init 2.0 | 49:1 |
| FEBind | adaptive binary focal | 0.90 | init 2.0 | 99:1 |
| FESBind | adaptive binary focal | 0.90 | init 2.0 | 99:1 |
| HEMBind | adaptive binary focal | 0.90 | init 2.0 | 199:1 |
| SSBind | adaptive binary focal | 0.80 | init 2.0 | 15:1 |

[a] LigCys models use fixed binary focal loss with the shown $\alpha$ and $\gamma$. LigBind and all functional-context models (ZNBind, CUBind, FEBind, FESBind, HEMBind, SSBind) use adaptive binary focal loss with 20 confidence bins; $\gamma$ is initialized as shown, updated every 5 epochs, clamped to [0.5, 5.0], and includes a calibration penalty term ($\lambda = 0.1$). "Batch Neg:Pos" denotes the negative-to-positive sampling ratio used when constructing training batches (not the dataset class prevalence); for functional context tasks, this ratio is set based on label prevalence in the corresponding training data. Functional context models are sequence-only and have no structure-aware variants. All other shared training settings are described in Supplemental Methods.

Table S8: Performance of sequence-only functional-context models (Cys-only)[a]

| Target | POS(%) | AUROC | AUPRC | Top-1 |
|---|---|---|---|---|
| **Zn** | 55.6 | 93.6 | 95.3 | 96 |
| **Cu** | 22.2 | 97.1 | 96.0 | 100 |
| **Fe** | 11.7 | 81.7 | 85.1 | 87 |
| **Fe–S** | 59.9 | 94.4 | 93.6 | 93 |
| **Heme** | 2.1 | 99.5 | 96.6 | 93 |
| **Disulfide** | 40.7 | 88.7 | 89.4 | 83 |

[a] Sequence-only functional-context annotation models (trained using LatentLift; see Supplementary Methods). Metrics are reported as percentages and computed on a per-protein basis on held-out test sets. Although trained on all residues, performance is evaluated on cysteine positions only (matching downstream application to LigCys); POS(%) denotes cysteine-site prevalence in the test split. Top-1 denotes the fraction of proteins for which the highest-scoring cysteine is labeled positive. Test-set sizes (proteins): Zn ($n$=277), Cu ($n$=69), Fe ($n$=91), Fe–S ($n$=116), Heme ($n$=200), S–S ($n$=1009).

### References

(1) Backus, K. M.; Correia, B. E.; Lum, K. M.; Forli, S.; Horning, B. D.; González-Páez, G. E.; Chatterjee, S.; Lanning, B. R.; Teijaro, J. R.; Olson, A. J.; Wolan, D. W.; Cravatt, B. F. Proteome-Wide Covalent Ligand Discovery in Native Biological Systems. *Nature* **2016**, *534*, 570–574.

(2) Vinogradova, E. V. et al. An Activity-Guided Map of Electrophile-Cysteine Interactions in Primary Human T Cells. *Cell* **2020**, *182*, 1009–1026.e29.

(3) Cao, J.; Boatner, L. M.; Desai, H. S.; Burton, N. R.; Armenta, E.; Chan, N. J.; Castellón, J. O.; Backus, K. M. Multiplexed CuAAC Suzuki–Miyaura Labeling for Tandem Activity-Based Chemoproteomic Profiling. *Anal. Chem.* **2021**, *93*, 2610–2618.

(4) Kuljanin, M.; Mitchell, D. C.; Schweppe, D. K.; Gikandi, A. S.; Nusinow, D. P.; Bulloch, N. J.; Vinogradova, E. V.; Wilson, D. L.; Kool, E. T.; Mancias, J. D.; Cravatt, B. F.; Gygi, S. P. Reimagining High-Throughput Profiling of Reactive Cysteines for Cell-Based Screening of Large Electrophile Libraries. *Nat. Biotechnol.* **2021**, *39*, 630–641.

(5) Yan, T.; Desai, H. S.; Boatner, L. M.; Yen, S. L.; Cao, J.; Palafox, M. F.; Jami-Alahmadi, Y.; Backus, K. M. SP3-FAIMS Chemoproteomics for High-Coverage Profiling of the Human Cysteinome**. *ChemBioChem* **2021**, *22*, 1841–1851.

(6) Yang, F.; Jia, G.; Guo, J.; Liu, Y.; Wang, C. Quantitative Chemoproteomic Profiling with Data-Independent Acquisition-Based Mass Spectrometry. *J. Am. Chem. Soc.* **2022**, *144*, 901–911.

(7) Tao, Y.; Remillard, D.; Vinogradova, E. V.; Yokoyama, M.; Banchenko, S.; Schwefel, D.; Melillo, B.; Schreiber, S. L.; Zhang, X.; Cravatt, B. F. Targeted Protein Degra-

dation by Electrophilic PROTACs That Stereoselectively and Site-Specifically Engage DCAF1. *J. Am. Chem. Soc.* **2022**, *144*, 18688–18699.

(8) Koo, T.-Y.; Lai, H.; Nomura, D. K.; Chung, C. Y.-S. N-Acryloylindole-alkyne (NAIA) Enables Imaging and Profiling New Ligandable Cysteines and Oxidized Thiols by Chemoproteomics. *Nat. Commun.* **2023**, *14*, 3564.

(9) Yan, T.; Boatner, L. M.; Cui, L.; Tontonoz, P. J.; Backus, K. M. Defining the Cell Surface Cysteinome Using Two-Step Enrichment Proteomics. *JACS Au* **2023**, *3*, 3506–3523.

(10) Njomen, E.; Hayward, R. E.; DeMeester, K. E.; Ogasawara, D.; Dix, M. M.; Nguyen, T.; Ashby, P.; Simon, G. M.; Schreiber, S. L.; Melillo, B.; Cravatt, B. F. Multi-Tiered Chemical Proteomic Maps of Tryptoline Acrylamide–Protein Interactions in Cancer Cells. *Nat. Chem.* **2024**, *16*, 1592–1604.

(11) Biggs, G. S. et al. Robust Proteome Profiling of Cysteine-Reactive Fragments Using Label-Free Chemoproteomics. *Nat. Commun.* **2025**, *16*, 73.

(12) Tian, C.; Sun, L.; Liu, K.; Fu, L.; Zhang, Y.; Chen, W.; He, F.; Yang, J. Proteome-Wide Ligandability Maps of Drugs with Diverse Cysteine-Reactive Chemotypes. *Nat. Commun.* **2025**, *16*, 4863.

(13) Bar-Peled, L. et al. Chemical Proteomics Identifies Druggable Vulnerabilities in a Genetically Defined Cancer. *Cell* **2017**, *171*, 696–709.e23.

(14) Burton, N. R.; Polasky, D. A.; Shikwana, F.; Ofori, S.; Yan, T.; Geiszler, D. J.; Veiga Leprevost, F. D.; Nesvizhskii, A. I.; Backus, K. M. Solid-Phase Compatible Silane-Based Cleavable Linker Enables Custom Isobaric Quantitative Chemoproteomics. *J. Am. Chem. Soc.* **2023**, *145*, 21303–21318.

(15) Burton, N. R.; Backus, K. M. Functionalizing Tandem Mass Tags for Streamlining Click-Based Quantitative Chemoproteomics. *Commun. Chem.* **2024**, *7*, 80.

(16) Liu, R.; Clayton, J.; Shen, M.; Bhatnagar, S.; Shen, J. Machine Learning Models to Interrogate Proteome-Wide Covalent Ligandabilities Directed at Cysteines. *JACS Au* **2024**, *4*, 1374–1384.

(17) Zhang, C.; Zhang, X.; Freddolino, L.; Zhang, Y. BioLiP2: An Updated Structure Database for Biologically Relevant Ligand–Protein Interactions. *Nucl. Acids Res.* **2024**, *52*, D404–D412.

(18) ESM Team. ESM Cambrian: Revealing the Mysteries of Proteins with Unsupervised Learning. https://www.evolutionaryscale.ai/blog/esm-cambrian.

(19) Fu, L.; Niu, B.; Zhu, Z.; Wu, S.; Li, W. CD-HIT: Accelerated for Clustering the next-Generation Sequencing Data. *Bioinformatics* **2012**, *28*, 3150–3152.

(20) Takahashi, M. et al. DrugMap: A Quantitative Pan-Cancer Analysis of Cysteine Ligandability. *Cell* **2024**, *187*, 2536–2556.e30.

(21) Hayes, T. et al. Simulating 500 Million Years of Evolution with a Language Model. *Science* **2025**, *387*, 850–858.

(22) Sanner, M. F.; Olson, A. J.; Spehner, J.-C. Reduced Surface: An Efficient Way to Compute Molecular Surfaces. *Biopolymers* **1996**, *38*, 305–320.

(23) Mitternacht, S. FreeSASA: An Open Source C Library for Solvent Accessible Surface Area Calculations. *F1000Res.* **2016**, *5*, 189.

(24) Krapp, L. F.; Abriata, L. A.; Cortés Rodriguez, F.; Dal Peraro, M. PeSTo: Parameter-Free Geometric Deep Learning for Accurate Prediction of Protein Binding Interfaces. *Nat. Commun.* **2023**, *14*, 2175.

(25) Benjamini, Y.; Hochberg, Y. Controlling the false discovery rate: a practical and powerful approach to multiple testing. *Journal of the Royal statistical society: series B (Methodological)* **1995**, *57*, 289–300.

(26) Dayhoff II, G. W.; Uversky, V. N. Rapid Prediction and Analysis of Protein Intrinsic Disorder. *Protein Sci.* **2022**, *31*, e4496.

(27) Mészáros, B.; Erdős, G.; Dosztányi, Z. IUPred2A: Context-Dependent Prediction of Protein Disorder as a Function of Redox State and Protein Binding. *Nucl. Acids Res.* **2018**, *46*, W329–W337.

(28) Uhlen, M.; Oksvold, P.; Fagerberg, L.; Lundberg, E.; Jonasson, K.; Forsberg, M.; Zwahlen, M.; Kampf, C.; Wester, K.; Hober, S.; Wernerus, H.; Björling, L.; Ponten, F. Towards a Knowledge-Based Human Protein Atlas. *Nat. Biotechnol.* **2010**, *28*, 1248–1250.

(29) Rice, P.; Longden, I.; Bleasby, A. EMBOSS: the European molecular biology open software suite. *Trends in genetics* **2000**, *16*, 276–277.

(30) Qu, Z. et al. Discovery of a First-in-Class Covalent Allosteric SHP1 Inhibitor with Immunotherapeutic Activity. *Angew. Chem. Int. Ed.* **2026**, *65*, e25126.
